# Activation-Free Upgrading of Carboxylic Acids to Aldehydes and Alcohols

**DOI:** 10.1101/2025.07.28.667276

**Authors:** William B. Black, Samer Saleh, Sean Perea, Emma Luu, Youtian Cui, Jiasi Sun, Zhaoxi Wang, Shyanne Lambrecht, Salman Awachi, Declan Hayworth, Andrea Wang, Chloe Chuayiuso, Raine Hagerty, Patrick C. Gilcrease, Feng Jiao, Zhen He, Justin B. Siegel, Han Li

## Abstract

Advances in organic and gas waste valorization have enabled high-yield production of carboxylic acids, positioning them as promising feedstocks for the bioeconomy. However, carboxylic acids must be activated before downstream use, typically requiring ATP, CoA, or reduced ferredoxin to overcome unfavorable thermodynamics. These activators are costly to generate and divert carboxylic acids into CO_2_-releasing pathways, reducing carbon efficiency. Here, we demonstrate that aldehyde dehydrogenases (ALDHs) can directly reduce carboxylic acids to aldehydes without prior activation, a process previously thought to be biologically inaccessible. Screening 133 ALDHs revealed that this activity is remarkably widespread within the protein family, enabling production of aliphatic aldehydes and alcohols, diols, and aromatic alcohols, at titers >1 g/L, in some cases, after optimization of thermodynamic driving forces. Additionally, we applied this system to upgrade waste-derived carboxylic acid effluent streams from wastewater sludge, food waste, and waste gas (CO_2_). This activation-free process, termed “reverse aldehyde oxidation” (rAOX), establishes a broadly applicable, energy-efficient platform for carboxylic acid valorization at 100% carbon yield. Analogous to the reverse tricarboxylic acid cycle (rTCA) and reverse β-oxidation (rBOX), rAOX exemplifies that metabolic reactions classically defined as unidirectional may have unexpected plasticity to operate in reverse and open new avenues in biomanufacturing.

## Introduction

Landmark work on discovering and engineering new metabolic pathways has redefined the traditional doctrine that metabolic pathways are constrained by inherent thermodynamic directionality. The discovery of the reverse tricarboxylic acid (rTCA) carbon fixation cycle challenged the long-standing assumption that the TCA cycle is thermodynamically feasible only in the oxidative, CO_2_-releasing direction^1–5^. The engineered reverse β-oxidation cycle (rBOX) demonstrated that fatty acid degradation, long believed to be unidirectionally favored in the catabolic direction, can not only operate in the anabolic direction, but it is also more efficient in doing so than the natural fatty acid synthesis pathways by omitting ATP activation of extender units^6, 7^.

In this work, we report that oxidative degradation of aldehydes to carboxylic acids using NAD^+^, the metabolic reaction utilized universally by divergent organisms to detoxify aldehydes, can proceed in the reductive direction using NADH, which is against the long-established belief that this reaction is thermodynamically irreversible^8,9^. Before this work, natural and engineered metabolic pathways that activate carboxylic acids to produce aldehydes and alcohols require ATP^10–14^, CoA-activated intermediates^10,11^, or strong reducing equivalent reduced ferredoxins^14,15^. Here, we show that the reverse aldehyde oxidation (rAOX), demonstrated as a cell-free system, operates without these high-energy metabolic currencies, and rAOX can accept a wide range of carboxylic acids to upgrade them into aliphatic and aromatic alcohols and diols, as well as aldehydes, including reaching >1 g/L titers of propanol, butanol, and isobutanol. These compounds have applications as fuels, commodities, and specialty chemicals. We also showed that rAOX can directly process carboxylic acid feedstocks in effluent streams of food waste, wastewater, and waste gas (CO_2_) treatments.

The problem we tackle here, namely carbon-efficient upgrading of carboxylic acids, represents a critical need in bioeconomy. Organic acids are common products in microbial fermentation of biomass-derived sugars^16^. Innovations in pathway design, such as the construction of non-oxidative glycolysis (NOG)^17,18^, have maximized the carbon yield of such processes. Recent technologies valorizing cheap, carbon-laden waste streams also largely result in carboxylic acids. These emerging technologies include arrested anaerobic digestion of municipal and agricultural wastes^19,20^, waste C1 gas fermentation^21^, and electrochemical carbon capture^22,23^. In particular, due to the large quantity of wet wastes available at a negligible cost (>50 million dry tons or 0.7 quadrillion British thermal units in energy content generated annually in the United States)^24^, upcycling wet wastes to carboxylic acids has the potential to provide truly economically scalable feedstocks to biorefineries^20^. However, biological utilization of carboxylic acids requires activation (as mentioned above), which ultimately limits the theoretical carbon yield. This is because to generate the activating reagent such as ATP, either a fraction of the carboxylic acid substrates need to be directed to CO_2_-releasing pathways, including TCA cycle^25^, or co-substrate feeding may be used^15^. Through eliminating the need for activation, the rAOX process reported here features a 100% theoretical carbon yield in upgrading carboxylic acids.

Activation of carboxylic acids also represents a key energy hurdle that shapes biology^26–28^. Producing carboxylic acids is spontaneous: Carboxylic acids are the thermodynamic minima of metabolites across a wide range of physiologically relevant pH and cofactor reduction potentials^26^, which is consistent with their accumulation as fermentation products from different feedstocks as mentioned above. Reacting with carboxylic acids is energy-intensive; reducing carboxylic acids to a more reactive aldehyde state is highly unfavorable^26^. In the Calvin cycle of CO_2_ fixation, 5 out of 7 ATPs (∼70%) invested to produce one pyruvate are exclusively used to power the reduction of carboxylic acid functional groups. In other known carbon fixation cycles, this number ranges from ∼50% to 100%^27^. Here, we demonstrate that ATP-free activation of carboxylic acids is feasible, which may motivate metabolic engineers to innovate outside the constraints of natural biology to create more energy-efficient pathways.

## Results

### Reversible activity of aldehyde dehydrogenases (ALDHs) is widespread in protein sequence space

Aldehyde dehydrogenases (ALDHs) naturally catalyze the oxidation of aldehydes to their respective carboxylic acids using NAD^+^ or NADP^+^ in a CoA-independent fashion. Since their initial characterization in the early 1900’s^29,30^, ALDHs have been characterized as “irreversible” enzymes^8,9^, which is consistent with the reduction of carboxylic acids using NADH being theorized to be greatly unfavorable based on standard Gibbs free energy calculation (discussed below). However, a smattering of publications have suggested that ALDHs may exhibit reversible activity, albeit very low, if tested outside of biologically accessible reaction conditions^31,32^. For example, Minteer and co-workers developed a fusion protein of an ALDH from *Bacillus cereus* and an alcohol dehydrogenase (ADH) from *Escherichia coli* to achieve direct reduction of acetate with an electrochemical redox cofactor cycling system^31^. Over 48 hours, 0.16 mM acetaldehyde and 1.09 mM ethanol were produced. Tommasi and co-workers demonstrated the immobilization of ALDH from *Saccharomyces cerevisiae* and ADH from *S. cerevisiae* onto mesoporous siliceous materials for the direct reduction of propionate, using high NADH loading (50 mM) to produce sub-micromolar concentrations of propionaldehyde and propanol^32^. Similar enzymatic cascades have been developed to produce methanol, where excess supply of NADH, aggressive cofactor recycling, and co-localization of enzymes were the keys to increase productivity^33^. Bioprospecting efforts for alternative formaldehyde dehydrogenases have yielded mixed reports of improved function^34–36^.

These previous efforts motivated our systematic search for enhanced reversible activity across the broad sequence space of the ALDH family (InterPro Pfam PF00171) (**Figure 1A**) and inspired our design of a screening platform comprised of an in vitro multi-enzyme cascade (**Figure 1B**). Where in the event that the candidate ALDH reduces a carboxylic acid molecule to an aldehyde, an alcohol dehydrogenase from *S. cerevisiae* (ScADH) immediately sequesters any aldehyde produced by reduction to an alcohol. Additionally, any oxidized cofactor (NAD^+^) is rapidly recycled back to NADH using phosphite dehydrogenase (PTDH). Alcohol production thus serves as the signal output for carboxylic acid reduction activity.

**Figure 1:**
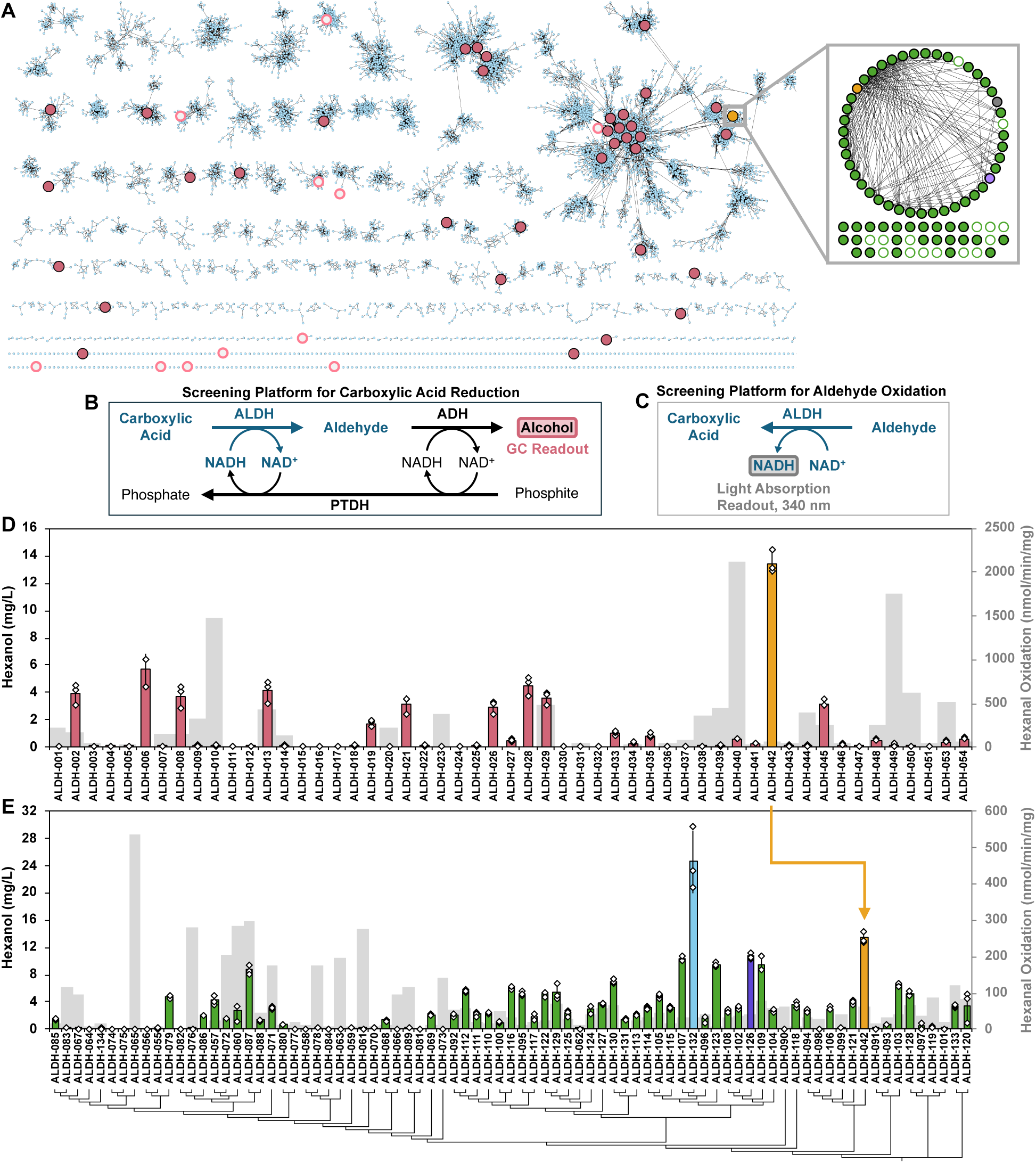
Screening Aldehyde Dehydrogenases for Carboxylic Acid Reduction Activity. **A)** Sequence Similarity Network visualizing the ALDH InterPro Pfam PF00171. Round 1 screening candidates shown in pink circles. Round 2 Screening Candidates shown in green circles. Yellow circle, GtALDH. Purple circle, ALDH-126. Blue circle, ALDH-132. Open circles are negative hits. **B)** Enzymatic cycling system for screening ALDHs for carboxylic acid reduction activity. If the ALDH can reduce the carboxylic acid to an aldehyde, the aldehyde is rapidly reduced to its respective alcohol using an alcohol dehydrogenase (ADH). Oxidized NADH (NAD^+^) is recycled back to NADH by phosphite dehydrogenase (PTDH). **C)** Screening platform for ALDH aldehyde oxidation activity. **D)** Round 1 ALDH candidate screening for hexanoate reduction (colored bars) and hexanal oxidation (grey bars). **E)** Round 2 ALDH candidate screening. Investigating the ALDH-042 (GtALDH, yellow bar) clade for improved carboxylic acid reduction activity. Candidates are arranged using a phylogenetic tree generated from their respective protein sequences. Values represent the average of three biological replicates, unless stated otherwise. ALDH-050 and ALDH-051 are an average of two biological replicates. Error bars represent one standard deviation.

Using the bioinformatics tool EFI-EST, we mapped the ∼500,000 sequences within the ALDH family, yielding a visual sequence similarly network of distinct sequence clusters. In this network, nodes are comprised of enzyme sequences with high similarity, and these nodes are connected by edges to group the nodes into subnetworks (**Figure 1A**). In the first round, we selected 53 representative sequences (indicated with pink and yellow circles on the Network (**Figure 1A**)) from different subnetworks to survey a broad diversity of sequence space. Additionally, we selected sequences which were well characterized in literature and sequences which were easily isolatable from genomic DNA already present in our laboratory, totaling 53 candidate ALDH sequences in Round 1 of screening. These ALDH candidates were tested for carboxylic acid reduction activity with hexanoate (rAOX-desired direction) (**Figure 1B**) and for aldehyde oxidation activity with hexanal (natural direction) (**Figure 1C**). Remarkably, 39 of the 53 initial ALDHs screened successfully reduced hexanoate (**Figure 1D, Supplementary Table 1**). 5 of the 14 candidates which did not show rAOX activity also did not exhibit hexanal oxidation activity with NAD^+^, and we categorized them to be non-active with this substrate or NAD(H) (**Supplementary Table 1**). Excluding these non-active candidates, a remarkable 81% of the initial candidates screened exhibited hexanoate reduction capabilities. Control reactions with individual reaction component drop-outs indicated the ALDH candidates were specifically enabling this conversion (**Supplementary Figure 1**). One candidate from Round 1 screening, ALDH-042 from *Geobacillus thermodenitrificans* (GtALDH) (**Figure 1A, yellow circle; Figure 1D, yellow bar**), stood out as an exceptional carboxylic acid reduction candidate, producing 13.45 mg/L of hexanol.

### Identifying a high sequence-similarity group enriched with ALDH reversible activity

Next, we performed a second round of ALDH bio-prospecting surrounding GtALDH. In Round 2 of screening, we selected an additional 80 ALDH candidates. Among them, 45 candidates encompassed all ALDH sequences with >90% sequence homology to GtALDH in the UniRef database at the time of screening and 35 additional candidates between 55-89% sequence homology (the majority being >70% similar) to GtALDH. These Round 2 ALDH candidates are shown with green circles on the Network (**Figure 1A**). In this screen, nearly all candidates with >90% sequence similarity to GtALDH exhibited hexanoate reduction activity, where only two candidates exhibited “not detected” activity, 6 exhibited trace activity (<5% of GtALDH at detectable levels), 36 exhibited moderate activity (>5% of GtALDH, <100% of GtALDH), and 1 candidate (ALDH-132) exhibited greater activity than GtALDH (**Figure 1E, Supplementary Table 2**). Candidates with lower than 90% sequence similarity to GtALDH performed notably worse, where 13 candidates exhibited “not detected” activity, 10 exhibited trace activity (<5% of GtALDH at detectable levels), and 12 exhibited moderate activity (>5% of GtALDH, <100% of GtALDH). Only one candidate did not show activity in the hexanal oxidation direction. This distinct density of ALDH enzymes with rAOX capabilities concentrating in the GtALDH clade may warrant future studies to understand the sequence-function relationship. The highest performing candidate, ALDH-132, and second highest candidate, ALDH-126, were selected for further characterization alongside GtALDH.

### Optimizing rAOX platform and expanding its alcohol product scope

Varying factors, such as carboxylic acid species, enzyme, substrate, and cofactor concentrations, and pH, revealed the different degrees of sensitivity of the system performance to these parameters **(Supplementary Figure 2A-D)**. With the optimized conditions (10 g/L ALDH, 1.5 g/L PTDH, 1-5 g/L ADH, and 3 mM NADH), we tested GtALDH, ALDH-126, and ALDH-132 with various carboxylic acids. For reactions with aliphatic acid substrates, acetate, propionate, and butyrate were supplied at 750 mM. Hexanoate was supplied at 50 mM. All systems showed robust ability to reduce aliphatic substrates, with ALDH-126 trending slightly higher than the other two candidates in most cases, producing 485 mg/L ethanol, 1757 mg/L propanol, 1848 mg/L butanol, and 375 mg/L hexanol (**Figure 2A-C, 2F**).

**Figure 2:**
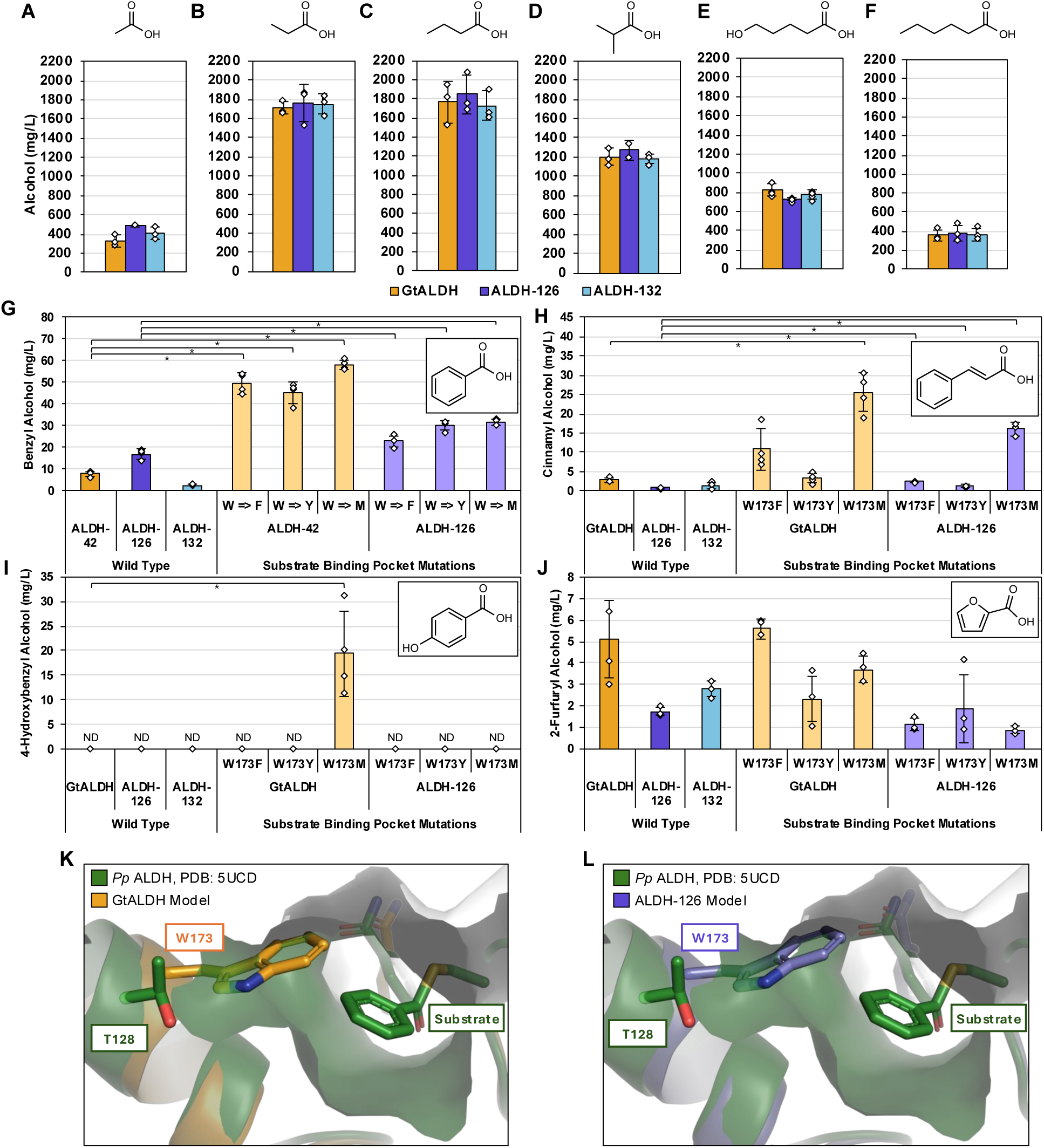
Expanding rAOX with Top ALDH Candidates. The rAOX system with top hits from ALDH bioprospecting screens was tested with different carboxylic acid species under optimized reaction conditions. **A)** rAOX using acetate. **B)** rAOX using propionate. **C)** rAOX using butyrate. **D)** rAOX using isobutyrate. TcADH was used in place of ScADH for these reactions. **E)** rAOX using 5-hydroxypentanote. TcADH was used in place of ScADH for these reactions. **F)** rAOX using hexanoate. **G)** rAOX using benzoate. Substrate binding pocket mutations on GtALDH and ALDH-126 scaffolds increased system activity. **H)** rAOX using trans-cinnamate. Substrate binding pocket mutations on GtALDH and ALDH-126 increased system activity. **I)** rAOX using 4-hydroxybenzoate. Substrate binding pocket mutation on GtALDH enabled new activity with this substrate. **J)** rAOX using 2-furoate. Substrate binding pocket mutations did not improve activity with GtALDH or ALDH-126. **K,L)** Structural overlay of AlphaFold3 model of GtALDH **(K)** or ALDH-126 **(L)** and the crystal structure of a benzaldehyde-utilized ALDH from *Pseudomonas putida*, PDB: 5UCD. In 5UCD, a substate analog is covalently bound to the catalytic cystine. **A-J)** values represent the average of at least three biological replicates. Error bars represent one standard deviation. Two-tailed *t*-tests were used to determine statistical significance (\**P*<0.05). “ND”: not detected.

When the system was translated to the branched carboxylic acid isobutyrate, the GtALDH-ScADH mediated system exhibited poor alcohol titers (**Supplementary Figure 3**), despite GtALDH’s broad substrate scope^37^. ScADH was identified as the bottleneck^38^, which was replaced by a broad substrate range alcohol dehydrogenase from *Tupaia chinensis* (TcADH), which had not been characterized before. This change in alcohol dehydrogenase significantly improved isobutanol production in GtALDH-mediated isobutyrate reduction, relative to ScADH (**Supplementary Figure 3**). When paired with TcADH, GtALDH, ALDH-126, and ALDH-132 reached isobutanol titers of 1202 mg/L, 1275 mg/L, and 1180 mg/L, respectively, from 522 mM isobutyrate (**Figure 2D**). Leveraging the promiscuity of TcADH, we also achieved diol production from the corresponding hydroxy acid. 818 mg/L, 725 mg/L, and 770 mg/L of 1,5-pentanediol was produced with GtALDH, ALDH-126, and ALDH-132, respectively, from 750 mM 5-hydroxypentanoic acid (**Figure 2E**).

When rAOX was tested with aromatic substrates (benzoate, trans-cinnamate, 4-hydroxybenzoate, and 2-furoate), the system performance was notably low (**Figure 2G-2J**). Since GtALDH is capable of performing the oxidation of benzaldehyde, but additional functionalization or bulk on the phenolic ring substantially reduces its activity^37^, we theorized that enhanced ALDH acceptance of these bulky substates required engineering. We compared the substrate entry channel of an AlphaFold3 model of GtALDH and ALDH-126 with a crystal structure of the benzaldehyde-utilizing ALDH from *Pseudomonas putida*, *Pp* ALDH (PDB: 5UCD) (**Figure 2K and 2L**). We noted residue W173 in both GtALDH and ALDH-126 corresponds to a threonine in *Pp* ALDH at the equivalent position (**Figure 2K and 2L**), potentially opening up room for the entry of bulkier substrates. When W173 was mutated to conserved, but smaller residues M, F, and Y, aromatic carboxylic acid reduction activity was significantly improved. GtALDH W173M increased benzyl alcohol formation by ∼7-fold over GtALDH wild type, increasing benzyl alcohol formation from 8 mg/L to 58 mg/L (**Figure 2G**). These mutations translated well to ALDH-126, improving benzoate reduction with all mutants (**Figure 2G**). Similar improvements were shown with trans-cinnamate, where GtALDH W173M increased cinnamyl alcohol production from 3 mg/L to 26 mg/L (**Figure 2H**). Remarkably, the W173M mutation on GtALDH enabled reduction activity with 4-hydroxybenzoate, producing 20 mg/L 4-hydroxybenzyl alcohol, while GtALDH wild type exhibiting no detectable reduction activity (**Figure 2I**). Reactions with 2-furoate responded differently compared to the other substrates, with no mutants exhibiting activity increases relative to their wild type parents with this smaller, more hydrophilic substrate (**Figure 2J**), suggesting alternative engineering strategy is needed.

### Interrogating the thermodynamic driving forces to enable arresting of rAOX at aldehyde

Direct reduction of carboxylic acids with NADH faces a formidable thermodynamic challenge, a ΔG° = +44.1 kJ/mol at standard reaction conditions with isobutyrate^39^ (See Methods), for example. To more clearly understand the contribution of different driving forces that make rAOX feasible, we performed a more detailed analysis **(Figure 3A)**.

**Figure 3:**
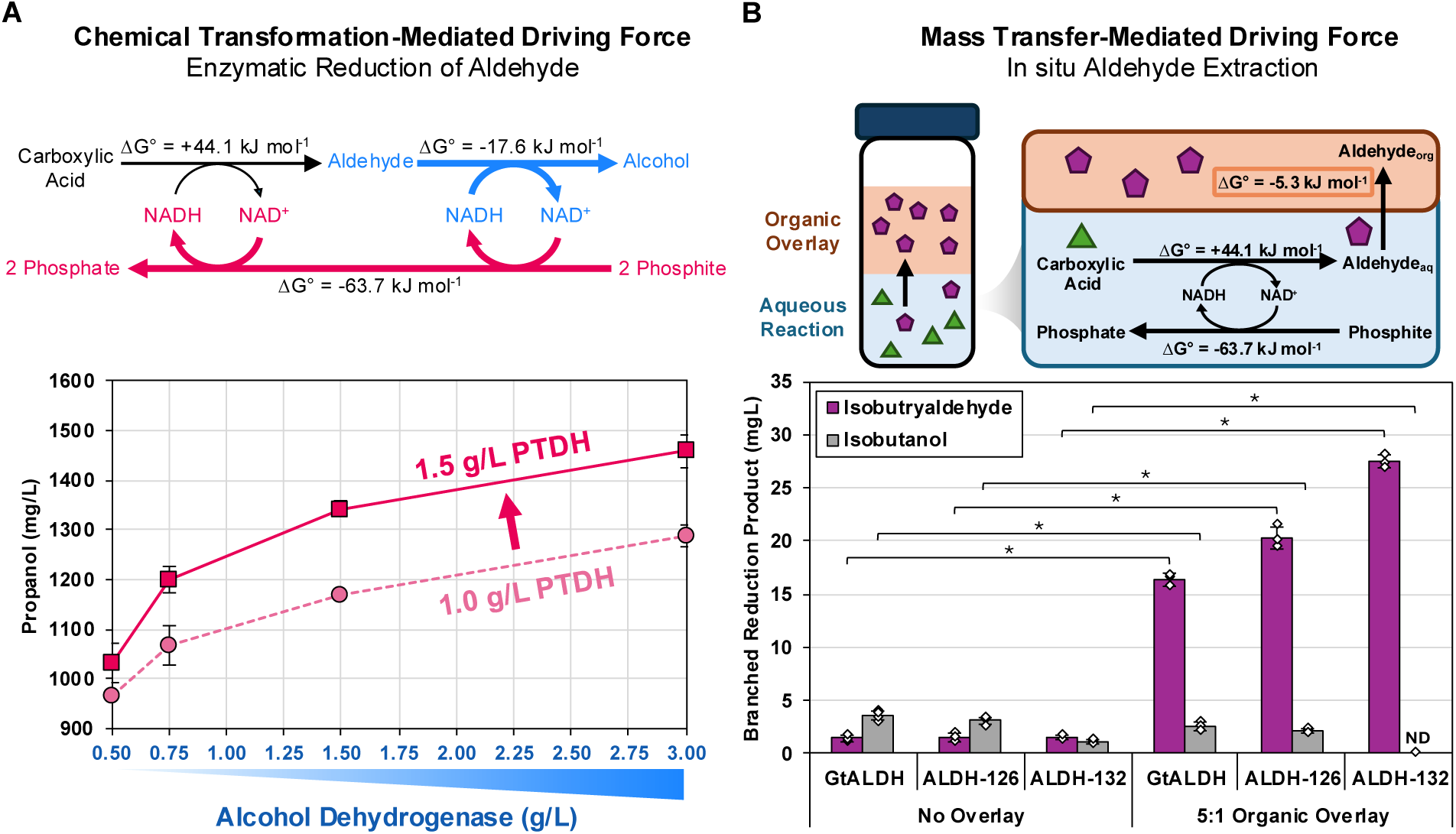
Optimized Driving Forces Facilitate Improved rAOX Activity. **A)** Chemical transformation-mediated driving forces improve reaction thermodynamics by sequestering aldehyde product away from the ALDH (blue) and recycling redox cofactor back to NADH (pink). Increasing the concentration of ADH and PTDH in GtALDH-mediated reduction of propionate improve propanol titers. GtALDH was added to the reactions at 5 g/L. **B)** Mass transfer-mediated driving forces improve reaction thermodynamics by diffusion of aldehyde out of the aqueous reaction mixture. In this application, an organic solvent overlay of ethoxybenzene was applied to continuously extract isobutyraldehyde in situ. Mass transfer-mediated driving forces preserved aldehyde as the terminal product. When organic overlay was added to reactions, five volumes of ethoxybenzene were added to the reaction vessel for every one volume of aqueous reaction volume. Organic overlay was added at t = 0. Values represent the average of at least three biological replicates. Error bars represent one standard deviation. Two-tailed *t*-tests were used to determine statistical significance (\**P*<0.05). “ND”: not detected. *

At standard reaction conditions, the ADH-mediated reduction of isobutyraldehyde to isobutanol yields a ΔG° = -17.6 kJ/mol (see Methods). Addition of NADH regeneration via PTDH yields an additional ΔG° = -63.7 kJ/mol in thermodynamic driving force (see Methods). Pairing these two driving forces with an ALDH capable of facilitating the direct reduction of carboxylic acids resulted in a net negative ΔG° for the overall reaction. The standard Gibbs free energy calculation here has limitations, such as not reflecting real reaction conditions and chemical concentrations, as well as not taking into account the kinetics of enzymes, both nontrivial. Nonetheless, it provides a rough indication of reaction favorability.

Consistent with these calculations, increasing PTDH or ADH concentrations resulted in significant improvement of rAOX performance **(Figure 3A)**. Only a modest increase of PTDH concentration from 1 to 1.5 g/L resulted in ∼15% increase in propanol production from propionate mediated by GtALDH. When ScADH concentration increased across a 6-fold gradient from 0.5 to 3 g/L, propanol titers increased ∼50%. While GtALDH likely remains the limiting enzyme in this system (**Supplementary Figure 4**), we hypothesize that increasing ScADH and PTDH concentrations minimized NAD^+^ and aldehyde build-up, maintaining driving force for carboxylic acid reduction.

Our experimental results and calculations above both suggest that the thermodynamic contribution of PTDH is greater than ADH. In fact, even without ADH, the overall reaction consisted of GtALDH and PTDH is predicted to already have a net-negative ΔG° **(Figure 3B)**. Indeed, when ADH is omitted from the carboxylic acid reduction system containing isobutyrate (**Figure 3B**), 1.5 mg/L, 1.5 mg/L, and 1.6 mg/L of isobutyraldehyde was produced using GtALDH, ALDH-126, or ALDH-132, respectively. Interestingly, these reactions also produced trace amounts of isobutanol despite the lack of ADH in the reaction, 3.5 mg/L, 3.1 mg/L, or 1.1 mg/L of isobutanol, respectively (**Figure 3B**). This is likely due to trace aldehyde reduction capabilities characterized in the ALDH family^40^.

To enhance the specific production of aldehydes, we leveraged a mass transfer-mediated driving force using an organic solvent to sequester isobutyraldehyde from the aqueous reaction phase to the organic phase during the reaction, which is predicted to contribute a ΔG° = -5.3 kJ/mol (**Figure 3B**). When we applied ethoxybenzene as an overlay solvent^41^ to ALDH-PTDH coupled reactions at a 5:1 solvent: aqueous ratio, the isobutyraldehyde production increased to 16.4 mg/L, 20.4 mg/L, and 27.5 mg/L using GtALDH, ALDH-126, and ALDH-132, respectively (**Figure 3B**). With the in-situ overlay solvent extraction, non-specific isobutanol production was decreased in all samples, relative to the “No Overlay” samples (**Figure 3B**). Remarkably, reactions with ALDH-132 achieved 100% aldehyde purity, with no isobutanol detected in the reaction. Continuous exchange of the organic layer or selection of an alternative bio-compatible solvent with improved analyte partitioning coefficient may further improve productivity. Other processes to rapidly separate and recover aldehydes, such as ones that leverage their high vapor pressures^42,43^, are also promising alternative approaches.

Thermodynamic analysis and subsequent experimental validation demonstrate that rAOX is not obligately tied to alcohol production. Arresting rAOX at aldehydes opens a new route for the biosynthesis of these high-value chemicals^44^.

### rAOX-Mediated Upgrading of Waste-Derived Carboxylic Acid

We sought to apply rAOX to upgrade crude carboxylic acid effluents produced from waste (**Figure 4A**). Three different effluent streams of mixed carboxylic acids produced from food waste, wastewater, or industrial waste gases were tested. Each of these streams have different compositions and backgrounds due to their varying sourcing and production methods (See **Methods**, **Supplementary Table 3**, **Supplementary Table 4**). Cell-free processes are uniquely suited to receive these untreated effluents, as factors like sterility, presence of growth inhibitory factors, and catabolite repression are not of concern^41,45^. Since pre-treatment of these effluent streams prior to upgrading is expensive at scale, we opted to leave these effluent streams largely untreated.

**Figure 4:**
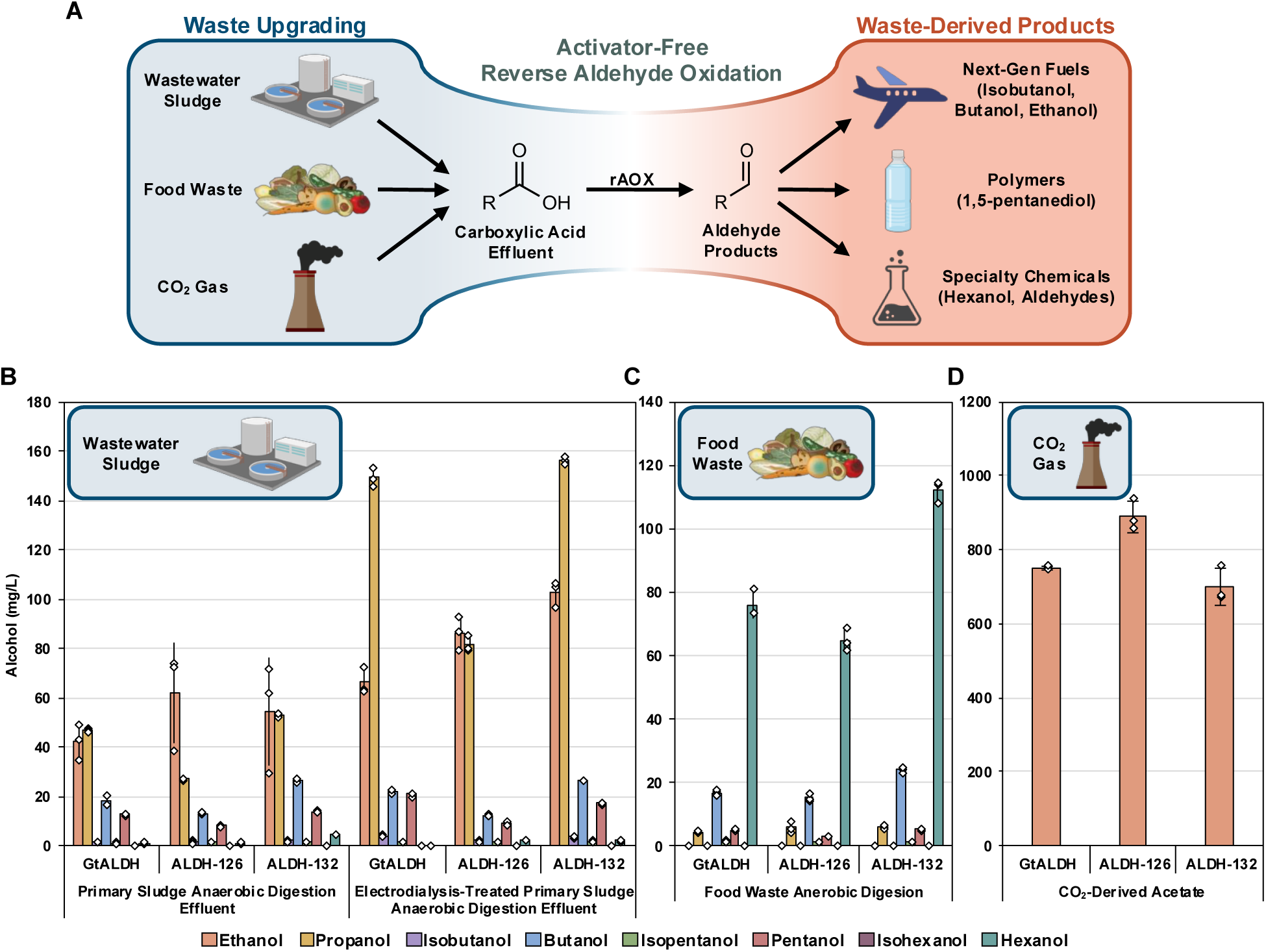
Waste Stream Valorization using rAOX. **A)** Waste streams are upgraded into carboxylic acid monomers. rAOX upgrades the carboxylic acid monomers into aldehydes. Aldehydes can be subsequently modified or directly used in the global bioeconomy. **B)** rAOX upgrading of effluent sourced from arrested anaerobic digestion of wastewater sludge. **C)** rAOX upgrading of effluent sourced from arrested anaerobic digestion of food waste. **D)** rAOX upgrading of effluent sourced from electrochemically generated acetate. Bars represent the average of three biological replicates. Error bars represent one standard deviation. A portion of this figure was generated at www.biorender.com.

Wastewater treatment plant-sourced sewage sludge anaerobic digestion solutions contained a broad range of aliphatic and branched carboxylic acids (**Supplementary Table 3, Supplementary Table 4**). We tested two iterations of these digestates: 1) a primary sludge anaerobic digestion effluent and 2) an electrodialysis-concentrated solution of the primary sludge anaerobic digestion effluent. When supplied to reactions with GtALDH, ALDH-126, or ALDH-132, a mixture of alcohols was produced, largely trending with the concentration of substrates, producing a total of 124 mg/L, 114 mg/L, and 156 mg/L C2-C6 alcohol products, respectively (**Figure 4B**). The electrodialyzed effluent contained a higher concentration of carboxylic acids, supporting more robust final alcohol titers, reaching 265 mg/L, 196 mg/L, and 311 mg/L of total C2-C6 alcohol production, respectively (**Figure 4B**).

Food waste-derived solutions were digested anaerobically under a condition shown to favor carboxylic acid chain elongation, enhancing butyrate and hexanoate yields (**Supplementary Table 4**). When supplied to ALDH-mediated reduction reactions, butanol (C4) and hexanol (C6) were produced as the primary products, 24 mg/L of butanol and 112 mg/L of hexanol ALDH-132 (**Figure 4C**). Small differences in product composition between GtALDH, ALDH-126, and ALDH-132-mediated systems suggest directed pairing of ALDH enzyme and target product can dictate specific product formation.

Finally, we evaluated CO_2_-derived acetate for ALDH-mediated reduction reactions. We previously demonstrated selective acetate production via a tandem CO_2_ electrolysis process, involving CO_2_ reduction to CO followed by CO reduction to acetate^23^. This process has a near 100% carbon efficiency^23^. Here, effluent from the electrochemical CO-to-acetate step was supplied to ALDH-mediated reactions. High acetate concentration in the effluent enabled a more concentrated acetate input to the reactions, when compared to anerobic digestate streams. After incubation, reactions containing GtALDH, ALDH-126, and ALDH-132 produced 752 mg/L, 890 mg/L, and 701 mg/L of ethanol respectively (**Figure 4D**). Because ethanol was produced as a side product during electrolysis, it was present at the start of the reactions (t = 0) at 425 mg/L. Accordingly, the ALDH-specific ethanol production was 327 mg/L, 465 mg/L, and 276 mg/L, respectively. Using this electrochemical method and rAOX sequentially, a 100% carbon conversion from CO_2_ to ethanol can be envisioned.

## Discussion

This work establishes rAOX as a new platform technology to directly reduce carboxylic acids to aldehydes and alcohols without the use of ATP, CoA, ferredoxins, or CO_2_ loss. This work mirrors the discovery that citrate synthase, classically regarded as catalyzing an irreversible step in the oxidative TCA cycle, surprisingly can reverse to enable rTCA^3,5^. Prior to this work, this ALDH enzyme class was nearly universally accepted by the scientific populous to be “irreversible”^8,9^, with only two prior works, to our knowledge, indicating that this activity was attainable^31,32^. Screening across 133 ALDH scaffolds demonstrated that this direct carboxylic acid reduction activity is not only attainable, but that it is remarkably prevalent throughout the ALDH protein family. When compared to the two prior examples of ALDHs performing this activity, we found that the *B. cereus* ALDH and *S. cerevisiae* ALDH described by Minteer^31^ and Tommasi^32^ were just the proverbial “tip-of-the-iceberg”, with enzymes throughout the ALDH family exhibiting strong carboxylic acid reduction capabilities.

Enzymes are catalysts, which do not change reaction thermodynamics. However, we found members of ALDH family exhibit vastly different capabilities to carry out reaction in oxidative versus reductive directions **(Supplementary Table 1, Supplementary Table 2)**. This difference may be attributed to the varied kinetic control exerted by enzymes^3,5,46^, such as different relative affinity for product versus reactant, and selective stabilizing of different transition states allowing different catalytic mechanisms. Further studies and engineering are needed to understand and augment the desirable traits of reversible ALDHs.

Instead of ATP activation, the driving forces that enable rAOX are two-fold: strong reducing power and thermodynamically downhill terminal steps to sequester the products. rTCA and rBOX employ the same principles^4,6^. The reducing power here is provided by NADH regeneration through a PTDH-catalyzed, highly exergonic reaction using phosphite as an inexpensive sacrificial electron source, while no oxygen-sensitive ferredoxins are required. Biomimetics of NAD^+^ with lower reduction potentials hold the promise to deliver stronger reducing power^47,48^. Unlike the energy sources ATP and ferredoxin, these simpler, noncanonical redox cofactors can be low-cost and more stable^49,50^. Ongoing work in our lab aims at powering rAOX with these noncanonical redox cofactors.

The product sequestering step demonstrated in this work is either accomplished by aldehyde reduction to alcohol catalyzed by ADH, or using in-situ extraction of aldehydes into organic phase. However, this final step can vary. Aldehydes’ reactivity make them highly versatile enzymatic substrates^44^. Integrating rAOX with other downstream reactions^51,52^, preferably ones that offer additional thermodynamic driving force, may enable access of a broad product scope from carboxylic acids.

We envision rAOX could also be implemented in vivo given engineered cellular and environmental conditions because no host-dependent ferredoxin cofactors are needed and all components of the enzymatic cascade have been shown to be readily translatable into industrial model organisms^6,16,38,48^, for which advanced metabolic engineering tools exist. To achieve this, metabolic engineering is needed to provide the desired redox environment in the cells. Since NAD(P)^+^/NADPH ratios fluctuate and are regulated by the cells, bio-orthogonal redox cofactor may also be needed to provide unwavering driving force^53^.

## Methods

### Molecular Cloning

ALDH genes were purchased from Integrated DNA Technologies as N-terminal polyhisitidine tagged synthetic gene constructs with terminal homologous arms for DNA assembly. Gene constructs were codon optimized for expression in *E. coli* using Integrated DNA Technologies’ codon optimization tool. Synthetic gene constructs were assembled into the expression vector by Gibson assembly. Gibson assembled products were transformed into *E. coli* XL1-Blue (Agilent Technologies) and plated on 2x Yeast Extract Tryptone media (2xYT, 16 g/L tryptone, 10 g/L yeast extract, 5 g/L NaCl) containing 100 mg/L ampicillin and incubated at 37 °C overnight. Solidified media for cell plating contained 15 g/L agar. After incubation, a single cell colony was picked from the plate and inoculated into 4 mL 2x YT media containing 100 mg/L ampicillin in a cell culture tube. The cultures were shaken at 37 °C overnight. Plasmids were isolated from the cultures using the Qiagen QIAprep Spin Miniprep Kit. ALDH gene sequences were confirmed by Sanger sequencing. Alcohol dehydrogenase from *Saccharomyces cerevisiae* (ScADH) was purchased as a lyophilized powder from Sigma Aldrich. Strains and plasmids used in this study are listed in **Supplementary Table 5**. The amino acid sequences of ALDHs are listed in **Supplementary Table 6**. The amino acid sequence of TcADH is listed in **Supplementary Table 7**.

### Protein expression

Plasmids containing ALDH or PTDH were transformed into *E. coli* BL-21 (Invitrogen) and plated on 2x YT media containing 100 mg/L ampicillin and incubated at 30 °C overnight. A single cell colony was picked from the plate and inoculated into 4 mL 2x YT media containing 100 mg/L ampicillin in a cell culture tube and incubated overnight at 30 °C. 500 µL of the cell culture was inoculated into 100 mL of 2x YT media containing 200 mg/L ampicillin in a 250 mL baffled shake flask. The flasks were shaken at 37 °C until the OD600 of the culture reached ∼ 0.6. The flasks were then shaken at room temperature for 30 minutes to cool. After cooling, the protein expression was induced by spiking isopropyl β-D-1-thiogalatopyranoside (IPTG) to a final concentration of 0.5 mM. The flasks were then shaken at room temperature overnight. Cells were harvested by centrifugation at 4000 xg at 4 °C for 12 minutes.

His-tagged proteins were purified from the cells using nickel-affinity resin (Hispur Ni-NTA Resin, ThermoFisher). Cell pellets were resuspended in ice-cold Binding Buffer containing 50 mM sodium phosphate buffer pH 7.7, 300 mM sodium chloride, 10 mM imidazole, and 0.03% Triton X-100. The resuspended cells were transferred to a pre-chilled 2 mL bead beating tube containing ∼0.5 mL 0.1 mm soda lime glass beads (BioSpec Products). The cells were disrupted by bead beading using a MP Biomedical FastPrep-24 by beating at 6 m/s for 35 seconds, followed by 2 minutes of cooling in an ice water bath. The lysis process was repeated five times for each sample. After lysis, the tubes were centrifuged at 20,000 xg at 4 °C for 20 minutes to clarify the lysates. The clarified lysates were transferred to clean, pre-chilled microcentrifuge tubes for protein immobilization. Ni-NTA resin was washed five times with Binding Buffer to remove any storage buffer. 150 µL of equivalent hydrated resin bed was added to each clarified lysate. The tubes were incubated at 4 °C for ∼1 hour using an end-over-end inverting rotator. The resin was separated from the lysate by centrifugation at 500 xg at 4 °C for 2 minutes. The resin was then transferred to a Zymo Research Zymo-Spin P1 column. The resins were washed twice with 300 µL of Wash Buffer containing 50 mM sodium phosphate pH 7.7, 300 mM sodium chloride, 50 mM imidazole, and 0.03% Triton X-100. The proteins were eluted with Elution Buffer containing 50 mM sodium phosphate buffer pH 7.7, 300 mM sodium chloride, and 250 mM imidazole. The purified proteins were mixed with a 50% glycerol solution to a final concentration of 20% glycerol for storage at - 80 °C until use. Protein concentration was determined by Bradford Assay using a dilution series of bovine serum albumin as standards.

### Generation of ALDH Sequence Similarity Network

Sequence similarity networks were generated using the EFI-EST webtool supplied with the ALDH protein family (Interpro PF00171) as the input query, filtered by UniRef50 sequences. Generated networks were visualized using Cytoscape.

### Screening ALDHs for carboxylic acid reduction activity in Round 1

Reactions contained 100 mM sodium phosphate pH 7.0, 100 mM sodium phosphite, 50 mM sodium hexanoate, 3 mM NADH, 0.3 mg/mL ALDH, 0.75 mg/mL ScADH and 1 mg/mL TS-PTDH at a final volume of 50 µL. A master mix of all components except ALDH was prepared on ice. ALDH was added to a 96-well PCR plate. To initiate the reactions, master mix was spiked into the PCR plate containing ALDH and mixed well. The reactions were then transferred to 2 mL glass vials sealed with a PTFE-lined cap. Reactions were incubated at 30 °C for 24 hours. After incubation, the reactions were mixed with an equal volume of ethyl acetate by shaking for 5 minutes. The mixture was clarified by centrifugation at 20,000 xg for two minutes and the organic layer was transferred to a GC vial for analysis.

### Screening ALDHs for carboxylic acid reduction activity in Round 2

Reactions contained 100 mM sodium phosphate pH 7.0, 100 mM sodium phosphite, 50 mM sodium hexanoate, 3 mM NADH, 0.3 mg/mL ALDH, 0.75 mg/mL *Saccharomyces cerevisiae* alcohol dehydrogenase (Sigma Aldrich), and 1 mg/mL TS-PTDH at a final volume of 30 µL. A master mix of all components except ALDH was prepared on ice. ALDH was added to a 96-well PCR plate. To initiate the reactions, master mix was spiked into the PCR plate containing ALDH and mixed well. The reactions were sealed with domed PCR tube cap strips. Reactions were incubated at 30 °C for 24 hours. After incubation, the reactions were mixed with 120 µL ethyl acetate thoroughly by pipetting. The PCR plate was then centrifuged at room temperature at 2000 xg for 5 minutes. The organic layer was transferred to a GC vial for analysis.

### Screening ALDHs for aldehyde oxidation activity

Specific activity assays measuring aldehyde oxidation of ALDH candidates were carried out as follows. A 200 μL reaction composed of 100 mM potassium phosphate buffer pH 7, 5 mM aldehyde (propionaldehyde or hexanal), 5 mM dithiothreitol, 3 mM cofactor (NAD^+^ or “no cofactor” addition of deionized water), and varying concentrations of purified ALDH (0.01-0.5 mg/mL) was initiated by the addition of purified ALDH to the reaction mixture. The reaction was subsequently transferred to a 96-well plate, and the formation of reduced cofactor was monitored kinetically at 340 nm. Reactions were carried out at 30 °C. All activity data was corrected by “no-cofactor” controls, where the enzymatic activity with NAD^+^ supplied was subtracted by the activity measured when the cofactor was substituted with water in the reaction mixture.

### Expansion of rAOX System to broader carboxylic acid substrates under optimized conditions

For reactions with aliphatic acid substrates (acetate, propionate, butyrate, and hexanoate), reactions contained 100 mM sodium phosphate pH 7.0, 100 mM sodium phosphite, 3 mM NADH, 10 g/L ALDH, 5 g/L ScADH, and 1.5 g/L TS-PTDH. Sodium acetate, sodium propionate, or sodium butyrate were supplied to reactions at 750 mM. Sodium hexanoate was supplied to reactions at 50 mM. A master mix of all components except ALDH was prepared on ice. ALDH was added to a 96-well PCR plate. To initiate the reactions, master mix was spiked into the PCR plate containing ALDH and mixed well. The reactions were transferred to a 2 mL glass vial and capped with a PTFE-lined cap. The reactions were incubated at 30 °C without shaking. Samples were taken periodically to monitor alcohol production. Samples were extracted by vigorously shaking with ethyl acetate at a 4:1 solvent to sample ratio. Samples were centrifuged at 20,000 xg for 2 minutes to separate the two fractions, and the organic layer was transferred to a GC vial for analysis. Sodium acetate, sodium propionate, and sodium hexanoate reactions were incubated for 72 hours. Sodium butyrate reactions were incubated for 96 hours. When detecting ethanol, low amounts of ethanol was found in control reactions without ALDH. Ultimately, this ethanol contamination was traced to the NADH purchased from Sigma Aldrich, which we theorized used ethanol in the final stages of the manufacturing process. Therefore, we subtracted the amount of ethanol from control reactions without ALDH (175 mg/L) from the amount of ethanol measured in samples containing ALDH to yield the final concentrations presented in this work. No other contaminating alcohols were observed for the other substrates tested in this manuscript.

Reactions with isobutyrate contained 100 mM sodium phosphate pH 7.0, 100 mM sodium phosphite, 3 mM NADH, 10 g/L ALDH, 1 g/L TcADH, and 1.5 g/L TS-PTDH. Sodium isobutyrate was supplied to the reactions at 522 mM. A master mix of all components except ALDH was prepared on ice. ALDH was added to a 96-well PCR plate. To initiate the reactions, master mix was spiked into the PCR plate containing ALDH and mixed well. The reactions were transferred to a 2 mL glass vial and capped with a PTFE-lined cap. The reactions were incubated at 30 °C without shaking for 96 hours. Samples were taken periodically to monitor alcohol production. Samples were extracted by vigorously shaking with ethyl acetate at a 4:1 solvent to sample ratio. Samples were centrifuged at 20,000 xg for 2 minutes to separate the two fractions, and the organic layer was transferred to a GC vial for analysis.

Reactions with 5-hydroxypentaonic acid contained 100 mM sodium phosphate pH 7.0, 100 mM sodium phosphite, 3 mM NADH, 10 g/L ALDH, 1 g/L TcADH, and 1.5 g/L TS-PTDH. Sodium 5-hydroxypentanoate was supplied to reactions at 750 mM. A master mix of all components except ALDH was prepared on ice. ALDH was added to a 96-well PCR plate. To initiate the reactions, master mix was spiked into the PCR plate containing ALDH and mixed well. The reactions were transferred to a 2 mL glass vial and capped with a PTFE-lined cap. The reactions were incubated at 30 °C without shaking for 158 hours. To terminate the reaction, 10 µL of sample was mixed with 30 µL of 250 mM sulfuric acid. The samples were subsequently filtered through a AcroPrep Advance 96-well filter plate 350 µL well volume, 0.2 µm wwPTFE membrane (Cytiva) by centrifugation at 1,500 xg for 5 minutes. The flowthrough was transferred to an HPLC vial for analysis.

For reactions with aromatic acid substrates (benzoate, trans-cinnamate, 4-hydroxybenzoate, or 2-furoate), reactions contained 100 mM sodium phosphate pH 7.0, 100 mM sodium phosphite, 3 mM NADH, 3 g/L ALDH, 0.75 g/L TcADH, and 2 g/L TS-PTDH. Trans-cinnamic acid, 4-hydroxybenozate, and 2-furoic acid were supplied at 50 mM. Sodium benzoate was supplied to reactions at 100 mM. A master mix of all components except ALDH was prepared on ice. ALDH was added to a 96-well PCR plate. To initiate the reactions, master mix was spiked into the PCR plate containing ALDH and mixed well. The reactions were transferred to a 2 mL glass vial and capped with a PTFE-lined cap. The reactions were incubated at 37 °C without shaking for 48 hours. Benzoate and trans-cinnamate samples were extracted by shaking vigorously with n-hexane at 2:1 solvent to sample ratio. 2-furoate samples were extracted with ethyl acetate at a 1:1 ratio. Samples were centrifuged at 20,000 xg for 2 minutes to separate the two fractions, and the organic layer was transferred to a GC vial for analysis. 4-hydroxybeznoate reactions were terminated by the addition of a solution containing 45% v/v methanol with 5% v/v formic acid. The terminating solution was added at a 2:1 solution:sample ratio. Reactions were mixed and then centrifuged at 20,000 xg for 15 minutes to separate out any precipitate and transferred to an HPLC vial for analysis.

### Investigation of Driving Forces for Improved rAOX Activity

Reactions investigating the driving forces with varying protein concentrations were performed similarly as stated above. Reactions contained 100 mM sodium phosphate pH 7.0, 100 mM sodium phosphite, 750 mM sodium propionate, 3 mM NADH, varying ALDH, varying ScADH, and varying PTDH concentrations at a final volume of 30 µL. A master mix of all components except ALDH and ScADH was prepared on ice. ALDH and ScADH were added to a 96-well PCR plate. To initiate the reactions, master mix was spiked into the PCR plate containing ALDH and ScADH and mixed well. The reactions were sealed with domed PCR tube cap strips. Reactions were incubated at 30 °C for 24 hours. After incubation, the reactions were mixed with 120 µL ethyl acetate thoroughly by pipetting. The PCR plate was then centrifuged at room temperature at 2000 xg for 5 minutes. The organic layer was transferred to a GC vial for analysis.

To investigate the effects of adding an organic overlay to the reactions for in situ extraction, reactions were performed in parallel with and without the organic overlay. Reactions contained 100 mM sodium phosphate pH 7.0, 100 mM sodium phosphite, 522 mM sodium isobutyrate, 3 mM NADH, 10 g/L ALDH, and 1.5 g/L PTDH at a final volume of 60 µL. A master mix of all components except ALDH was prepared on ice. ALDH was added to a 96-well PCR plate. To initiate the reactions, master mix was spiked into the PCR plate containing ALDH and mixed well. The reactions were transferred to a 2 mL glass vial and capped with a PTFE-lined cap. For the samples containing organic overlay, 300 µL of ethoxybenzene was added to the vials at t = 0. The reactions were incubated at 30 °C without shaking for 38 hours. After incubation, samples without organic overlay were extracted with a 5:1 ethoxybenzene:sample ratio by vortexing and then centrifuged at 20,000 xg for 2 minutes. The organic layer was transferred to a GC vial for analysis. For samples containing organic overlay, the samples were gently mixed to ensure consistent distribution of solutes in the overlay. Then, 20 µL of overlay was directly transferred to a GC vial for analysis.

### rAOX Upgrading of Waste-Stream-Derived Carboxylic Acid Effluents

Solutions from wastewater and food waste anaerobic digestion effluent streams were clarified by centrifugation at 20,000 xg for 5 minutes. Reactions for wastewater and food waste effluent streams contained 100 mM sodium phosphate pH 7.0, 100 mM sodium phosphite, 3 mM NADH, 0.75 g/L ALDH, 0.6 g/L TcADH, 1 g/L TS-PTDH, and a two-fold dilution of the centrifugally-clarified effluent stream. Reactions were initiated with the spiking of the clarified effluent stream. The reactions were performed in a 96-well PCR plate sealed with domed PCR cap strips, and they were incubated at 30 °C for 24 hours without shaking. Composition of the carboxylic acid profile in the reactions can be found in **Supplementary Table 6**. After incubation, the reactions were sampled by removing two volumes of liquid, one for extraction with ethyl acetate, one for extraction with ethoxybenzene. For ethyl acetate extraction, samples were extraction with a 4:1 v/v ethyl acetate:sample ratio by mixing vigorously. The ethyl acetate extractions were used to quantify alcohol species of equal or greater length than propanol. For ethoxybenzene extraction, samples were extracted with a 2:1:1 v/v/v ethoxybenzene:1 g/mL potassium phosphate dibasic solution:sample ratio by mixing vigorously. After mixing all samples were centrifuged at 20,000 xg for two minutes, and the organic layer was transferred to a GC vial for analysis. Ethoxybenzene extractions were used to quantify ethanol.

Gas-derived acetate effluent stream was generated in highly alkaline conditions (see Electrochemical Acetate Production section). Therefore, the solution was titrated to pH 7 using concentrated HCl prior to addition to reactions. Reactions for gas-derived acetate effluent contained 100 mM sodium phosphate pH 7.0, 100 mM sodium phosphite, 3 mM NAD^+^, 2 g/L ALDH, 1 g/L ScADH, 1 g/L TS-PTDH, and 750 mM acetate effluent stream. Reactions were initiated by spiking of the acetate effluent into the reactions. Reactions were performed in at 2 mL glass vial sealed with a PTFE-lined cap, and they were incubated at 30 °C for 48 hours. To analyze ethanol production, samples were extracted with a 2:1:1 v/v/v ethoxybenzene:1g/mL potassium phosphate dibasic solution:sample ratio by mixing vigorously. After mixing, all samples were centrifuged at 20,000 xg for two minutes, and the organic layer was transferred to a GC vial for analysis.

### Isobutyrate rAOX Reactions Comparing ScADH and TcADH

Reactions comparing ScADH and TcADH as alcohol dehydrogenases in isobutyrate reduction contained 100 mM sodium phosphate pH 7.0, 100 mM sodium phosphite, 200 mM sodium isobutyrate, 3 mM NADH, 0.5 g/L GtALDH, 0.7 g/L ScADH or TcADH, and 1 g/L PTDH. A master mix of all components except alcohol dehydrogenase was prepared on ice. Alcohol dehydrogenase was added to a 96-well PCR plate. To initiate the reactions, master mix was spiked into the PCR plate containing alcohol dehydrogenase and mixed well. The reactions were transferred to a 2 mL glass vial and capped with a PTFE-lined cap. The reactions were incubated at 30 °C without shaking. After 24 hours, the reactions were extracted with equal parts ethyl acetate by vigorously shaking. Samples were transferred to a microcentrifuge tube and centrifuged for 2 minutes at 20,000 xg to separate the two phases. The organic layer was transferred to a GC vial for analysis.

### Thermodynamic Calculations

Standard Gibbs free energies for carboxylic acid and aldehyde reduction reactions were calculated using the eQuilibrator biochemical thermodynamics calculator^39^. Standard Gibbs free energy of phosphite dehydrogenase reaction was determined from literature^54,55^. To determine the Gibbs free energy change of isobutyraldehyde partitioning into ethoxybenzene, solutions of ethoxybenzene with a known concentrations of isobutyraldehyde were prepared. Then, half of the volume of each solution was extracted with reaction buffer, and the remaining half was left untreated. The organic layer of the extracted sample and the untreated solution were collected and analyzed by gas chromatography. Peak areas were compared to a standard curve generated in the same manner of treatment to determine the absolute isobutyraldehyde concentration. The partition coefficient, as defined by Equation 1 was used to calculate ΔG° as defined by Equation 2.

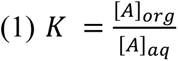

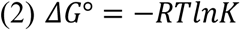

### Generation of Carboxylic Acid Effluents from Wastewater Treatment Plant

Carboxylic acid effluents derived from wastewater treatment plant sludge were produced as previously described^56^. Briefly, for the anaerobic digestion of primary sludge, the primary sludge was collected from the primary sedimentation unit of the Missouri River Wastewater Treatment Plant (St. Louis, MO, USA). Samples were stored at 4 °C prior to use as feedstock for a semi-continuous anaerobic digestion (AD) reactor. The AD reactor consisted of a 200 mL glass bottle operated at 37 °C with continuous shaking at 150 rpm. Every 3 days, 40 mL of digestate was replaced with 40 mL of fresh primary sludge supplemented with hydrogen peroxide (final concentration: 80 mg/L), yielding a solids retention time of 15 days. After each feeding, the reactor was flushed with nitrogen gas for 5 min to maintain anaerobic conditions. Detailed compositional characteristics of the primary sludge are provided in **Supplementary Table 5**.

To concentrate VFAs produced from the anaerobic digestion of primary sludge, VFAs were recovered from anaerobic digestate using a benchtop electrodialysis (ED) system (PCCell GmbH, Germany) equipped with alternating cation and anion exchange membranes to form two concentrate and two dilute chambers. The digestated primary sludge was centrifuged and filtered (0.45 μm, Bioemd Scientific, China), and the clarified liquid was fed into the dilute chamber. A 0.05 M H_2_SO_4_ solution served as the initial concentrate electrolyte. The ED system was operated in batch mode at 15 V for 2.5 h per cycle. After each cycle, the dilute chamber was replenished with 30 mL fresh digestate, while the concentrate was retained to accumulate VFAs. The pH of the concentrate was adjusted below 4 after each cycle to maintain VFAs in their molecular form and to minimize phosphate precipitation. A total of six ED enrichment cycles were performed for continuously concentrated VFAs from digested primary sludge.

### Generation of Carboxylic Acid Effluent from Food Waste

Roughly 120 lbs of University cafeteria food waste was naturally pre-fermented (uncontrolled pH and ambient temperature ∼17 °C) in a 900 L tank with a 225 L working volume for 16 days, resulting in ethanol, lactic, acetic, and butyric acid concentrations of 62.9, 128, 20.8, and 5.1 mM, respectively. After pre-fermentation, the food waste was centrifuged at 18,480 xg for 5 min, filtered through a 104 μm sieve filter to remove floating solids, and diluted with RO water (50/50 dilution). 1.5 L of this diluted filtrate was transferred into a 2 L agitated reactor, to which cornmeal solids (15 g/L, size range of 298 – 589 μm), and UASB granules (12.8 g/L, wastewater treatment system, New Belgium Brewery, Fort Collins, CO) were added. Batch anaerobic digestion was conducted at pH 5, 35 °C, 250 RPM (3.5 W/m^3^, marine impeller) for 15 days.

### Electrochemical Acetate Production

Acetate effluent streams were generated through CO electrolysis in a custom-designed zero-gap electrolyzer (5 cm² active area, serpentine flow field) operated at a constant current density of 200 mA cm⁻². The cathode was prepared by airbrushing a catalyst ink onto carbon paper (Sigracet 39BB, Fuel Cell Store). The catalyst ink contained commercial Cu nanopowder (40-60 nm, ≥99.5%, Sigma-Aldrich), PTFE (100-150 nm, Nanoshel-UK Ltd.), and Nafion ionomer (D2021, Ion Power, Inc.), each at 10 wt% relative to Cu, dispersed in 40 mL of a 1:1 (v/v) isopropanol-Milli-Q water mixture. The mixture was ultrasonicated for 30 minutes for dispersion, then sprayed onto a carbon paper placed on a 100 °C hot plate. The final catalyst loading was 1 mgCu cm⁻².

To fabricate the NiFeO_x_ anode, porous Ni foam (>99.99%, MTI Corporation) was ultrasonicated for 10 minutes in 5 M HCl and a 1:1 (v/v) ethanol-Milli-Q water solution, then rinsed with Milli-Q water. Electrodeposition was carried out in a three-electrode setup using the pretreated Ni foam as the working electrode, a platinized titanium felt (Fuel Cell Store) as the counter electrode, and an Ag/AgCl reference electrode. The electrolyte contained 3 mM Ni(NO₃)₂·6H₂O and 3 mM Fe(NO₃)₃·9H₂O. A potential of -1.0 V vs. Ag/AgCl was applied until 58 C of total charge was passed. After rinsing with Milli-Q water, the electrode was dried at 60 °C overnight and compressed at 5000 lbs using a Carver heated press.

The electrochemical acetate production was carried out in 1.4 M NaOH with a CO feed rate at 30 sccm to produce different acetate-rich effluents. A peristaltic pump circulated 100 mL of electrolyte through the anode chamber at 5 mL min⁻¹. The electrolyte was replenished once the acetate concentration reached the target level (cation-to-acetate ratio of 1:0.85), as indicated by a pH value of 12.7. Gas and liquid products were analyzed using online gas chromatography and high-performance liquid chromatography, respectively.

### Analytical Methods

All GC analysis was performed on an Agilent 6890 gas chromatograph equipped with a split/splitless inlet and a flame-ionization detector. Separations were performed using an Agilent DB-WAX UI column (30 m x 0.25 mm x 0.5 µm). For all analyses, helium was used as the carrier gas. Hydrogen and air were supplied to the FID at 30 mL/min and 350 mL/min, respectively. Nitrogen was used as a makeup gas with a flowrate of 25 mL/min. The instrument was operated in constant flow mode. Oven programs and injection parameters for each separation method are detailed below. The front inlet was maintained at 250 °C. The FID was maintained at 300 °C

For detection of hexanol. The oven was held at 100 °C for 2 minutes, then ramped at 20 °C/min to 230 °C, then held at 230 °C for 5 min. The carrier gas flow rate was 2 mL/min. 2 µL of sample was injected. The split ratio was 5:1.

For the detection of propanol, butanol, isobutanol, and isobutyraldehyde, the oven was held at 65 °C for 4 min, then ramped at 100 °C/min to 230 °C, then held at 230 °C for 2.4 min. The carrier gas flowrate was 2 mL/min. 1 µL of sample was injected. The split ratio was 10:1. This GC method was used for samples extracted in ethyl acetate and ethoxybenzene.

For detection of ethanol, the oven was held at 40 °C for 5.5 minutes, then ramped at 70 °C/min to 230 °C, then held at 230 °C for 3.3 min. The carrier gas flowrate was 2 mL/min. 1 µL of sample was injected. The split ratio was 10:1.

For detection of benzyl alcohol and trans-cinnamyl alcohol, the oven was held at 120 °C for 2 min, then ramped at 10 °C/min to 230 °C, then held at 230 °C for 5 min. The carrier gas flowrate was 2 mL/min. 1 µL of sample was injected. The split ratio was 5:1.

For detection of furfuryl alcohol, the oven was held at 50 °C for 5 min, then ramped at 10 °C/min to 230 °C, then held at 230 °C for 12 minutes. The carrier gas flowrate was 1.8 mL/min. 1 µL of sample was injected. The split ratio was 5:1.

All HPLC analysis was carried out on an Agilent 1100 HPLC equipped with a diode array detector and a refractive index detector. For the detection of 1,5-pentanediol, a Bio-Rad Fast Acid Analysis HPLC Column (100 x 7.8 mm) was used for separation. A mobile phase of 20 mM sulfuric acid was flowed through the column at 1 mL/min. The column was maintained at 55 °C. The refractive index detector was maintained at 55 °C. 15 µL of sample was injected into the system. For the detection of 4-hydroxybenzyl alcohol, a Phenomenex Luna Omega 5µm Polar C18 100Å column (150 x 4.6 mm) was used for separation. Isocratic separation was performed using a mobile phase of 50:50 0.1% (v/v) formic acid:0.1% (v/v) formic acid in methanol was flowed through the column at 1 mL/min. The column was maintained at 40 °C. The diode array detector was monitored at 240 nm. 5 µL of sample was injected.

## Author Contributions

W.B.B., S.S., and H.L. designed all experiments apart from the generation of carboxylic acid effluent streams. E.L., Y.C., S.S., and J.B.S. performed bioinformatic analysis and sequence similarity network construction. E.L., Y.C., S.S., W.B.B., J.B.S., and H.L. selected the ALDH candidates. S.S. performed the aldehyde oxidation experiments. W.B.B., S.P., S.S., S.A., D.H, A.W., C.C., and R.H. performed the carboxylic acid reduction experiments. J.S. and Z.H. performed the production of short chain fatty acids from wastewater treatment plant-derived sludge. S.L. and P.C.G. performed the production of short chain fatty acids from food waste. Z.X.W. and F.J. conducted the electrolysis experiments. All authors analyzed the data, wrote the manuscript, and approved its contents.

## Acknowledgements

W.B.B., S.S., S.P., S.A., D.H, A.W., C.C., R.H., and H.L. acknowledge support from Advanced Research Projects Agency–Energy EcoSynBio program (Award no. DE-AR0001508), the National Science Foundation (NSF) (Award no. 2328145), the National Institutes of Health (NIH) (Award no. 1R35GM153401-01), Department of Energy (DOE) (Award no. EE0008923) and Sloan Research Fellowship. E.L., Y.C. and J.B.S. acknowledge the funding from Advanced Research Projects Agency–Energy EcoSynBio program (Award no. DE-AR0001508), the National Institute of Environmental Health Sciences (Grant no. P42ES004699), the NIH (Grant no. R01 GM 076324-11), and the NSF (Grant nos. 2328145, 1627539, 1805510, and 1827246). S.L. and P.C.G. acknowledge funding from the U.S. Department of Energy, Bioenergy Technologies Office, (Award no. DE-EE0008923). Z.X.W. and F.J. acknowledge financial support from the NSF (Award no. 2330245). J.S. and Z.H. acknowledge the support of the National Science Foundation (Award no. 2150613).

## References

1. Evans, M. C., Buchanan, B. B. & Arnon, D. I. A new ferredoxin-dependent carbon reduction cycle in a photosynthetic bacterium. Proc. Natl. Acad. Sci. 55, 928–934 (1966).

2. Shiba, H., Kawasumi, T., Igarashi, Y., Kodama, T. & Minoda, Y. The CO2 assimilation via the reductive tricarboxylic acid cycle in an obligately autotrophic, aerobic hydrogen-oxidizing bacterium, Hydrogenobacter thermophilus. Arch. Microbiol. 141, 198–203 (1985).

3. Nunoura, T. et al. A primordial and reversible TCA cycle in a facultatively chemolithoautotrophic thermophile. Science 359, 559–563 (2018).

4. Steffens, L. et al. High CO2 levels drive the TCA cycle backwards towards autotrophy. Nature 592, 784–788 (2021).

5. Mall, A. et al. Reversibility of citrate synthase allows autotrophic growth of a thermophilic bacterium. Science 359, 563–567 (2018).

6. Dellomonaco, C., Clomburg, J. M., Miller, E. N. & Gonzalez, R. Engineered reversal of the β-oxidation cycle for the synthesis of fuels and chemicals. Nature 476, 355–359 (2011).

7. Lian, J. & Zhao, H. Reversal of the β-Oxidation Cycle in Saccharomyces cerevisiae for Production of Fuels and Chemicals. ACS Synth. Biol. 4, 332–341 (2015).

8. Shortall, K., Djeghader, A., Magner, E. & Soulimane, T. Insights into Aldehyde Dehydrogenase Enzymes: A Structural Perspective. Front. Mol. Biosci. 8, (2021).

9. Heider, J. & Hege, D. The aldehyde dehydrogenase superfamilies: correlations and deviations in structure and function. Appl. Microbiol. Biotechnol. 109, 106 (2025).

10. Nielsen, J. Synthetic Biology for Engineering Acetyl Coenzyme A Metabolism in Yeast. mBio 5, 10.1128/mbio.02153-14 (2014).

11. Zhang, S. et al. Metabolic engineering for efficient supply of acetyl-CoA from different carbon sources in Escherichia coli. Microb. Cell Factories 18, 130 (2019).

12. Winkler, M. Carboxylic acid reductase enzymes (CARs). Curr. Opin. Chem. Biol. 43, 23–29 (2018).

13. Kunjapur, A. M., Tarasova, Y. & Prather, K. L. J. Synthesis and Accumulation of Aromatic Aldehydes in an Engineered Strain of Escherichia coli. J. Am. Chem. Soc. 136, 11644–11654 (2014).

14. Nissen, L. S. & Basen, M. The emerging role of aldehyde:ferredoxin oxidoreductases in microbially-catalyzed alcohol production. J. Biotechnol. 306, 105–117 (2019).

15. Basen, M. et al. Single gene insertion drives bioalcohol production by a thermophilic archaeon. Proc. Natl. Acad. Sci. 111, 17618–17623 (2014).

16. Wang, J., Lin, M., Xu, M. & Yang, S.-T. Anaerobic Fermentation for Production of Carboxylic Acids as Bulk Chemicals from Renewable Biomass. in Anaerobes in Biotechnology (eds Hatti-Kaul, R., Mamo, G. & Mattiasson, B.) 323–361 (Springer International Publishing, Cham, 2016). doi:10.1007/10_2015_5009.

17. Bogorad, I. W., Lin, T.-S. & Liao, J. C. Synthetic non-oxidative glycolysis enables complete carbon conservation. Nature 502, 693–697 (2013).

18. Hellgren, J., Godina, A., Nielsen, J. & Siewers, V. Promiscuous phosphoketolase and metabolic rewiring enables novel non-oxidative glycolysis in yeast for high-yield production of acetyl-CoA derived products. Metab. Eng. 62, 150–160 (2020).

19. Holtzapple, M. T. et al. Microbial communities for valorizing biomass using the carboxylate platform to produce volatile fatty acids: A review. Bioresour. Technol. 344, 126253 (2022).

20. Bhatt, A. H., Ren, Z. (Jason) & Tao, L. Value Proposition of Untapped Wet Wastes: Carboxylic Acid Production through Anaerobic Digestion. iScience 23, (2020).

21. Liew, F. E. et al. Carbon-negative production of acetone and isopropanol by gas fermentation at industrial pilot scale. Nat. Biotechnol. 40, 335–344 (2022).

22. Wang, H. et al. CO2 electrolysis toward acetate: A review. Curr. Opin. Electrochem. 39, 101253 (2023).

23. Crandall, B. S. et al. Kilowatt-scale tandem CO2 electrolysis for enhanced acetate and ethylene production. *Nat*. Chem. Eng. 1, 421–429 (2024).

24. Biofuels and Bioproducts from Wet and Gaseous Waste Streams: Challenges and Opportunities. Energy.gov https://www.energy.gov/eere/bioenergy/articles/biofuels-and-bioproducts-wet-and-gaseous-waste-streams-challenges-and (2017).

25. Kutscha, R. & Pflügl, S. Microbial Upgrading of Acetate into Value-Added Products— Examining Microbial Diversity, Bioenergetic Constraints and Metabolic Engineering Approaches. Int. J. Mol. Sci. 21, 8777 (2020).

26. Jinich, A. et al. A thermodynamic atlas of carbon redox chemical space. Proc. Natl. Acad. Sci. 117, 32910–32918 (2020).

27. Bar-Even, A., Flamholz, A., Noor, E. & Milo, R. Thermodynamic constraints shape the structure of carbon fixation pathways. Biochim. Biophys. Acta BBA - Bioenerg. 1817, 1646– 1659 (2012).

28. Bar-Even, A., Flamholz, A., Noor, E. & Milo, R. Rethinking glycolysis: on the biochemical logic of metabolic pathways. Nat. Chem. Biol. 8, 509–517 (2012).

29. Dixon, M. & Lutwak-Mann, C. Aldehyde Mutase. Nature 139, 548–549 (1937).

30. Racker, E. Aldehyde Dehydrogenase, a Diphosphopyridine Nucleotide-Linked Enzyme. J. Biol. Chem. 177, 883–892 (1949).

31. Kummer, M. J. et al. Substrate Channeling by a Rationally Designed Fusion Protein in a Biocatalytic Cascade. JACS Au 1, 1187–1197 (2021).

32. Pietricola, G. et al. Covalent Immobilization of Aldehyde and Alcohol Dehydrogenases on Ordered Mesoporous Silicas. Waste Biomass Valorization 13, 4043–4055 (2022).

33. Cazelles, R. et al. Reduction of CO 2 to methanol by a polyenzymatic system encapsulated in phospholipids–silica nanocapsules. New J. Chem. 37, 3721–3730 (2013).

34. Guo, B., Ji, X., Xue, Y. & Huang, Y. Formaldehyde dehydrogenase SzFaldDH: an indispensable bridge for relaying CO 2 bioactivation and conversion. Green Chem. 26, 11540–11547 (2024).

35. Singh, R. K. et al. Insights into Cell-Free Conversion of CO2 to Chemicals by a Multienzyme Cascade Reaction. ACS Catal. 8, 11085–11093 (2018).

36. Kuk, S. K., et al. Photoelectrochemical Reduction of Carbon Dioxide to Methanol through a Highly Efficient Enzyme Cascade. Angew. Chem. Int. Ed. 56, 3827–3832 (2017).

37. Li, X. et al. Characterization of a broad-range aldehyde dehydrogenase involved in alkane degradation in *Geobacillus thermodenitrificans* NG80-2. Microbiol. Res. 165, 706–712 (2010).

38. Brat, D., Weber, C., Lorenzen, W., Bode, H. B. & Boles, E. Cytosolic re-localization and optimization of valine synthesis and catabolism enables increased isobutanol production with the yeast Saccharomyces cerevisiae. Biotechnol. Biofuels 5, 65 (2012).

39. Flamholz, A., Noor, E., Bar-Even, A. & Milo, R. eQuilibrator—the biochemical thermodynamics calculator. Nucleic Acids Res. 40, D770–D775 (2012).

40. Hong, S.-H. et al. Alternative Biotransformation of Retinal to Retinoic Acid or Retinol by an Aldehyde Dehydrogenase from Bacillus cereus. Appl. Environ. Microbiol. 82, 3940–3946 (2016).

41. Sherkhanov, S. et al. Isobutanol production freed from biological limits using synthetic biochemistry. Nat. Commun. 11, 4292 (2020).

42. Atsumi, S., Higashide, W. & Liao, J. C. Direct photosynthetic recycling of carbon dioxide to isobutyraldehyde. Nat. Biotechnol. 27, 1177–1180 (2009).

43. Rodriguez, G. M. & Atsumi, S. Isobutyraldehyde production from Escherichia coli by removing aldehyde reductase activity. Microb. Cell Factories 11, 90 (2012).

44. Zhou, J., Chen, Z. & Wang, Y. Bioaldehydes and beyond: Expanding the realm of bioderived chemicals using biogenic aldehydes as platforms. Curr. Opin. Chem. Biol. 59, 37–46 (2020).

45. Kay, J. E. & Jewett, M. C. A cell-free system for production of 2,3-butanediol is robust to growth-toxic compounds. Metab. Eng. Commun. 10, e00114 (2020).

46. Rauwerdink, A. et al. Evolution of a Catalytic Mechanism. Mol. Biol. Evol. 33, 971–979 (2016).

47. Weusthuis, R. A., Folch, P. L., Pozo-Rodríguez, A. & Paul, C. E. Applying Non-canonical Redox Cofactors in Fermentation Processes. iScience 23, (2020).

48. Black, W. B., Perea, S. & Li, H. Design, construction, and application of noncanonical redox cofactor infrastructures. Curr. Opin. Biotechnol. 84, 103019 (2023).

49. A. Rollin, J., Kin Tam, T. & Percival Zhang, Y.-H. New biotechnology paradigm: cell-free biosystems for biomanufacturing. Green Chem. 15, 1708–1719 (2013).

50. Zachos, I., Döring, M., Tafertshofer, G., Simon, R. C. & Sieber, V. carba Nicotinamide Adenine Dinucleotide Phosphate: Robust Cofactor for Redox Biocatalysis. Angew. Chem. Int. Ed. 60, 14701–14706 (2021).

51. Chou, A., Lee, S. H., Zhu, F., Clomburg, J. M. & Gonzalez, R. An orthogonal metabolic framework for one-carbon utilization. Nat. Metab. 3, 1385–1399 (2021).

52. Liu, J., Hsu, C.-C. & Wong, C.-H. Sequential aldol condensation catalyzed by DERA mutant Ser238Asp and a formal total synthesis of atorvastatin. Tetrahedron Lett. 45, 2439–2441 (2004).

53. Aspacio, D. et al. Shifting redox reaction equilibria on demand using an orthogonal redox cofactor. Nat. Chem. Biol. 20, 1535–1546 (2024).

54. White, A. K. & Metcalf, W. W. Microbial Metabolism of Reduced Phosphorus Compounds. Annu. Rev. Microbiol. 61, 379–400 (2007).

55. Costas, A. M. G., White, A. K. & Metcalf, W. W. Purification and Characterization of a Novel Phosphorus-oxidizing Enzyme from Pseudomonas stutzeri WM88 * 210. J. Biol. Chem. 276, 17429–17436 (2001).

56. Sun, J., Zhang, X., Guan, J. & He, Z. Volatile Fatty Acid Production through Arresting Methanogenesis by Electro-Synthesized Hydrogen Peroxide in Anaerobic Digestion and Subsequent Recovery by Electrodialysis. ACS EST Eng. 4, 2964–2973 (2024).

