## Supplementary Information for "Activation-Free Upgrading of Carboxylic Acids to Aldehydes and Alcohols"

**Affiliations:**
[1]Department of Chemical and Biomolecular Engineering, University of California, Irvine, Irvine, USA. [2]Genome Center, University of California, Davis, Davis, CA, USA. [3]Department of Energy, Environmental and Chemical Engineering, Washington University in St. Louis, St. Louis, MO, USA. [4]Karen M Swindler Department of Chemical and Biological Engineering, South Dakota School of Mines and Technology, Rapid City, SD, USA. [5]Department of Pharmaceutical Sciences, University of California, Irvine, Irvine, CA, USA. [6]Department of Chemistry, University of California, Davis, Davis, CA, USA. [7]Department of Biochemistry and Molecular Medicine, University of California, Davis, Davis, CA, USA. [8]UCI Center for Synthetic Biology (CSB), University of California, Irvine, Irvine, CA, USA. [9]Department of Biological Chemistry, University of California, Irvine, Irvine, CA, USA.

[α]These authors contributed equally: William B. Black, Samer Saleh.

*Corresponding author

**Corresponding Author**

**Han Li -** *Department of Chemical and Biomolecular Engineering, University of California, Irvine, Irvine, California 92697, USA; UCI Center for Synthetic Biology (CSB), University of California, Irvine, Irvine, California 92697-3900, USA; Department of Biological Chemistry, University of California, Irvine, Irvine, California 92697-3900, USA.*

### Supplementary Information

#### A. Supplementary Figures and Tables

Supplementary Figure 1: Dropout Control Reactions for Carboxylic Acid Reduction Screening Platform.

Supplementary Figure 2: Optimization of rAOX System Substrates.

Supplementary Figure 3: TcADH supports improved isobutanol production in GtALDH rAOX System.

Supplementary Figure 4: Combinatorial Testing of Protein Supplementation in rAOX System.

Supplementary Table 1: Round 1 ALDH Activity Screening.

Supplementary Table 2: Round 2 ALDH Activity Screening.

Supplementary Table 3: Characteristics of Primary Sludge from Wastewater Treatment Plant.

Supplementary Table 4: Carboxylic Acid Concentrations in Waste Stream Upgrading Reactions.

Supplementary Table 5: Strains and Plasmids Used in this Study.

Supplementary Table 6: Amino Acid Sequences of Bioprospected Aldehyde Dehydrogenases.

Supplementary Table 7: Amino Acid Sequence of TcADH.

#### B. Supplementary References

### A. Supplementary Figures and Tables

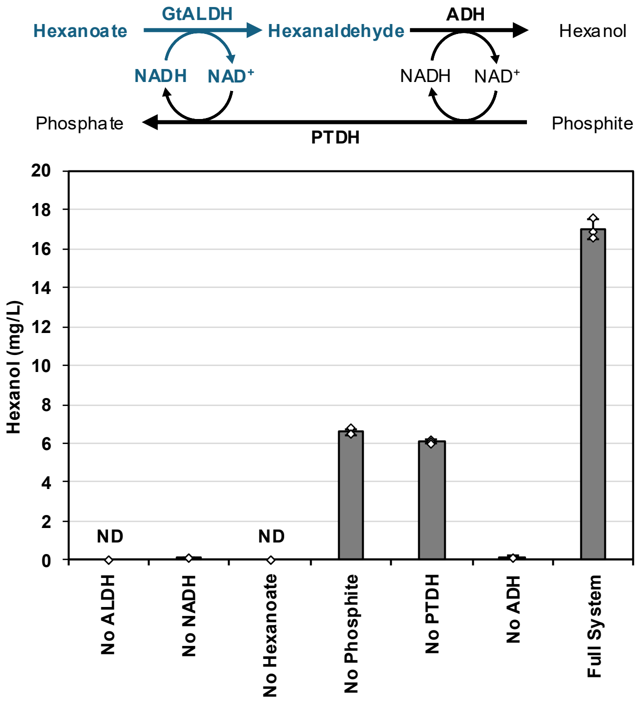


**Supplementary Figure 1: Dropout Control Reactions for Carboxylic Acid Reduction Screening Platform.** The platform is constructed of three proteins, aldehyde dehydrogenase (ALDH), alcohol dehydrogenase (ADH), and phosphite dehydrogenase (PTDH). Sodium hexanoate is fed to the enzymatic system, where ALDH catalyzes the reduction of hexanoate to hexanaldehyde using NADH. Hexanaldehyde is the sequestered away from the ALDH by further reduction to hexanol, catalyzed by ADH using NADH. Oxidized NADH ($NAD^+$) is reduced back to NADH by PTDH. Individual components were removed to examine the effect on hexanol production. Reactions contained 100 mM sodium phosphate pH 7.0, 100 mM sodium phosphite, 3 mM NADH, 50 mM sodium hexanoate, 0.3 g/L GtALDH, 0.75 g/L ScADH, and 1.0 g/L PTDH. The reactions were performed in 2 mL glass vials sealed with PTFE-lined cap. Reactions were incubated at 30 °C for 24 hours. Values represent an average of three biological replicates. Error bars represent one standard deviation. “ND”: hexanol not detected.

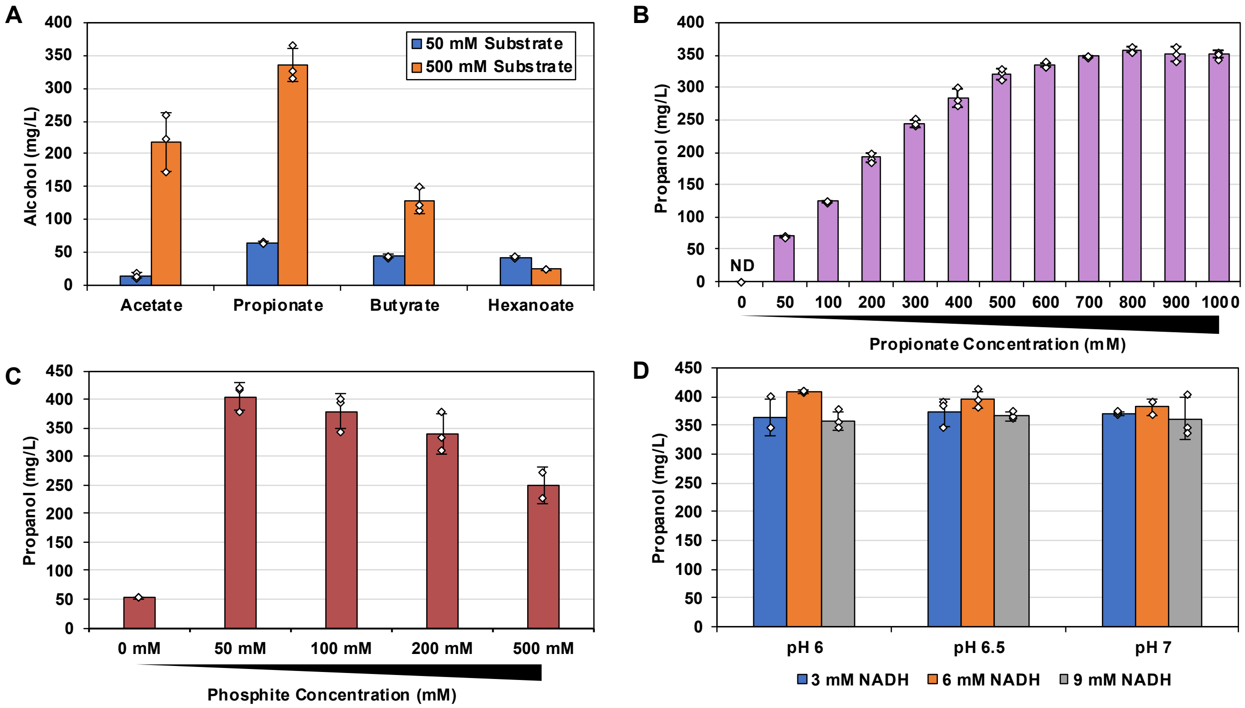


**Supplementary Figure 2: Optimization of rAOX System Substrates**

**A)** GtALDH has been previously characterized to exhibit aldehyde oxidation activity across a broad scope of aldehyde substrates[1]. Consistent with this, GtALDH exhibited strong rAOX activity using carboxylic acid substrates of varying carbon chain lengths. Reactions contained 100 mM sodium phosphate pH 7.0, 100 mM sodium phosphite, 3 mM NADH, 50 or 500 mM sodium salts of carboxylic acids, 0.75 g/L ALDH, 0.75 g/L ScADH, and 1 g/L PTDH. **B)** Step-wise gradient of sodium propionate supplementation to the GtALDH-mediated rAOX system. Reactions contained 100 mM sodium phosphate pH 7.0, 100 mM sodium phosphite, 3 mM NADH, varying sodium propionate, 0.75 g/L ALDH, 0.75 g/L ScADH, 1.0 g/L ALDH. **C)** Gradient of phosphite supplementation to the GtALDH-mediated rAOX system. Reactions contained 100 mM sodium phosphate pH 7.0, 3 mM NADH, 500 mM sodium propionate, 0.75 g/L GtALDH, 0.75 g/L ScADH, 1.0 g/L PTDH. **D)** Pairwise screening of reaction pH and NADH concentration in rAOX. These parameters were optimized together due to high rates of reduced redox cofactor (NADH) degradation in low pH solutions[2]. We hypothesized that increasing NADH concentrations may enable the retention of a NADH pool for catalysis at lower pH levels while simultaneously supporting improved catalysis, if NADH was limiting. However, under the conditions tested, changes in propanol production were small. 3 mM NADH supplementation was selected for future experiments to aid overall process economics. Reactions contained 100 mM sodium phosphate at pH 6.0-7.0, 100 mM sodium phosphite, varying NADH concentrations, 500 mM sodium phosphite, 0.75 g/L ALDH, 0.75 g/L ScADH, 1.0 g/L PTDH. All reactions were performed in 2 mL glass vials sealed with a PTFE-lined cap, and they were incubated at 30 °C for 24 hours. Values represent an average of three biological replicates. In subfigure C, value for reactions containing 500 mM phosphite was an average of two biological replicates. Error bars represent one standard deviation. “ND”: alcohol product not detected.

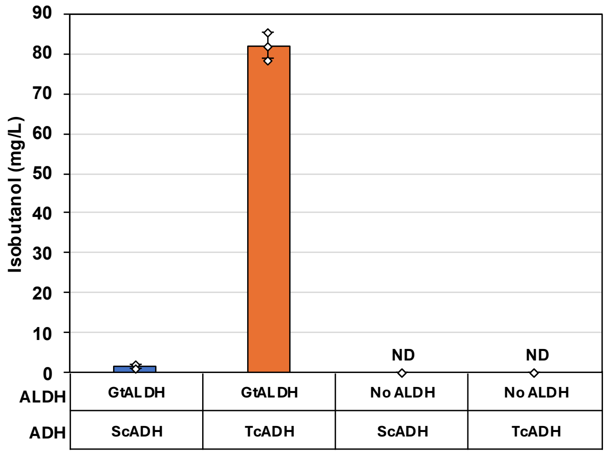


**Supplementary Figure 3: TcADH Supports Improved Isobutanol Production in GtALDH rAOX System.** Reactions contained 100 mM sodium phosphate pH 7.0, 100 mM sodium phosphite, 200 mM sodium isobutyrate, 3 mM NADH, 0.5 g/L ALDH, 0.7 g/L ScADH or TcADH, and 1.0 g/L PTDH. Reactions were performed in 2mL glass vials sealed with a PTFE-lined cap. Reactions were incubated at 30 °C for 24 hours. Values represent the average of three biological replicates, unless stated otherwise. Value for GtALDH+ScADH is an average of two biological replicates. Error bars represent one standard deviation. “ND”, alcohol product not detected.

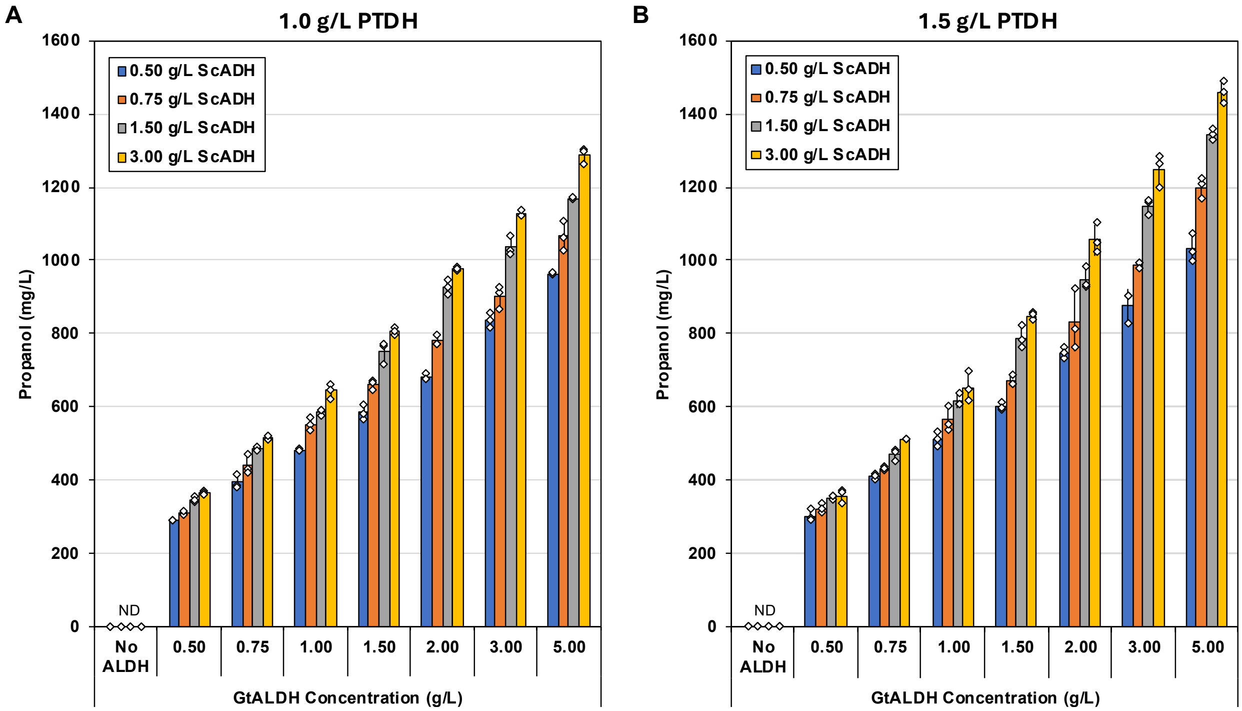


**Supplementary Figure 4: Combinatorial Testing of Protein Supplementation in rAOX System.** Combinatorial test of ALDH and ADH concentration in rAOX at 1.0 g/L PTDH **(A)** and 1.5 g/L PTDH **(B)**. GtALDH-mediated rAOX system was given different concentrations of ALDH, ScADH, and PTDH. Reactions were incubated in 2 mL glass vials sealed with a PTFE-lined cap. Reactions were incubated at 30 °C for 24 hours. Values represent the average of three biological replicates. The 3 g/L ALDH+3 g/L ADH+1.0 g/L PTDH condition is an average of two biological replicates. Error bars represent one standard deviation. “ND”, alcohol product not detected.

**Supplementary Table 1:** Round 1 ALDH Activity Screening

| Uniprot ID | ALDH # | Hexanal Oxidation Activity (nmol/mg/min) | Hexanoate Reduction Activity (mg hexanol/L) |
|---|---|---|---|
| P47771 | ALDH-001 | 222.1 ± 6.3 | ND |
| P54114 | ALDH-002 | 171.9 ± 12.8 | 3.88 ± 0.75 |
| P46367 | ALDH-003 | 45.2 ± 1.4 | 0.02 ± 0.04 |
| P40047 | ALDH-004 | 11.3 ± 1.3 | 0.02 ± 0.04 |
| P54115 | ALDH-005 | 22.3 ± 0.6 | 0.02 ± 0.04 |
| Q04458 | ALDH-006 | 20.1 ± 0.8 | 5.71 ± 1.16 |
| A0A6P3Q7G7 | ALDH-007 | 146.3 ± 5.6 | 0.04 ± 0.04 |
| A0A5J5DF59 | ALDH-008 | 152.4 ± 13.9 | 3.75 ± 0.79 |
| A0A0D6KE96 | ALDH-009 | 322.2 ± 93.2 | 0.05 ± 0.09 |
| L8N0N6 | ALDH-010 | 1481.9 ± 23.4 | 0.06 ± 0.05 |
| A0A2V7VB71 | ALDH-011 | 4.1 ± 3.8 | ND |
| UPI000407F0B2 | ALDH-012 | 1.1 ± 0.2 | 0.02 ± 0.04 |
| A0A2Z4LU87 | ALDH-013 | 423.2 ± 4.9 | 4.11 ± 0.82 |
| A0A0S8BPB2 | ALDH-014 | 135.7 ± 4.3 | 0.03 ± 0.06 |
| UPI000E14A9E8 | ALDH-015 | ND | ND |
| A0A7Y1V182 | ALDH-016 | 4.2 ± 2.1 | ND |
| A0A517R638 | ALDH-017 | ND | ND |
| A0A1F8MB45 | ALDH-018 | ND | 0.06 ± 0.10 |
| A0A346XWA7 | ALDH-019 | 3.9 ± 1.8 | 1.71 ± 0.21 |
| A0A2E3KN37 | ALDH-020 | 214.1 ± 2.2 | ND |
| UPI000A40ADAD | ALDH-021 | ND | 3.13 ± 0.63 |
| A0A139NAP6 | ALDH-022 | ND | 0.03 ± 0.05 |
| A0A1X6ZLY8 | ALDH-023 | 371.8 ± 24.6 | ND |
| A0A060QGV9 | ALDH-024 | ND | ND |
| UPI0001E31496 | ALDH-025 | ND | 0.04 ± 0.08 |
| A0A537X7T4 | ALDH-026 | ND | 2.93 ± 0.52 |
| A0A2E8CRC4 | ALDH-027 | ND | 0.50 ± 0.09 |
| A0A2V7BEQ0 | ALDH-028 | ND | 4.50 ± 0.67 |
| A0A1T1H988 | ALDH-029 | 489.4 ± 60.0 | 3.56 ± 0.50 |
| A0A382Q6S7 | ALDH-030 | ND | ND |
| M4Z1V7 | ALDH-031 | 55.6 ± 1.1 | ND |
| A0A420XXS9 | ALDH-032 | ND | ND |
| A0A1F6LNY1 | ALDH-033 | ND | 1.02 ± 0.16 |
| A0A7H4GQ81 | ALDH-034 | 27.8 ± 11.5 | 0.29 ± 0.06 |
| R7Z1F3 | ALDH-035 | ND | 0.85 ± 0.15 |
| A0A3D2UMU3 | ALDH-036 | 43.7 ± 4.6 | ND |
| P05091 | ALDH-037 | 79.1 ± 4.1 | ND |
| P51647 | ALDH-038 | 363.8 ± 5.4 | 0.03 ± 0.05 |
| P77674 | ALDH-039 | 453.7 ± 86.6 | 0.03 ± 0.05 |
| P23883 | ALDH-040 | 2117.9 ± 53.6 | 0.54 ± 0.01 |
| UPI001AE6B188 | ALDH-041 | 23.8 ± 0.2 | 0.26 ± 0.02 |

| | | | |
|---|---|---|---|
| A4IT08 | ALDH-042 (GtALDH) | 60.5 ± 5.2 | 13.45 ± 0.86 |
| P25526 | ALDH-043 | 53.2 ± 2.6 | 0.05 ± 0.05 |
| See footnote[‡] | ALDH-044 | 389.8 ± 10.3 | 0.08 ± 0.01 |
| P51977 | ALDH-045 | 261.5 ± 5.6 | 3.17 ± 0.31 |
| UPI00019FFBE0 | ALDH-046 | 55.2 ± 0.8 | 0.19 ± 0.07 |
| Q9LRI6 | ALDH-047 | 13.4 ± 0.4 | 0.02 ± 0.03 |
| C9DIJ2 | ALDH-048 | 261.2 ± 15.4 | 0.49 ± 0.07 |
| Q65NX0 | ALDH-049 | 1759.3 ± 31.1 | 0.14 ± 0.12 |
| UPI00005BF137 | ALDH-050 | 616.8 ± 3.2 | 0.03 ± 0.05 |
| P20000 | ALDH-051 | 47.2 ± 1.0 | ND |
| A0A6H1TS81 | ALDH-053 | 510.6 ± 47.2 | 0.41 ± 0.07 |
| A0A7L1RJ35 +A0A7L1RY59[†] | ALDH-054 | 65.1 ± 2.8 | 0.61 ± 0.09 |

"ND" = activity not detected. [‡]ALDH-044 was isolated from *E. coli* BL21 genomic DNA, and this specific sequence does not have a readily searchable accession number. See Supplementary Table 6 for the appropriate amino acid sequence. [†]To clone a complete ALDH-054 from fragment sequences, A0A7L1RJ35 (residues 1-112) was concatenated to A0A7L1RY59 (residues 1-339).

**Supplementary Table 2:** Round 2 ALDH Activity Screening.

| Uniprot ID | ALDH # | Hexanal Oxidation Activity (nmol/mg/min) | Hexanoate Reduction Activity (mg hexanol/L) |
|---|---|---|---|
| A0A6N7YK66 | ALDH-055 | 6.4 ± 0.1 | 0.17 ± 0.05 |
| A0A2T0SSF3 | ALDH-056 | 3.0 ± 1.1 | ND |
| A0A918MZJ5 | ALDH-057 | 23.0 ± 5.2 | 4.25 ± 0.60 |
| A0A522WLN6 | ALDH-058 | 13.7 ± 0.1 | ND |
| A0A858QBL4 | ALDH-059 | 7.3 ± 1.0 | 0.26 ± 0.02 |
| A0A1G9VJH9 | ALDH-060 | 283.0 ± 30.1 | 2.67 ± 1.32 |
| A0A1B1YTA3 | ALDH-061 | 278.4 ± 13.2 | 0.06 ± 0.05 |
| R4G109 | ALDH-062 | 12.8 ± 0.1 | 0.02 ± 0.03 |
| A0A261U014 | ALDH-063 | 194.8 ± 20.3 | ND |
| I3XUN2 | ALDH-064 | 5.9 ± 0.1 | ND |
| N8Z6I5 | ALDH-065 | 537.7 ± 8.5 | ND |
| A0A850GCI6 | ALDH-066 | 96.6 ± 5.7 | ND |
| UPI00234FF82E | ALDH-067 | 97.9 ± 5.1 | 0.08 ± 0.08 |
| A0A2U3D9B3 | ALDH-068 | 10.9 ± 0.9 | 1.32 ± 0.13 |
| A0A1W9L4H0 | ALDH-069 | 13.1 ± 0.2 | 2.14 ± 0.13 |
| A0A3S0VUQ7 | ALDH-070 | 8.2 ± 1.5 | 0.24 ± 0.03 |
| A0A177P8X5 | ALDH-071 | 173.8 ± 13.5 | 3.21 ± 0.20 |
| A0A7Z0SSR0 | ALDH-072 | 206.5 ± 0.6 | 1.65 ± 0.06 |
| UPI000423BFB2 | ALDH-073 | 142.7 ± 4.1 | ND |
| A0A543HTC7 | ALDH-074 | 5.4 ± 0.6 | ND |
| A0A4V3IHV4 | ALDH-075 | 4.3 ± 0.3 | ND |
| A0A2X4UZ81 | ALDH-076 | 281.3 ± 18.3 | 0.04 ± 0.07 |
| A0A2S0MQS8 | ALDH-077 | 22.1 ± 0.0 | ND |
| A0A849HPN9 | ALDH-078 | 174.1 ± 85.9 | 0.28 ± 0.04 |
| A0A4R6PRX6 | ALDH-079 | 5.7 ± 3.7 | 4.78 ± 0.20 |
| A0A251X6G7 | ALDH-080 | 13.3 ± 0.9 | 0.72 ± 0.05 |
| UPI0005427CB8 | ALDH-081 | 1.6 ± 0.6 | ND |
| A0A395JK05 | ALDH-082 | ND | ND |
| A0A2E2EWH7 | ALDH-083 | 115.2 ± 3.9 | 0.26 ± 0.03 |
| Q9ZA11 | ALDH-084 | 6.7 ± 1.5 | ND |
| A0A3B0T5Y3 | ALDH-085 | 15.2 ± 1.5 | 1.51 ± 0.14 |
| A0A3B1AP81 | ALDH-086 | 5.5 ± 2.3 | 2.11 ± 0.04 |
| A0A443K8I0 | ALDH-087 | 298.2 ± 1.7 | 8.72 ± 0.77 |
| A0A4P6V3E3 | ALDH-088 | 46.8 ± 0.1 | 1.36 ± 0.16 |
| A0A7Y9PEX6 | ALDH-089 | 118.2 ± 10.4 | 0.22 ± 0.02 |
| A0A023CJ10 | ALDH-090 | 64.1 ± 0.4 | ND |
| A0A0J0V889 | ALDH-091 | 14.7 ± 1.1 | 0.02 ± 0.04 |
| A0A0K6GMR6 | ALDH-092 | 53.5 ± 2.0 | 2.06 ± 0.28 |
| A0A142D425 | ALDH-093 | 31.1 ± 2.6 | 0.74 ± 0.08 |
| UPI0007A942CE | ALDH-094 | 34.6 ± 0.8 | 2.84 ± 0.22 |
| A0A178T558 | ALDH-095 | 14.1 ± 0.5 | 5.25 ± 0.36 |

| | | | |
|---|---|---|---|
| A0A1I0TE05 | ALDH-096 | 40.9 ± 3.5 | 1.65 ± 0.51 |
| UPI0009ADBF94 | ALDH-097 | 35.5 ± 2.2 | 0.45 ± 0.51 |
| A0A1V9B5J1 | ALDH-098 | 30.4 ± 1.6 | ND |
| UPI00017E6B4A | ALDH-099 | 40.7 ± 0.9 | 0.31 ± 0.01 |
| UPI000A26C654 | ALDH-100 | 48.1 ± 2.1 | 1.08 ± 0.05 |
| A0A2M9T1Z9 | ALDH-101 | 19.5 ± 0.0 | 0.06 ± 0.05 |
| A0A327YHK1 | ALDH-102 | 35.3 ± 2.7 | 3.12 ± 0.25 |
| A0A4Q1RV81 | ALDH-103 | 41.9 ± 1.9 | 6.66 ± 0.24 |
| A0A4R1QFD0 | ALDH-104 | 57.5 ± 2.8 | 2.74 ± 0.21 |
| A0A6G9J022 | ALDH-105 | 44.1 ± 2.0 | 4.90 ± 0.41 |
| UPI00050099C5 | ALDH-106 | 2.3 ± 2.6 | 3.09 ± 0.27 |
| A0A7U3YC36 | ALDH-107 | 31.4 ± 4.0 | 10.39 ± 0.36 |
| A0A7V9YZI5 | ALDH-108 | 35.0 ± 0.7 | 2.72 ± 0.23 |
| A0A7W0BY09 | ALDH-109 | 77.9 ± 3.5 | 9.51 ± 1.15 |
| A0A7W8JJ24 | ALDH-110 | 17.6 ± 0.2 | 2.38 ± 0.12 |
| A0A7W9YP99 | ALDH-111 | 22.0 ± 0.9 | 2.29 ± 0.38 |
| A0A7W9YPV4 | ALDH-112 | 56.4 ± 1.8 | 5.63 ± 0.25 |
| A0A840DPH9 | ALDH-113 | 26.6 ± 1.2 | 2.10 ± 0.12 |
| UPI001363667A | ALDH-114 | 31.9 ± 1.8 | 3.20 ± 0.24 |
| C5D8G6 | ALDH-115 | 15.5 ± 1.1 | 3.20 ± 0.25 |
| UPI0002BF8D7A | ALDH-116 | 32.6 ± 1.8 | 6.14 ± 0.26 |
| M8CWA2 | ALDH-117 | 21.1 ± 1.3 | 1.91 ± 0.47 |
| S5ZGT3 | ALDH-118 | 8.4 ± 1.5 | 3.61 ± 0.47 |
| S7SR93 | ALDH-119 | 88.8 ± 7.1 | 0.34 ± 0.06 |
| U2WRT1 | ALDH-120 | 81.9 ± 0.2 | 3.54 ± 2.24 |
| UPI0004DFAE51 | ALDH-121 | ND | 4.06 ± 0.36 |
| UPI001EEBB712 | ALDH-122 | 26.0 ± 2.0 | 5.19 ± 0.40 |
| UPI0015843A2B | ALDH-123 | 7.1 ± 0.5 | 9.52 ± 0.31 |
| UPI0014923DAD | ALDH-124 | 40.2 ± 0.0 | 3.05 ± 0.58 |
| UPI00030EE752 | ALDH-125 | 29.6 ± 3.1 | 2.37 ± 0.51 |
| UPI0005CD4570 | ALDH-126 | 40.0 ± 0.3 | 10.76 ± 0.56 |
| UPI00228641DC | ALDH-127 | 58.3 ± 0.3 | 3.79 ± 0.10 |
| UPI00135864F6 | ALDH-128 | 67.3 ± 0.1 | 5.34 ± 0.40 |
| UPI00208DAAE2 | ALDH-129 | 72.2 ± 0.9 | 5.45 ± 1.15 |
| UPI001315BED2 | ALDH-130 | 67.7 ± 9.2 | 7.09 ± 0.25 |
| UPI000489A7CA | ALDH-131 | 5.4 ± 0.1 | 1.46 ± 0.08 |
| UPI001FCBF132 | ALDH-132 | 32.5 ± 1.7 | 24.67 ± 4.58 |
| UPI0001D589C3 | ALDH-133 | 121.4 ± 7.2 | 3.35 ± 0.21 |
| M2XRT2 | ALDH-134 | 17.9 ± 0.7 | 0.12 ± 0.09 |

"ND" = activity not detected.

**Supplementary Table 3:** Characteristics of Primary Sludge from Wastewater Treatment Plant

| Parameter | Primary Sludge |
|---|---|
| Total solids (TS; g/L) | 35.9 ± 1.8 |
| Volatile solids (VS; g/L) | 30 ± 1.65 |
| pH | 5.59 ± 0.01 |
| Total Chemical Oxygen Demand (TCOD; mg/L) | 35100 ± 8833 |
| Soluble Chemical Oxygen Demand (SCOD; mg/L) | 4780 ± 311 |

**Supplementary Table 4:** Carboxylic Acid Concentrations in Waste Stream Upgrading Reactions

| | Wastewater Anerobic Digestion Effluent | | Food Waste Anerobic Digestion Effluent |
|---|---|---|---|
| Substrate | Primary Fermentation | Electrodialyzed Primary Fermentation | |
| Acetic Acid | 15.9 mM | 46.35 mM | 10.6 mM |
| Propionic Acid | 14.65 mM | 45.6 mM | 1.4 mM |
| Isobutyric Acid | 1.2 mM | 2.4 mM | - |
| Butyric Acid | 6.4 mM | 8.1 mM | 5.3 mM |
| Isovaleric Acid | 1.95 mM | 3.2 mM | 0.9 mM |
| Valeric Acid | 3.05 mM | 4.5 mM | 1.1 mM |
| Isocaproic Acid | - | 0.4 mM | - |
| Caproic Acid | 0.5 mM | 0.4 mM | 23.9 mM |

**Supplementary Table 5:** Strains and Plasmids Used in this Study.

| Strains | Description | Reference |
|---|---|---|
| XL-1 blue | *E. coli* cloning strain | Stratagene |
| BL21 (DE3) | *E. coli* protein expression strain | Invitrogen |
| **Plasmids** | **Description** | **Reference** |
| pQElac | Empty vector used in recombinant protein expression and purification. $P_{LlacO1}$:: empty, ColE1 *ori*, $Amp^R$, N-terminal 6x His-tag | [3] |
| pLZ313 | pQElac gap *Pst* TS PTDH, ColEI *ori*, $Amp^R$ | [4] |
| pSAP179 | $P_{LlacO1}$::*T. chinensis adh*, ColE1 *ori*, $Amp^R$, N-terminal 6x His-tag | This work |

**Supplementary Table 6:** Amino Acid Sequences of Bioprospected Aldehyde Dehydrogenases

| Uniprot ID | ALDH # | Sequence |
|---|---|---|
| P47771 | ALDH-001 | MPTLYTDIEIPQLKISLKQPLGLFINNEFCPSSDGKTIETVN<br>PATGEPITSFQAANEKDVDKAVKAARAAFDNVWSKTSSEQRG<br>IYLSNLLKLIEEEQDTLAALETLDAGKPYHSNAKGDLAQILQ<br>LTRYFAGSADKFDKGATIPLTFNKFAYTLKVPFGVVAQIVPW<br>NYPLAMACWKLQGALAAGNTVIIKPAENTSLSLLYFATLIKK<br>AGFPPGVVNIVPGYGSLVGQALASHMDIDKISFTGSTKVGGF<br>VLEASGQSNLKDVTLECGGKSPALVFEDADLDKAIDWIAAGI<br>FYNSGQNCTANSRVYVQSSIYDKFVEKFKETAKKEWDVAGKF<br>DPFDEKCIVGPVISSTQYDRIKSYIERGKREEKLDMFQTSEF<br>PIGGAKGYFIPPTIFTDVPQTSKLLQDEIFGPVVVVSKFTNY<br>DDALKLANDTCYGLASAVFTKDVKKAHMFARDIKAGTVWINS<br>SNDEDVTVPFGGFKMSGIGRELGQSGVDTYLQTKAVHINLSL<br>DN* |
| P54114 | ALDH-002 | MPTLYTDIEIPQLKISLKQPLGLFINNEFCPSSDGKTIETVN<br>PATGEPITSFQAANEKDVDKAVKAARAAFDNVWSKTSSEQRG<br>IYLSNLLKLIEEEQDTLAALETLDAGKPFHSNAKQDLAQIIE<br>LTRYYAGAVDKFNMGETIPLTFNKFAYTLKVPFGVVAQIVPW<br>NYPLAMACRKMQGALAAGNTVIIKPAENTSLSLLYFATLIKK<br>AGFPPGVVNVIPGYGSVVGKALGTHMDIDKISFTGSTKVGGS<br>VLEASGQSNLKDITLECGGKSPALVFEDADLDKAIEWVANGI<br>FFNSGQICTANSRVYVQSSIYDKFVEKFKETAKKEWDVAGKF<br>DPFDEKCIVGPVISSTQYDRIKSYIERGKKEEKLDMFQTSEF<br>PIGGAKGYFIPPTIFTDVPETSKLLRDEIFGPVVVVSKFTNY<br>DDALKLANDTCYGLASAVFTKDVKKAHMFARDIKAGTVWINQ<br>TNQEEAKVPFGGFKMSGIGRESGDTGVDNYLQIKSVHVDLSL<br>DK* |
| P46367 | ALDH-003 | MFSRSTLCLKTSASSIGRLQLRYFSHLPMTVPIKLPNGLEYE<br>QPTGLFINNKFVPSKQNKTFEVINPSTEEEICHIYEGREDDV<br>EEAVQAADRAFSNGSWNGIDPIDRGKALYRLAELIEQDKDVI<br>ASIETLDNGKAISSSRGDVDLVINYLKSSAGFADKIDGRMID<br>TGRTHFSYTKRQPLGVCGQIIPWNFPLLMWAWKIAPALVTGN<br>TVVLKTAESTPLSALYVSKYIPQAGIPPGVINIVSGFGKIVG<br>EAITNHPKIKKVAFTGSTATGRHIYQSAAAGLKKVTLELGGK<br>SPNIVFADAELKKAVQNIILGIYYNSGEVCCAGSRVYVEESI<br>YDKFIEEFKAASESIKVGDPFDESTFQGAQTSQMQLNKILKY<br>VDIGKNEGATLITGGERLGSKGYFIKPTVFGDVKEDMRIVKE<br>EIFGPVVTVTKFKSADEVINMANDSEYGLAAGIHTSNINTAL<br>KVADRVNAGTVWINTYNDFHHAVPFGGFNASGLGREMSVDAL<br>QNYLQVKAVRAKLDE* |
| P40047 | ALDH-004 | MLSRTRAAAPNSRIFTRSLLRLYSQAPLRVPITLPNGFTYEQ<br>PTGLFINGEFVASKQKKTFDVINPSNEEKITTVYKAMEDDVD<br>EAVAAAKKAFETKWSIVEPEVRAKALFNLADLVEKHQETLAA<br>IESMDNGKSLFCARGDVALVSKYLRSCGGWADKIYGNVIDTG<br>KNHFTYSIKEPLGVCGQIIPWNFPLLMWSWKIGPALATGNTV<br>VLKPAETTPLSALFASQLCQEAGIPAGVVNILPGSGRVVGER<br>LSAHPDVKKIAFTGSTATGRHIMKVAADTVKKVTLELGGKSP<br>NIVFADADLDKAVKNIAFGIFYNSGEVCCAGSRIYIQDTVYE<br>EVLQKLKDYTESLKVGDPFDEEVFQGAQTSDKQLHKILDYVD |

| | | |
|---|---|---|
| | | VAKSEGARLVTGGARHGSKGYFVKPTVFADVKEDMRIVKEEV<br>FGPIVTVSKFSTVDEVIAMANDSQYGLAAGIHTNDINKAVDV<br>SKRVKAGTVWINTYNNFHQNVPFGGFGQSGIGREMGEAALSN<br>YTQTKSVRIAIDKPIR* |
| P54115 | ALDH-005 | MTKLHFDTAEPVKITLPNGLTYEQPTGLFINNKFMKAQDGKT<br>YPVEDPSTENTVCEVSSATTEDVEYAIECADRAFHDTEWATQ<br>DPRERGRLLSKLADELESQIDLVSSIEALDNGKTLALARGDV<br>TIAINCLRDAAAYADKVNGRTINTGDGYMNFTTLEPIGVCGQ<br>IIPWNFPIMMLAWKIAPALAMGNVCILKPAAVTPLNALYFAS<br>LCKKVGIPAGVVNIVPGPGRTVGAALTNDPRIRKLAFTGSTE<br>VGKSVAVDSSESNLKKITLELGGKSAHLVFDDANIKKTLPNL<br>VNGIFKNAGQICSSGSRIYVQEGIYDELLAAFKAYLETEIKV<br>GNPFDKANFQGAITNRQQFDTIMNYIDIGKKEGAKILTGGEK<br>VGDKGYFIRPTVFYDVNEDMRIVKEEIFGPVVTVAKFKTLEE<br>GVEMANSSEFGLGSGIETESLSTGLKVAKMLKAGTVWINTYN<br>DFDSRVPFGGVKQSGYGREMGEEVYHAYTEVKAVRIKL* |
| Q04458 | ALDH-006 | MSNDGSKILNYTPVSKIDEIVEISRNFFFEKQLKLSHENNPR<br>KKDLEFRQLQLKKLYYAVKDHEEELIDAMYKDFHRNKIESVL<br>NETTKLMNDILHLIEILPKLIKPRRVSDSSPPFMFGKTIVEK<br>ISRGSVLIIAPFNFPLLLAFAPLAAALAAGNTIVLKPSELTP<br>HTAVVMENLLTTAGFPDGLIQVVQGAIDETTRLLDCGKFDLI<br>FYTGSPRVGSIVAEKAAKSLTPCVLELGGKSPTFITENFKAS<br>NIKIALKRIFFGAFGNSGQICVSPDYLLVHKSIYPKVIKECE<br>SVLNEFYPSFDEQTDFTRMIHEPAYKKAVASINSTNGSKIVP<br>SKISINSDTEDLCLVPPTIVYNIGWDDPLMKQENFAPVLPII<br>EYEDLDETINKIIEEHDTPLVQYIFSDSQTEINRILTRLRSG<br>DCVVGDTVIHVGITDAPFGGIGTSGYGNYGGYYGFNTFSHER<br>TIFKQPYWNDFTLFMRYPPNSAQKEKLVRFAMERKPWFDRNG<br>NNKWGLRQYFSLSAAVILISTIYAHCSS* |
| A0A6P3Q7G7 | ALDH-007 | MLRAAALAAAGLGPRLGRRLLSGAATHGVPAPNQQPEVFYNQ<br>IFINNEWHDAVSKKTFPTVNPSTGEVICQVAEGDKEDVDRAV<br>KAARAAFKLGSPWRRMDASDRGQLLNRLADLIERDRTYLAAL<br>ETLDNGKPYIISYLMDLNKVIKCLRYYAGWADKHHGKTIPID<br>GDFFSYTRLEPLGVCGQIIPVTMTLLVQLLSSSSWELRGVGQ<br>CSKEWPDLVKFSRSPVVRLLESSSCGVLCSVLGAFGPFSSPE<br>SLSFPASLQVGQLIQVAAGSSNLKRVTLELGGKSPNIIMSDA<br>DMDWAVEQAHFALFFNQGQCCTAGSRTFVQEDVYAEFVERSV<br>TRAKSRVVGNPFDSRTEQGPQVDETQFKKILGYIKSGKEEGA<br>KLLCGGGAAADRGYFIQPTVFGDVQDSMTIAKEEIFGPVMQI<br>LKFKTIEEVVERANSSKYGLAAAVFTKDLDKANYVSHALQAG<br>TVWVNCYDVFEAQSPFGGYKVSGNGRELGEDSLQAYTEVKTV<br>TIKVPQKNS* |
| A0A5J5DF59 | ALDH-008 | MLRAVLSRGLPRLSRPSVCHYSAAAVPAPSGQPEVLFTKLFI<br>NNQWQDAASGKTFPTINPATGEVICQVAEADGADVDKAVKAA<br>RDAFRLGSPWRRMDASHRGLLLNRLADAIERDSAYLAELETL<br>DNGKPYAVSYMVDLPNVVKCFRYYAGWADKWEGKTIPIDGDY<br>FCYTRHEPIGVCGQIIPAWKLGPALATGNTVVMKVAEQTPLT<br>ALYVASLIKEVGFPEGVVNIVPGMGPSAGAAIASHMDVDKLA<br>FTGSTEVGHLIQKASGSSNLKKVTLELGGKSPNIVLSDADME<br>DAVEQSHFALFFNQGQCCCAGSRTFVQADIYDEFLERSAERA<br>RRRVVDEEQFNKILGYISSGKREGAKLMCGGGVAADRGYFIQ |

| | | |
|---|---|---|
| | | PTVFGDVQDNMTIAREEIFGPVMQILKFKTLEEVVTRANDSK<br>YGLAAAVFTKDIDKAHYVSNALRAGTVWINCYDVFGAQAPFG<br>GYKASGNGRELGEYGLDNYTEVKTVTIKVPQKNS* |
| A0A0D6KE96 | ALDH-009 | MTTQLTPPQSQTTIRTYQNFINGKYISPLSGVNYERKSPLTG<br>ETIVQIPWSNQADTDVAIQAARQVFDDGTWSTSHARVRHDIL<br>RKTAELLTTKTSEIAAVICQEVGRPIGMCIGEVQMTAQVFDY<br>FAALTLNLQGESTTQYDRNAIGLTVHEPVGVVGIITPWNFPL<br>LLVAWKIAPAIAAGCTMVVKPSEFTPSTAFMLAEILSEAGLP<br>DGVINIVTGDGPVVGEHLVESPLVDKIAFTGSTAVGRRIMAK<br>GAPTLKRMSLELGGKSPNIVFGDADLSQAIPGALFGIYINSG<br>QVCQAGSRLLLHESIKDVFIEQFLAATQTFQIGNPTDNTTMM<br>GPVINEIQFERIQNYIQLGEKEGAKLLIGGSGRYLVPGFEQG<br>LFIKPTVFDHVTNEMAIAQEEIFGPVLSIMTFKDKAEALQIA<br>NQTMYGLTAAIWTKNLDTAFKMAKGIRAGTVWVNSYHTSGLE<br>PTMPYGGYKQSGIGREVGKNGLEEYLETKAIHIKLA* |
| L8N0N6 | ALDH-010 | METLPINNLDIPATIAKQRIFFDGNKTKDYEFRVSQLKKLAQ<br>LIKENEKLILDAVYADLRKPTIEIFGSEILIALSEIRYVIKH<br>LKAWMKPQKVGTPLNLFPSSSYIYTEPLGVVLIVAPWNYPFS<br>LNIQPLIGAIAAGNCAILKPSEYAPHTSNAIAKIINEHFDPN<br>FITVIEGGLEINQALLAEKFDHIFFTGSTAIGKIVMEAAAKH<br>LTPVTLELGGKSPCIVDEECDLETTAKRIIWGKFYNAGQTCV<br>APDYLLVSKSIKPVLIEKLLGYVKTFFGENPQQSPDFARIVN<br>DRQFDRLVSLLNEGKILIGGQTDKSDRYIAPTIIDGISIHAK<br>IMGEEIFGPILPVLEYDQLSEAIALIKSQSQPLALYLFSNNK<br>QKQEKILQEISFGGGCFNDTILHLANLELPFGGVGNSGMGSY<br>HGKATFDRFSHRKSVLKNSFRFDLKLRYPPYRVSIDTLKKFI<br>N* |
| A0A2V7VB71 | ALDH-011 | LKAMAGERLDIPLVIGGKEVRTGDKAKAVMPHDHRHVLGDWH<br>KASREHVAQAIDAAAKAHGEWSRWPWEDRAAVFLKAADLLAT<br>TWRATLNAATMLGQSKTVFQAEIDSACELVDFWRFNPAYAQE<br>LYAEQPLSSAGMWNQSEYRPLEGFVYAITPFNFTAIGGNLPT<br>SPALMGNTVVWKPASAAIPSGYWIMKLLEAAGLPPGVVNFVP<br>GDAVTVSDTVLTHRDLAGVHFTGSTEVFNSMWKTIGGSMSRY<br>RSYPRIVGETGGKDFIVAHPSAEPQALAVAIARGGFEYQGQK<br>CSAASRVYVPRSLWSDVRDRTVAIIKEIRVGDVTDFRNFMGA<br>VIDKKAFDKISEYIGDARKNATIVAGGGANGETGYFIEPTLV<br>EAREPGYRLLCEEIFGPVVTVYVYPDEKWEETLAVVDQTSAY<br>ALTGAVFATDRGAVRQAASALRHAAGNFYVNDKPTGAVVGQQ<br>PFGGSRGSGTNDKAGSKLNLVRWVSARSIKETFNPPRDYRYP<br>FMAEE* |
| UPI000407F0B2 | ALDH-012 | MNIPLEPLAATTDRRIEAIALGRTYTTQRRAIVHDLLSIPVA<br>EMTVAPPIYIHKAISSMREAPSLSLAEIHAAMARAADSYQYD<br>TIAGLSPDEYSRLLQRTTGLPETVTQNALATVADALRNMPDI<br>INAGRPQGARWSWDDADALAGYSLFSRKGDVFAVLAAGNGPG<br>IHALWPQAVALGYRTLVRPSTREPFTAQRVICAMVQAGLEHY<br>VALIPTDYRGADELVASADLALAYGGQDIVDKYRHHPQVRVQ<br>GPGRAKILIGADVCMEEAVSLVATSMTDLGGAACVSASAVMV<br>EGDVSDFCRRLRQTLQQLPEKVLPRASQQTVDWLNSVIDTPI<br>EASLMSEGYLLRPVVTEVAEPDDPRIHRELPFPCVTVAPYHP<br>TRSAQILSGSLVVTVFSRQPQLLKSIIADASISNVYVGDIPT<br>TWMSPLVPHDAYLSDFLMCNRGFRIAASWIEAESKGEKL* |

| | | |
|---|---|---|
| A0A2Z4LU87 | ALDH-013 | MFKFNISGRNFIGYNRSSKGDTTFQAKNPSTGEGLITSYYEA<br>TLDEVNQAIELAEKAFTGYREKTGQEKASFLEAIAEELAQLE<br>EGLVGLCVQETGLPEGRLKGELGRTMGQLRLFASVLREGSWV<br>DARIDFAVPDRKPSPRPDIRYMQKSLGPVGIFGASNFPLAFS<br>VAGGDTASALAAGCSIVVKAHPSHPGTSELVAMAINEAAKKS<br>NMPDGIFSMLHGVSNTVGEAIVQHPLIKAIGFTGSFKGGKAL<br>YDKAVRRKEPIPVYAEMGSVNPIFILSNALKEQYKTIAKGLS<br>DSVQMGVGQFCTNPGITIVPNINETSLFKEELNNCISNSESA<br>TMLSASIQEGYETGLKRLKSNEIITSLSKGEQKEGHNQGVPE<br>ILSVSAKDFLSENTLEEEVFGPSTLLVHAQDGEEMLKIAKSL<br>HGHLTATIHGTEEDLQANVDLLKILERKVGRLLINGFPTGVE<br>VCHSMVHGGPFPATTDSRMTSVGTAAITRFTRPICFQNFPDV<br>LLPDELKDGNPLKILRMENGKYMKAD* |
| A0A0S8BPB2 | ALDH-014 | MQGSAAIPEAPHRIEPTSQSKLDDALAILDDHKQEWAALDLE<br>ERIELLEQMREGVVEVAEDWVRASVEAKGMTFGTAEEGEEWM<br>EGPATTLRNIMLLIAALRDIEEYGVPQLPKPAFTRPDGQVVA<br>PVLPASTWDKLLFQGFTAEVWMQPGVTLENMSESQAAFYRDK<br>APKGRVALVLGAGNVASIGPMDALYKLFVEGQVVILKMNPVN<br>EYLGPFIDHAFAALRERGFFRVVYGGAAEGDYLCRHERVEEI<br>HITGSDKTHDAIVFGFGEEGSRRKAEAEPRNTKRLTCELGNV<br>SPVIIVPGPWSQKDLDFHGVNLATSVVNNAGFNCSATRVIIQ<br>HEQWGKREALLASLRKAFQQAEDRRPYYPGSEERQQLFLEHH<br>PHAEEFGTKGEGHVPWTLIHHLDPSDPDEICFNTESWCGQTS<br>EVALPADSVAEYIDRAVEFCNERVWGTLNCSIFVHPKSMKDP<br>QIAAAVDRAIASLRYGAIAVNHWAGLNFALVTPTWGAFPGHT<br>TEDIRSGRGVVHNTYMFDRPQKSVVRGPFRVFPKPAWFIDHK<br>TAAEVGRKMTYFNADPSLARLPGLIWSSLFG* |
| UPI000E14A9E8 | ALDH-015 | MTPVINPAEPKDIVGYVREATHAEVEQALQNAANNAPIWFAT<br>PPQERAAILHRAAVLMEGQMQQLIGILVREAGKTFSNAIAEV<br>REAVDFLHYYAGQVRDDFDNETHRPLGPVVCISPWNFPLAIF<br>TGQIAAALAAGNSVLAKPAEQTPLIAAQGIAILLEAGVPPGV<br>VQLLPGQGETVGAQLTSDERVRGVMFTGSTEVATLLQRNIAT<br>RLDAQGRPIPLIAETGGMNAMIVDSSALTEQVVVDVLASAFD<br>SAGQRCSALRVLCLQDDVADHTLKMLRGAMAECRMGNPGRLT<br>TDIGPVIDKEAKTNIERHIQAMRAKGRPVFQAVRDNSDDARE<br>WQTGTFIAPTLIELESFDELQKEVFGPVLHVVRYTRNNLGSL<br>IEQINASGYGLTLGVHTRIDETIAQVTGSAHVGNLYVNRNMV<br>GAVVGVQPFGGEGLSGTGPKAGGPLYLYRLLANRPENALGIT<br>LARQDADYPVDAQLKAALVQPLEALREWATDRPSLHALCQQF<br>GELAQAGTQRLLPGPTGERNTWTLLPRERVLCIADDEQDALV<br>QVAAVASVGSLILWPDDAFHRELAKRLPAAVSGRIQFAKADN<br>HHRAAV* |
| A0A7Y1V182 | ALDH-016 | MPEVFQTISPVDGRVYVERSYAQLQDIDHTLNQAQQIQMEWQ<br>ATSIDERAQICRKAVHYLVAHSNKLAQELTWQMGRPIRYTPN<br>EILGGLQERAIHMIDIAASALAEESLEDGNQFKKVIKHGALG<br>TILVLAPWNYPYLTSINAIIPALMAGNTVILKHSQQTPLCAE<br>SYADAFNNAGLPAGVFQYLHMTHTQVAKVVVDSRIDFVSFTG<br>SVEGGYAIQEAVGKRFMRTGLELGGKDPAYVRADADLSYCSE<br>QIADGAFFNSGQSCCGVERIYVHTEVYDQFLEAFLSATTQLN<br>LDDPTKADTTLGPMIKPTAAAFVTDQIDAALAMGARPLVDVR<br>SFPNHEVKRGYMAPQVLVDVDHRMSIMQDETFGPAVGIMKVK |

| | | |
|---|---|---|
| | | DDAEAVHLMNDSRYGLTASIWTQDLERGEVLGKSIQTGTLFI NRCDYLDPSLAWSGVKDSGIGISLSHLGYLQVTRPKSFHIKK * |
| A0A517R638 | ALDH-017 | MMLWKPRPLLPDMIFSDRTVAATVSMDRPNMSTITAQRPNID RRSHSDDPDEVAFVSEALTQARSSQSAWAEQPLREKLKIVRR FREQIAARPDDFTYAVELPQRRNRAETLASELLPLADACQFL ESEAPRLLEPKKLSKRGRPGWLKGVIAEEHREPFGVVLIIAT WNYPLLLPGVQMLQALVAGNAVLLKPGKGSHAAAVALREAVV ACGLDPNLVTVLSEATSAAQTAITPPEGTSGADKVILTGAAA TGRKVLAQTAEHLTPAAMELSGCDAVYVRGDADLDLVVDCLA LGMTFNGSATCIAPRRVLVHNSICDELADRLSARFAELPSAA IEPTLAGRLSRLVDEACDDGATKLAGQVEGHVVAPILLRDAS PEMALLQADIFAPLLSIVRVTGDEQALEFDRRCPYALGATIF STDETAARSLAAQINAGCVVINDFLIPTADPRIAFGGRGESG FGVTRGGQGLIEMTQPKTVVVQRASWRPHLTPPDETYEQFFK NYIASAHAKSPWARAKAGFAFLSEAVKRQREQR* |
| A0A1F8MB45 | ALDH-018 | MNRVRTATTDDIFPWQPDEISGAKPAHLMNFIDGKWVKANKY REVVDPVSGEAFIQMPDTSEKELAPFIQSLAKCPKYGLHNPI LNGERYLMYGQISERLADFLRSSEGDSYFTRLIQRVMPKDTA QCRGEVSVAATFLENFSGDQVRFLAKGQTTPGDRLGHEAIDY RWPHGPVAVITPFNFPLEVMTLQTMGALYMGNKPVLKQATTT SIVAEAFVRLLLHCGMPAQDLLLIHCGGQVMEKLVTDPVIQF TQFTGSTSVAERLLKLTRGRCRIEDAGFDWKIVGPNAIPSML DFVAAQCDMDAYAASGQKCSAQSLLAVHENWMKLGLLARTAE LSARRTMKDLTIGPVLTWTNEMIEEHINKLLRIKGAKIQFGG KRLKRHSIPSCYGSFEPTAVFVPLKSIASHVALVTTEVFGPL QVITTWETKDDLKLLLNICNSMEHHLTAGVVDNDPDFLNNVL GQTVNGTTYAGIRARTTGAPQWHFFGPCGHPAAGGIGTIEAI RNIWSLNRTIIKDTNVPSLGE* |
| A0A346XWA7 | ALDH-019 | MERLEKLTAGMPLLVGGDRFTTVPTDIADAFEPGDAVLVADT GEVLHVPDAERRAATVAVDAAVAAFDLMGTATDEQVSAFYDT FANALADDTIFVAIAEANAADVEAARQKGRSTTRLELTTRMR NDMVEGLRGWRDMPGGRGEVVETVTHDGWRVEQVRDRLGVVG FVFEGRPNVFADACGVLRSGNTVVFRIGSDALGTARAIVTHA LRPALAAAGLHEGSAALVDSPSRAAGYALFSDPRLSLAVARG SGRAVSMLGGIARKAGIPASLHGTGGAWIVAGPTADADDFGE AVTRSLDRKVCNTLNTCAITADRADELVPVFLDALRAAGTAR GVEPKLHVSADSVEHVPSDWFDRQVPIARAEGEVKEAQAERI AADELGREWEWEDSPEVTLVVVDSIDDAVQRFNAQAPRFVAS LLATDPAEHDRFFATIDAPFVGNGLTRWVDGQYALGKPELGL SNWETGRLFARGGILAGDSVHTVRTRAVLDRPDIPR* |
| A0A2E3KN37 | ALDH-020 | MMGGTGGRTRTDTSAVVYHRRPKPTAPTMTSHSRQLELLRAD RSRWAGRFRHLILKGTEEFVRLADSELGKPRHETITAELLPL IASCRWHQRQARRILKTRRLKGRPIWLFGQQHRIQHVPLGTV GIIATWNYPIQLLGIQMLQAALAGNDLIIKPSEHAPQCQAFL IELAHKAGLDERALRSLPATREAGAAMIERESLDHLVFTGST RVGRLVAEACASRLIPSTLELSGRDSALVLEDADVQLAARSI WTAVTSNAGQTCMAPRRVLVHQDRYQAFCAALTPLAEKAIPR RLTLPEDAARIDSQVDEAVRAGGRAIPHTEAQSDASAFLPRV VLDCPDGTPLMDGDHFGPALAVHACASLNEMLEHHQGVGQYL ATSIWTGNPTAARGLAAELRSATVTINDVLIPTAHPGASISG |

| | | |
|---|---|---|
| | | HGPSGWGTSRGAAGLLSMSRPVHVTTTPRKMRMPTDVPTDAA LAKLEKLIGVRRPSVDDDSKVTSNSPQPHSGKSS* |
| UPI000A40ADAD | ALDH-021 | MLGASLDKDQPDNPINYFHRIPAHAKRLTTYRTMMNIFTACH PEDLDTAYHYLLKHEKTRHDIINDDTFATVIEKAFAYYKDVI APAYRPRLLDTQSAALLHSFSAQLKRLDVKNKFITCLLLNPI VDSVGFRCQRDAYKTIYACLFGDEHGPRLAEYINFLGVQAFC EKLDSRIQQAAHDREYDPMKEDTHASQEALQMSDAANHTAKS ETLNTSDQAPEAIDFDEIKARAEAYALHLRSHSESIAESLSG FECYNVAVDEIERCIEFLENIELNRSFFERRVNCVTSYLPLN QPIYATTCFGIIPSLLARSAWLRPPTAMHPHYKKLLNPLKLD HFFPNLHVSFADKDTFVSQTAAISDAVIFTGTPENAAKVRKS YLKRTLFILNGAGHNPLVVAADACITTAVESALRVVLYNQGQ DCAGPNSIMVHVEAYPEFIRQLRKELTRCEGLVGDYKCKKNI VGPNSDPDHTLKVMKMFRDCREHCTYGGEINPVSGLIRPTIF ERPLSLGGNYKEFFAPVFFVQLYSDDAELASYFEHPLYSPNS MYISLFGSSDYISGLIEQGKHDPCTVLSNTDLHIVEKGYAPY GGQGVAASCLYVNGVRIAKPTLPQRDIHEHLVAHPHH* |
| A0A139NAP6 | ALDH-022 | MTQTAQEAFAQLDPQAWTKVSIEERLQILAEIQANMRQYGAE LGQAEMAMKNRLTGADLYSQTDGMLQTLVAVGNVINASTFIY QTLAETGKMPEAKSIRDLGDGTFEVEVFPTAPVDQMTAATQH GYLRLVGQPKQVSPLDKEAGIIAVSGAGNYSSSIETIKAIFF DNKTVIHKAHRLNEATDKVWEKIFAPLVERKALSFAGVDYSR DLIQLEGLDAIYFTGSTAVAKNIMASTDTPLVSECGGNNPAI IVPGDRPWTAEEIKNQAELIVSISKGNGGAACGRPQTFITSK QWVQREEFLDAIRQAAQSTFAVGTYYPKSADVREAFLAAHPQ AEIIKPEGGQYPNTDFLFIPDMDKSAYGVTHEAFCQIMGEVA LDVPATAEAFLPAATAFANDELLGTLGCMILIDDETRANHEA SFQTALSELNYGGITVNTTPPMVWFNAYLTWGGCKETKENFV SGIGNFGNALNFEQVEKSILVEQFAATGFLYNDRQATDAMNQ QVINFTLGNME* |
| A0A1X6ZLY8 | ALDH-023 | MTATPDPALEAALDAFGPAEPAPDRAARKAHLEALEREMRSH AEAAAEAVAGDFGARPRPETLLTEVAMVIGAAQHARRHLRRW MRPERVVLPPHLWPSTARVDRVPLGRVGIIGPWNYPVQLALV PLVAAIAGGNRAILHPSEHTPRSAALIARIVERAIPDRARVL TGGADQARALAAAPLDGLFFTGSTATGRHIMAAAAQNLVPVV LELGGKSPAILRHDADIDAAARSIMAGKLLNAGQTCVAPDYA MVPREMLDRFVAALKTATEALYPDPAGPDYAAIARASDRDRL AALLDGLDPVPLMARPPAPPRMGAVAVIDPDPDHPLMREEIF GPILPVIPYDAPHEPRDFVAARPCPLALYVYGRDLAAARSEA EAIPAGGAVINEAVLHVGVQQLPFGGAGASGLGAYHGAEGFR AFTRPRSTMIARPSLARLVRPPYGRNVERILKSLIG* |
| A0A060QGV9 | ALDH-024 | MSDITTSFTRNSHGPAESLEASASLHDECEDKSSSTSSQEVC RDLKAMQPRWAAVPLRARLRVLRRFNRLMLLNAPSLIRLIAD HASLDVMTAEILPLMAASRFLCREASSMLKETRLGWLGRPLW LQGVMASVVRKPLGAVLILAPGNYPLMLASIQTLQALVAGNS VALKPAPGRTAVLRRFVALLEQAGLPKGVVQLVGEDSGQQAV SSGYDLIIILTGSAETGRKVALAAAETLTPTIMELSGADPVFV LPDADLALVARALHFGQTLKGGHTCIAPRRIFINEVQKRPLQ RELYRVFGGEGDNPAPDPSSKLAQLICSAKAAGGEVVVCGKT QVIFLNAHQARLADIDLFAPWFAVITTHSVEEAICLEGAATH ALGASIFGNEREALAIAKRIPAGTITINDIIVPSADPRLPFG |

| | | |
|---|---|---|
| | | GAHRSGFGVTRGREGLLALTRPVSISTRKRGAFHLLPRLRQW HGR* |
| UPI0001E31496 | ALDH-025 | MSINPELAENTLFLKRHESILKCLNYLIENRQEVMDILTQFS SYRAANAEIDSTILTLQGALQEVQTYQPSWQRSMAVFMPSNV ILYSYALYLLIPSLYVENIDFRPSSHVNEYVNMLHEKLQAVH GLPIYIRKVSQRVFMENSVMPADIVVFTGSYVNAEKIKKQIR KDQLYIFYGQGINPFVIGPDADLELAVTDVIRMRLFNSGQDC LGPDIILVHQEVSEHFKDLLIQRLDQLVFGANDDPNANYSPI FYKDALNSVSEYFNTNDKFIIYGGGIDFRTKKMEPTVVYSEL DQNLEIIEYFSPVFNVVSYQDDEQLIKRISSSYFSERAMGCS LYGSEHLTDVLRKKHTLTINQTLEDVDQGNKPFGGYGTMSNY IFYDFKLISKPILISEIVAEYLSEKRSLV* |
| A0A537X7T4 | ALDH-026 | MRSYPLYIDGRDDPGQGWTYTVRTSAFIDDPAETFGLKRRLE LEGDRAFGDDEPTAIGRCAWGDDAENRRAVDAAARASREFGR FAPRARRQIAEDFHDTVIAASDEFVEILVAEGHPRRLAEWEI KGILANCCPPTLDWVFAQLHQEYEDAEGRRVRLTRKPDGVVC VNPPQNAAGVMSVMGVGALFAGNAVVIKAPRSTPLGVMFAYR EIVAPVLERHGAPPGTLNVVSGDTRRTLRQWLSHPGVDDVFF VGDSTVGLKLGAQCVARGKKPILELSGNDPLVVWRDADLEGA AEALCESFYGSAQICMVPKQAIVHPAIAAEFVELFLDRVATI RPGYPEDPATLLSPVLKADRFLEFLAEARDAGAELLTGGRRV GLDGAPSESGLFFEPSVVRVDGLEDAARLRCVREETFFPLVP IVVPEDAPDHLLLERVIEFVNADEYGLRTSLWATEPRVVEEF TAGVWSSGTLRVNESHMGFSPPMATHGGTGRSGGPFGGLNFP ALSTSHLQGISIAPVRAPLPARETEDAAGAVPAAA* |
| A0A2E8CRC4 | ALDH-027 | MIAPHIPILRQGQVYKSLDTQTVNKLGSDEPAAEISFACADM IKYDIQNMGSAREALKKFSGEELVEITKKAGELFLNGDLPIG TEGELQSPEDYVFSLAATSGLPHSLIRFNMRRLGGLFGQIDD IFKGLSRGLEWSVLDRGYGEQGGAPVSFSPTTNELAVILPSN SPAVNALWIPAIAMKVPVLLKPGREEPWTPWRIIQAFIKAGC PPEAFSLYPTQHDGSGAIIRRAGRVMLFGDDSTVKQYENDER VEVHGTGYSKFVIGEDEIENWESYIDSMVESVSANSGRSCIC TSTIVVPKYGDEIAHALSKKLCEIKPLPQDDPEAKLSGFANP KFAEWIDEAVEEGLQSDGARDITADYRDGARFVERDGMNYLQ PTIVRVDSFDEDLALREFLFPFASVVECPQSDVVEKTGYSLV MTAVTKDVEWINELFDHPEIERLNIGNVPTNRISWNQPHEGN LFDFLYTRRSFAFAESA* |
| A0A2V7BEQ0 | ALDH-028 | MEPLGVVVVTPPWNFPLSIPAGGVLAALAAGNAVVLKPAPEA VLVGWHLANCLWDAGIPREILQFLPCPDDEIGRGLVTDSRVG GVILTGSAETARLFLGWRPDLPLFAETSGKNAIIITALADRD QAIRDLVRSAFGHNGQKCSAASLAICEAEVYDDADFRRQLRD GAQSLAVGTAWEPTSRITPLTQAPGAALRRALTVLDEGEEWL LEPRPATDNPQLWSPGIKLGVRAGSFFHRTECFGPVLGLMRA ENLDHAIELANAQPFGLTSGIQTLDDREIARWVDRIEAGNLY VNRPITGAIVGRQPFGGWKASSVGPGAKAGGPNYVLQLARWR QVARPAVDQEPLSESLATVLDRCLAGLTDADARSLLEASAAS YARAWREHFSREHDPSAIRGELNAFRYRPCRHVIARGMTARP EAAVALCQIILAAHVAGTRLTVSLSPESEPWVGLAECAGVEL VVEAEAGFVDRLAHPLYREHGWIERLRAWEPISTAARAAANG TGVTVIDAPVLANGRLELRWYLREQTVSRVLHRYGSVTEPDA * |

| | | |
|---|---|---|
| A0A1T1H988 | ALDH-029 | MTASTSSSNTGQCFINGVWQAGEGSEFTSLNPATGEVIWQGK EASAAQVELAVNAAREASVEWAMMPFADREAIARRFAELLGD NKEEMATIIATETGKPVWETRTEVGAMVGKIAISVNAYNERT GSRVSDVAGARAVLRHKPHGVVAVFGPYNFPGHLPNGHIVPS LLAGNTVLLKPSELTPHVAEFMVQLWEKAGLPAGVLNLLQGQ KDTGIALAGHDRIDGLFFTGSSRTGHILHEQFAGHPGKILAL EMGGNNPLIIDEVADMKAAVHETIQSAYITSGQRCTCARRLF VPVGEWGDQFIAQLQEAVSRIKVGGQFDEDAPFMGSLISEAA ADGIAAAQDNLIGLGASPLVKLEKLKPGTGFLSPGLIDVTAL VEADKLPDDEYFGPLLQVIRFSDFDEAIRQANNTQYGLSAGL FSDSEARFNYFYDRIRAGIVNWNKQLTGAASSAPFGGVGASG NHRASAYYAADYCSYPVAGMEADHLVLPENLSPGLTIK* |
| A0A382Q6S7 | ALDH-030 | AVHINAFNFPIWGMLEKIAVNLMAGVPAIVKPATLTCYLTEL MVREIIATQILPEGSLQLICGSANGILDHVCCEDVVTFTGSA STGKMLKAHSKLIDEAVPFNMEADSLNASIIGEDAIPGTEEF DLYIKEIQKEMTVKAGQKCTAIRRIIVPEKLVEDVQKALCDR LSKTIIGDPAIEGVRMGSLAGNSQVVEVSEKVNELAESQDII YGDLENFDVVGANKNKGAFIPPILFLNDNPFEKTDCHIIEAF GPVSTILPYKNLDEAIELARMGKGSLVCSIVTSDDNIAREFT VNAASMHGRILVLNKDCAKESTGHGSPMPLLTHGGPGRAGGG * |
| M4Z1V7 | ALDH-031 | MHTLSASANEVTFIVQRARIAQKCFEGAQQREIDLAVAAAGW RCYRDETAQELSSLAIEETELGNAVDSYQRLRKRILGTLRDL ADAVTVGLVHEDVARGVRKFAKPVGVIAGITPATAPAAAAIV NSLSSLKTRNAIIFCPNPRAHRTVGRVVELVRDALREVGAPV DLVQCVGVPTRSISEELMATADLTIASGGASTVRRAYRSGKP ALGAGVGNAVVVVDDTADLDAAASMIIAGKSFDYGTSCSSES CILVDESVSDQLIGKLVENGAYMCSGPEMASLRRTAWPDGEL AREIVGKSAKQIADRACIEVPQKTRVLLTLPETTHAEPLGGE KLSPILALWKFAQFDHAVGLVQRLVAASGAGHSCAIHTNVRE RAEILARTINVSRVLVNQSTGMGNSGSFDNGLPFSVTLSCGT WGGGSTTDNVNWRHFLNYTWLSETIPKNEPRPEDLFSEYWSV YHPTASYVGQLGGM* |
| A0A420XXS9 | ALDH-032 | MAQGISIKSPVGDKLDWDCQELKAKFPKTKDDIKEPIPSPES CLLLIDGKVVKSEKTATEVESPVITQDDNKHVIIGKFEMASE AQAMQALEAAQRAYGYGREEWSKMKLEERCKHMEGFANDLEK KTDEIAQLLMWEIGKTSKDAKSEVTRTVAYIQTAIKEAKALS ESESKWEEEKGILCQIRHLPVGVVLASSPFNYPLNEAYTTFI PALLMGNSVIVRAPRNGATPHFPTLELFAKHFPAGTIQFLTG SGREVMGSLIGTGKIDAVAFIGTASSAAGLFKGAPNPQKLRV MYEGEAKDSAIILPDADLDVAVESCASGMTSFNGQRCTAIKM IWVHESKAEEFLKKLGEKIDSMQLGMPWEEGVKITPLCEDTK PGYIKELIEDALKKGAHIVNNGGKCYATLCTLSILAPANKDM KIWSEEQFGPVTPVAFYTDLQEPIDYVAKSEYGLQASVYGYD EDDIAKVVDALSYHVGRLNINAPDQRGPDVFPFTGRGNSALG ILNAPEGLKFFSVPTMVATKTDDERNVKVLKAVNAGGKSQVV FGAK* |
| A0A1F6LNY1 | ALDH-033 | MASEQEIHQVVGKVMEKVRQRYTVGTGGKIFSSAPGRATSPS MDFRPRKAVGGDSTFDDPDAAAKAARQAQRELMALGLEKRFE IVAAMRAAALENARRLGELAVSETKFGKMPDKMQKVELAARK TPGPEIIHPVTFTGDHGLTLVEKAPYGVMVSITPTTNPPSTV |

| | | |
|---|---|---|
| | | VNNSIGMVSAGNAVIINPHPNAKEVSCEAARILQLAIESAGG<br>PKHLVCAIGNPSQESGTKLMNHPGVDATLVTGGREIVRVAMS<br>SGKKVFAAGPGNPPVVVDETCVLPKAAKDTVFGAQFDNCALC<br>TGEKEIFCVASICDDFKAEMGKNGSVEIKGRDAERLTKLIIV<br>EDSGSGERHPRVNKELIAKDAVKILAQAGIRAPAGITHAFLE<br>VDWDHPLVMAEQLMPITPIVRCKDYDQAKEWAILAEHRFRHT<br>FTMHSTNIARLSDMARASNANIFVKNGPSLAGLGFGGEGYTT<br>LSIAGWTGEGFTTAATFTRERRCTLVDYFRIV* |
| A0A7H4GQ81 | ALDH-034 | MNFEPTGIDPLLDEALSAIATRSAFSGFKESGSSRIHGTEKP<br>AAGKAAYEACLGQAFRLDDSEADFIAAEEVSPFTGEALGIRY<br>AGPDPDRQFAAARAAMKGWASASVEQRIKLCVEMLRQVDEHG<br>FEIAHAIMHTAGQSFPMAYAGSGANALDRGLEAVAWSWLMMN<br>ALPFQASWQKQVGAGDPVRLAKRWRLMPRGPAVVFCCASFPT<br>WNGYPAMMANLACGNPVLVKPHPTAVLPMAIAVRIIRKVLAE<br>AGFDPGLIQLVVDSSSKPLGKVLVEHDQAAIIDFTGSARFGS<br>WVEQHAGNRPCYTETSGINTVILDGAEDIDAALDSVVNTLCL<br>FSAQMCTSPQTFYVPASGMSDGNGRCLSADEVEAALAGRIDA<br>LAANPKRAASVMGCIQSPATLSLIDEWTGLAESGRYRLLHRS<br>DHYAHPAGPDARTATPLVLGLDLEDGTAPQEEVFGPIAFVIR<br>TADRDQALQHAADSARRHGAITAYLYSTDEDYIESATDAFAW<br>AGAALTCNVTGNMPLNFSAAYSDLHVSGLNPAGNATLAEPSF<br>ITGRFRYAQVRRPLSGDGDE* |
| R7Z1F3 | ALDH-035 | MPKPFQRIREAAIDGRTQNVYYRQSQLERLRNVLVQNEEALQ<br>QAIVNDSGNSVTEAKIEYGLALLALKDQYASLNPAAALEEEY<br>AITHGKDAPQRTEGAGIVYIVPTAHTLLYSIIAPLCAAIAAG<br>NCVIVELEQTTREVPSLLRRLLAAALDVDTFDITSSRTADAE<br>FRAQCLEVLQEGSQDVPKSTQLVSPSQARVVAVVDRTANLEE<br>AAKAIVTARFSFGGKSPYAPDVVLVNEFSKKAFMTAVVKESI<br>DLLTTENGSVGKGDSRRKQEGRKLLDEVRDSDLARIVTTGAN<br>GAILDVEKRGSDILHRKINECCLAVHSVKSLDDAIDFANRND<br>DLLASYTFATPATAKYLSQFLHSYVSFVNHVPMEMLVGPAAP<br>LNHPVSSSIRYPASLFTVPRPQYVTLSPRSSLIKQTLPTASS<br>VVLTQLLEEARADLSIDKKRPERVAVGFFEQGIITGLSFVLV<br>STLAGLGTLGYFGTRMVRTRFL* |
| A0A3D2UMU3 | ALDH-036 | MSFPEIKMPELEEVKKHFKLFDYIDGERIAPTVDMKAYVHNP<br>NTGEKIAKQMATSNENIERALQVAQRLHESGEWANTSPEKRA<br>EILDNAANHLMGLVPMIAAVESYNTGIVFSTTSFVNAIVWLA<br>FKAAAGVLKSGYAVTKVPGPNGDVEVLRKPLGVAVCIVPWNS<br>PAALAAHKIANALAAGCPVILKPTEWAPYSCLLLADGLEAAG<br>LPKGLFQIVNGGAEVGNKLVADKRVRAVSFTGGLIGGRAVAE<br>ACAKDFKSLQLELGGNNAMVVLQDADVDKVADGIVTGMTTLN<br>GQWCRALGRLVVHESIQEQVVKAAIEKFKQIKVGDAMSTESQ<br>LGPLANEVHKGRIVDSINALKQKGGQTHQSAPLPDSAGFYLS<br>PTLITGVAPEETLEEIFGPVATVHTFKTDDEAIKLANQTPYG<br>LGGYVYSKDESHAMEIARKMNTGGVKVNGVSLLELNVDAPRP<br>AWGLSGFGEEGTLETFDVFCGKSVVGIAGK* |
| P05091 | ALDH-037 | MLRAAARFGPRLGRRLLSAAATQAVPAPNQQPEVFCNQIFIN<br>NEWHDAVSRKTFPTVNPSTGEVICQVAEGDKEDVDKAVKAAR<br>AAFQLGSPWRRMDASHRGRLLNRLADLIERDRTYLAALETLD<br>NGKPYVISYLVDLDMVLKCLRYYAGWADKYHGKTIPIDGDFF<br>SYTRHEPVGVCGQIIPWNFPLLMQAWKLGPALATGNVVVMKV |

| | | |
|---|---|---|
| | | AEQTPLTALYVANLIKEAGFPPGVVNIVPGFGPTAGAAIASH EDVDKVAFTGSTEIGRVIQVAAGSSNLKRVTLELGGKSPNII MSDADMDWAVEQAHFALFFNQGQCCCAGSRTFVQEDIYDEFV ERSVARAKSRVVGNPFDSKTEQGPQVDETQFKKILGYINTGK QEGAKLLCGGGIAADRGYFIQPTVFGDVQDGMTIAKEEIFGP VMQILKFKTIEEVVGRANNSTYGLAAAVFTKDLDKANYLSQA LQAGTVWVNCYDVFGAQSPFGGYKMSGSGRELGEYGLQAYTE VKTVTVKVPQKNS* |
| P51647 | ALDH-038 | MSSPAQPAVPAPLANLKIQHTKIFINNEWHDSVSGKKFPVLN PATEEVICHVEEGDKADVDKAVKAARQAFQIGSPWRTMDASE RGRLLNKLADLMERDRLLLATIEAINGGKVFANAYLSDLGGS IKALKYCAGWADKIHGQTIPSDGDIFTFTRREPIGVCGQIIP WNFPLLMFIWKIGPALSCGNTVVVKPAEQTPLTALHMASLIK EAGFPPGVVNIVPGYGPTAGAAISSHMDVDKVAFTGSTQVGK LIKEAAGKSNLKRVTLELGGKSPCIVFADADLDIAVEFAHHG VFYHQGQCCVAASRIFVEESVYDEFVRKSVERAKKYVLGNPL TQGINQGPQIDKEQHDKILDLIESGKKEGAKLECGGGRWGNK GFFVQPTVFSNVTDEMRIAKEEIFGPVQQIMKFKSIDDVIKR ANNTTYGLAAGVFTKDLDRAITVSSALQAGVVWVNCYMILSA QCPFGGFKMSGNGRELGEHGLYEYTELKTVAMKISQKNS* |
| P77674 | ALDH-039 | MQHKLLINGELVSGEGEKQPVYNPATGDVLLEIAEASAEQVD AAVRAADAAFAEWGQTTPKVRAECLLKLADVIEENGQVFAEL ESRNCGKPLHSAFNDEIPAIVDVFRFFAGAARCLNGLAAGEY LEGHTSMIRRDPLGVVASIAPWNYPLMMAAWKLAPALAAGNC VVLKPSEITPLTALKLAELAKDIFPAGVINILFGRGKTVGDP LTGHPKVRMVSLTGSIATGEHIISHTASSIKRTHMELGGKAP VIVFDDADIEAVVEGVRTFGYYNAGQDCTAACRIYAQKGIYD TLVEKLGAAVATLKSGAPDDESTELGPLSSLAHLERVGKAVE EAKATGHIKVITGGEKRKGNGYYYAPTLLAGALQDDAIVQKE VFGPVVSVTPFDNEEQVVNWANDSQYGLASSVWTKDVGRAHR VSARLQYGCTWVNTHFMLVSEMPHGGQKLSGYGKDMSLYGLE DYTVVRHVMVKH* |
| P23883 | ALDH-040 | MNFHHLAYWQDKALSLAIENRLFINGEYTAAAENETFETVDP VTQAPLAKIARGKSVDIDRAMSAARGVFERGDWSLSSPAKRK AVLNKLADLMEAHAEELALLETLDTGKPIRHSLRDDIPGAAR AIRWYAEAIDKVYGEVATTSSHELAMIVREPVGVIAAIVPWN FPLLLTCWKLGPALAAGNSVILKPSEKSPLSAIRLAGLAKEA GLPDGVLNVVTGFGHEAGQALSRHNDIDAIAFTGSTRTGKQL LKDAGDSNMKRVWLEAGGKSANIVFADCPDLQQAASATAAGI FYNQGQVCIAGTRLLLEESIADEFLALLKQQAQNWQPGHPLD PATTMGTLIDCAHADSVHSFIREGESKGQLLLDGRNAGLAAA IGPTIFVDVDPNASLSREEIFGPVLVVTRFTSEEQALQLAND SQYGLGAAVWTRDLSRAHRMSRRLKAGSVFVNNYNDGDMTVP FGGYKQSGNGRDKSLHALEKFTELKTIWISLEA* |
| UPI001AE6B188 | ALDH-041 | MRYAYPTETDSLIQLRQRYGNFINNAFVPPVQGNYFTNTSPV NGSVIGEFPRSDKEDVERALDAAHAAAESWGKTSVQQRSLLL LKIADRLEENLERLAVNETWDNGKPVRETLAADLPLAVDHFR YFAGCLRAQEGSAAEIDEFTAAYHFHEPLGVVAQIIPWNFPL LMAAWKLAPALGAGNCVVLKPAEQTPLSITLFVDLIKDILPP GVLNVIHGFGREAGEALAGNVRIAKVAFTGSTATGGHILELA AKNLIPSTVELGGKSPNIFFEDIMQAEPSFIEKAAEGLVLGF |

| | | |
|---|---|---|
| | | LNQGEICTCPSRALVQESIYEPFLAAVMKRIATIKRGDPMDS<br>ETMVGAQASQQQFDKILSYLDIARQEGAKVLTGGGVEKLEGG<br>LATGYYIQPTLLKGNNTMRVFQEEIFGPVIGITTFKDEAEAI<br>AIANDSIYGLGAGVWTRDINRAYRVGRAIKAGRVWTNCYHLY<br>PAHAAFGGYKKSGIGRETHKMMLDHYQQTKNLLVSYSTEPLG<br>FF* |
| A4IT08 | ALDH-042 (GtALDH) | MIYAQPGQPGALVTFKKRYENFIGGKWVPPVDGEYFENITPI<br>TGKPYCEVPRSKAADIELALDAAHAAKDEWGRTSPAKRARLL<br>NKIADRMEENLELLAVAETWENGKPIRETLAADIPLAIDHFR<br>YFASCIRAQEGTISEIDHDTVAYHFKEPLGVVGQIIPWNFPI<br>LMAAWKLAPALAAGNCVVLKPAEQTPTSILVLIELIEDLLPP<br>GVVNIVNGFGLEAGKPLASNPRVAKVAFTGETTTGRLIMQYA<br>SQNIVPVTLELGGKSPNIFFADVMDKDDEFFDKALEGFTMFA<br>LNQGEVCTCPSRALIHESIYDAFMERALERVKQIKQGNPLDT<br>ETMIGAQASSEQLEKILSYIDIGKQEGAELLIGGERNMLEGE<br>LAGGYYVKPTIFKGHNKMRIFQEEIFGPVLAVTTFKDEDEAL<br>AIANETLYGLGAGVWTRNINTAYRFGRGIQAGRVWTNCYHVY<br>PAHAAFGGYKMSGIGRETHKMMLDHYQQTKNMLVSYSPKKLG<br>LF* |
| P25526 | ALDH-043 | MKLNDSNLFRQQALINGEWLDANNGEAIDVTNPANGDKLGSV<br>PKMGADETRAAIDAANRALPAWRALTAKERATILRNWFNLMM<br>EHQDDLARLMTLEQGKPLAEAKGEISYAASFIEWFAEEGKRI<br>YGDTIPGHQADKRLIVIKQPIGVTAAITPWNFPAAMITRKAG<br>PALAAGCTMVLKPASQTPFSALALAELAIRAGVPAGVFNVVT<br>GSAGAVGNELTSNPLVRKLSFTGSTEIGRQLMEQCAKDIKKV<br>SLELGGNAPFIVFDDADLDKAVEGALASKFRNAGQTCVCANR<br>LYVQDGVYDRFAEKLQQAVSKLHIGDGLDNGVTIGPLIDEKA<br>VAKVEEHIADALEKGARVVCGGKAHERGGNFFQPTILVDVPA<br>NAKVSKEETFGPLAPLFRFKDEADVIAQANDTEFGLAAYFYA<br>RDLSRVFRVGEALEYGIVGINTGIISNEVAPFGGIKASGLGR<br>EGSKYGIEDYLEIKYMCIGL* |
| See footnote‡ | ALDH-044 | MSVPVQHPMYIDGQFVTWRGDAWIDVVNPATEAVISRIPDGQ<br>AEDARKAIDAAERAQPEWEALPAIERASWLRKISAGIRERAS<br>EISALIVEEGGKIQQLAEVEVAFTADYIDYMAEWARRYEGEI<br>IQSDRPGENILLFKRALGVTTGILPWNFPFFLIARKMAPALL<br>TGNTIVIKPSEFTPNNAIAFAKIVDEIGLPRGVFNLVLGRGE<br>TVGQELAGNPKVAMVSMTGSVSAGEKIMATAAKNITKVCLEL<br>GGKAPAIVMDDADLELAVKAIVDSRVINSGQVCNCAERIYVQ<br>KGIYDQFVNRLGEAMQAVQFGNPAERNDIAMGPLINAAALER<br>VEQKVARAVEEGARVALGGKAVEGKGYYYPPTLLLDVLQEMS<br>IMHEETFGPVLPVVAFDTLEEAISMANDSDYGLTSSIYTQNL<br>NVAMKAIKGLKFGETYINRENFEAMQGFHAGWRKSGIGGADG<br>KHGLHEYLQTQVVYLQS* |
| P51977 | ALDH-045 | MSSSAMPDVPAPLTNLQFKYTKIFINNEWHSSVSGKKFPVFN<br>PATEEKLCEVEEGDKEDVDKAVKAARQAFQIGSPWRTMDASE<br>RGRLLNKLADLIERDRLLLATMEAMNGGKLFSNAYLMDLGGC<br>IKTLRYCAGWADKIQGRTIPMDGNFFTYTRSEPVGVCGQIIP<br>WNFPLLMFLWKIGPALSCGNTVVVKPAEQTPLTALHMGSLIK<br>EAGFPPGVVNIVPGYGPTAGAAISSHMDVDKVAFTGSTEVGK<br>LIKEAAGKSNLKRVSLELGGKSPCIVFADADLDNAVEFAHQG<br>VFYHQGQCCIAASRLFVEESIYDEFVRRSVERAKKYVLGNPL |

| | | |
|---|---|---|
| | | TPGVSQGPQIDKEQYEKILDLIESGKKEGAKLECGGGPWGNK<br>GYFIQPTVFSDVTDDMRIAKEEIFGPVQQIMKFKSLDDVIKR<br>ANNTFYGLSAGIFTNDIDKAITVSSALQSGTVWVNCYSVVSA<br>QCPFGGFKMSGNGRELGEYGFHEYTEVKTVTIKISQKNS* |
| UPI00019FFBE0 | ALDH-046 | MLKTNIELKPKVEAFLNEEIKMFINGEFVSAIGGKTFETYNP<br>ATEDVLAVVCEAQEEDIDAAVKAARSAFESGPWAEMTTAERA<br>HLIYKLADLIEEHREELAQLEALDNGKPYQVALDDDISATVE<br>NYRYYAGWTTKIIGQTIPISKDYLNYTRHEPVGVVGQIIPWN<br>FPLVMSSWKMGAALATGCTIVLKPAEQTPLSLLYAAKLFKEA<br>GFPNGVVNFVPGFGPEAGAAIVNHHDIDKVAFTGSTVTGKYI<br>MRQSAEMIKHVTLELGGKSPNIILEDADLEEAINGAFQGIMY<br>NHGQNCSAGSRVFVHRKHYETVVDALVKMANNVKLGAGMEKE<br>TEMGPLVSKKQQERVLNYIEQGKKEGATVAAGGERALEKGYF<br>VKPTVFTDVTDDMTIVKEEIFGPVVVVLPFDSTEEVIERANN<br>SSYGLAAGVWTQNIKTGHQVANKLKAGTVWINDYNLENAAAP<br>FGGYKQSGIGRELGSYALDNYTEVKSVWVNIK* |
| Q9LRI6 | ALDH-047 | MAARRAASSLLSRGLIARPSAASSTGDSAILGAGSARGFLPG<br>SLHRFSAAPAAAATAAATEEPIQPPVDVKYTKLLINGNFVDA<br>ASGKTFATVDPRTGDVIARVAEGDAEDVNRAVAAARRAFDEG<br>PWPRMTAYERCRVLLRFADLIEQHADEIAALETWDGGKTLEQ<br>TTGTEVPMVARYMRYYGGWADKIHGLVVPADGPHHVQVLHEP<br>IGVAGQIIPWNFPLLMFAWKVGPALACGNAVVLKTAEQTPLS<br>ALFVASLLHEAGLPDGVLNVVSGFGPTAGAALSSHMGVDKLA<br>FTGSTGTGKIVLELAARSNLKPVTLELGGKSPFIVMDDADVD<br>QAVELAHRALFFNQGQCCCAGSRTFVHERVYDEFVEKARARA<br>LQRVVGDPFRTGVEQGPQIDGEQFKKILQYVKSGVDSGATLV<br>AGGDRAGSRGFYIQPTVFADVEDEMKIAQEEIFGPVQSILKF<br>STVEEVVRRANATPYGLAAGVFTQRLDAANTLARALRVGTVW<br>VNTYDVFDAAVPFGGYKMSGVGREKGVYSLRNYLQTKAVVTP<br>IKDAAWL* |
| C9DIJ2 | ALDH-048 | MLRTATRTTFKSAQPSFMAAAAALRYYSHYPLSKSITLPNGK<br>VYEQPTGLFINGEFVASQQHKTFEVINPSNEDEICHVYEART<br>EDVDAAVDAAYNAFHSEWSKMDPSIRGEHLMKLAELMEKNKD<br>TLAAIESMDNGKALFMAEIDVKLVINYLKYCAGWADKLFGKV<br>VDTGSNYFNYIKREPIGVCGQIIPWNFPLLMWSWKVGPALAA<br>GNTIVLKTAESTPLSALYAAKLAKEAGIPDGVINIVSGFGKI<br>TGEAISTHPKIKKLAFTGSTATGKHIMKAAAESNLKKVTLEL<br>GGKSPNIVFNDADIKKAVNNLILGIFFNSGEVCCAGSRVFIQ<br>EGVYDQVLEEFKIAAEALKVGNPFEEGVFQGAQTSQQQLTKI<br>LGYVESGKDEGATLVTGGERLGDKGYFVKPTIFADVKPNMKI<br>YSEEIFGPFAVVTKFKTADEAIAMANDSEYGLAAGIHTTSLD<br>TATYVANNLEAGTVWINTYNDFHHNMPFGGFKQSGIGREMGE<br>AAFENYTQWKTVRIAINDGPQ* |
| Q65NX0 | ALDH-049 | MSVAAESKTYFNFINGRWVKAESGGMEQSLNPADTRDIVGLV<br>QKSSIEDVDRAVEAAKQAKKAWRKLSGAERGQFLYKAADIME<br>QRLDEIAECATREMGKTLPEAKGETARGIAILRYYAGEGLRK<br>TGDVIPSTDSSAFMYTDRVPLGVVGVISPWNFPVAIPIWKMA<br>PALIYGNTVVIKPATETAVTCLKVISCFEEAGIPSGVVNAVT<br>GPGSSAGQRLAEHPDVNGITFTGSNQTGKIIGRTAFERGAKY<br>QLEMGGKNPVIVADDADLDIAVEAVISGAFRSTGQKCTATSR<br>VIVLNGVYDRFKEKLLQQTKEITIGDSLKEDVWMGPIANKQQ |

| | | |
|---|---|---|
| | | LDNCLSYIAKGKQEGADLIFGGERLADGKYENGYYIRPAIFD<br>NVTSGMTIAQEEIFGPVIALIKADTLEEALETANDVKFGLSA<br>SIFTQNIRRMLSFTDEIEAGLIRVNAESAGVELQAPFGGVKQ<br>SSSHSREQGEAAKEFFTAVKTVFVKP* |
| UPI00005BF137 | ALDH-050 | MSAISEVVQRARAAFNSGRTRPLQFRVQQLEGLRRLIREREK<br>DLVGALAADLHKNEWTAYYEEIVYVLEEIDYMIRKLPEWAAD<br>EPVEKTPHTQQDEAYIHSEPLGVVLIIGSWNYPFNLTIQPMV<br>GAIAAGNAVVLKPSELSENTASLLATILPQYLDQDLYPVING<br>GVAETTEVLKERFDHILFTGSTGVGRVVMMAAAKHLTPVTLE<br>LGGKNPCYVDKDCDLDIACRRIAWGKFMNSGQTCVAPDYILC<br>DPSIQSQVVEKLKKSLKEFYGEDAKKSRDYGRIINSRHFQRV<br>MGLLEGQKVTYGGTGDATTRYIAPTILTDVDPESPVMQEEVF<br>GPVLPIMCVRSLEEAIQFITQREKPLALYVFSPNDKVIKKMI<br>AETSSGGVTANDVVVHISVHSLPYGGVGDSGMGSYHGRKSFE<br>TFSHRRSCLVRPLLNEETLKARYPPSPAKMPRH* |
| P20000 | ALDH-051 | MLRAVALAAARLGPRQGRRLLSAATQAVPTPNQQPEVLYNQI<br>FINNEWHDAVSKKTFPTVNPSTGDVICHVAEGDKADVDRAVK<br>AARAAFQLGSPWRRMDASERGRLLNRLADLIERDRTYLAALE<br>TLDNGKPYIISYLVDLDMVLKCLRYYAGWADKYHGKTIPIDG<br>DYFSYTRHEPVGVCGQIIPWNFPLLMQAWKLGPALATGNVVV<br>MKVAEQTPLTALYVANLIKEAGFPPGVVNVIPGFGPTAGAAI<br>ASHEDVDKVAFTGSTEVGHLIQVAAGKSNLKRVTLELGGKSP<br>NIIMSDADMDWAVEQAHFALFFNQGQCCCAGSRTFVQEDIYA<br>EFVERSVARAKSRVVGNPFDSRTEQGPQVDETQFKKVLGYIK<br>SGKEEGAKLLCGGGAAADRGYFIQPTVFGDVQDGMTIAKEEI<br>FGPVMQILKFKSMEEVVGRANNSKYGLAAAVFTKDLDKANYL<br>SQALQAGTVWVNCYDVFGAQSPFGGYKLSGSGRELGEYGLQA<br>YTEVKTVTVRVPQKNS* |
| A0A6H1TS81 | ALDH-053 | MLTATPVREIIERQRQFFATGRTKSVDFRIEQLKKLKQAILD<br>HEAEIIAAVQADLRKPHLEAYLTEIGSVSKIDYALKQIKSWV<br>KPQKVATGIEQFPASARVYSEPLGVVLIISPWNYPFNLAIEP<br>LIGAIAAGNCAIVKPSEVSANTSRVIAKLFGAVFDPGYISVV<br>EGDAEVSQGLLAEKFDHIFFTGGTAIGQRVMEAAAKQLTPVT<br>LELGGKSPCIVDPQINLETAATRVTWGKFLNAGQTCIAPDYL<br>LVDRRIQADFVAEIQKKLHQFFGPSPQESPDFGRIVSDKHFQ<br>RLASLLQDAQIVTGGELDPGDRYIAPTLVENVALDAPLMQEE<br>IFGPILPIIPYDRFDEAIAIVNQRPKPLALYLFSNDKEKQAR<br>IVRETSSGGVCLNDTIMHVGVAELPFGGVGPSGIGAYHGKAS<br>FDTFSHRKSVLKKSFWLDLDLRYPPYAGKLKLVKKFLGQ* |
| A0A7L1RJ35<br>+A0A7L1RY59[†] | ALDH-054 | MERMQQIVGRARAAFNSGRSRPLEFRIQQLKALERMVQEKEK<br>EILAALKADLNKCGHNAYSHEILGVLGELAQTMEKLPSWAAP<br>QPVKKNLLTMRDEAYINYEPLGVVLVIGAWNYPFVLVMQPLI<br>GAIAAGNAVVVKPSEVSENTAQLVAELLPQYLDKELYPVVTG<br>GVPETTELLTQRFDHILYTGNTAVGKIVMAAAAKHLTPVTLE<br>LGGKSPCYIDKDCDLAVACRRITWGKYMNCGQTCIAPDYILC<br>DPSIQGKVVENIKATLKEFYGEDVKSSPDYGRIVSQRHFKRV<br>MSLLEGQKIAHGGETDEASCFIAPTILTDVSPESKVMEEEIF<br>GPVLPIVTVRSVEEAIEFINRREKPLALYVFSNNKQLIKRVI<br>SETSSGGVTGNDVIMHFFLSTLPFGGVGHSGMGAYHGKHSFE<br>TFSHRRACLIKDLKMESTNKMRYPPGSQKKV* |

| | | |
|---|---|---|
| A0A6N7YK66 | ALDH-055 | MTRYADPGTAGSVVSFASRYGHFIGGEYVPPAKGEYFENITP<br>ITGRPFTEIARGTAEDIDRALDAAEAAAPAWGRTSVTERAGI<br>LHKMADRMEAGLEKLAVAESWDNGKPCRETLAADIPLAVDHL<br>RYFASTIRAQEGGISEIDGDTVAYHFHEPLGVVGQIIPWNFP<br>LLLAVWKIAPALAAGNTIVLKPAEQTSASIHILLDVVSDLLP<br>PGVLNVVTGFGIEAGKPLAASPRIRKISFTGETGTGRLIMQY<br>ASENLIPVTLELGGKSPNIFFEDVAAADDAFYDKAVEGFTMF<br>ALNQGEVCTCPSRAVIHSRVYDEFLQRAVERTEKITQGHPLD<br>TETMLGAQASAEQYEKILRYLDIGRGEGAKVRTGGEKADLGG<br>ELSGGYYIRPTVFEGDNSMRIFQEEIFGPVVSVTRFDDYAEA<br>VKIANDTRYGLGAAVWSRDGNTAYRAGRDIQSGRVWVNNYHA<br>YPAHAAFGGYKQSGIGRENHKMMLEHYQQTKNILVSYSQNAQ<br>GFY* |
| A0A2T0SSF3 | ALDH-056 | MAEYAAPGQPGSVVSYLDRYDHFIGGEYVAPAKGDYFENPTP<br>VTGRAFTEIARGTADDVERALDAAHGAARAWGRTSPAERANV<br>LNRIADRMEQNLEALAVAECWENGKAVRETLNADIPLAIDHF<br>RYFAGVIRAQEGGISQIDQDLVAYHFQEPLGVVGQIIPWNFP<br>ILMATWKLAPALAAGNCVVLKPAEQTPASIHVLLSFIADLLP<br>PGVLNVVNGFGVEAGKPLASSSRIAKIAFTGETTTGRLIMQY<br>ASENIIPVTLELGGKSPNVFFSDVASSRDSFYDKALEGFTMF<br>ALNQGEVCTCPSRALIQSSIYDSFLADATERTKLIKQGNPLD<br>TDTMMGAQASNDQLEKILSYFDIGRQEGAKVVTGGERADLGG<br>DLSGGYYVQPTIFEGDNKMRIFQEEIFGPVVSVTRFDDYDDA<br>VKTANDTLYGLGAAVWSRDGGTAYRAGRDIQAGRVWVNNYHA<br>YPAHAAFGGYKDSGIGRENHKMMLDHYQQTKNLLVSYSQNAQ<br>GFF* |
| A0A918MZJ5 | ALDH-057 | MDLSELKQFGLDVAYPYKKKYENFIGGQWVAPVSGEYFENIS<br>PITGKPFCQVARSGAQDVELALDKAHAARRAWGKTSVMDRSN<br>LLLRMADRMEKNLQLLAVAETIDNGKPIRETMAADLPLAIDH<br>LRYFAGCIRAQEGAISEIDETTVAYHFQEPLGVVGQIIPWNF<br>PILMAIWKLAPALAAGCCVVLKPAEQTPASILVLMELTADIF<br>PPGVINVINGFGKEAGNALATNKRIAKIAFTGSTATGKHILH<br>AAAENLIPSTVELGGKSPNIFFADVMDQSDAFLDKALEGLAM<br>FSLNQGEVCTCPSRILIQESIYEAFIEKAIKRVQAIKTGNPL<br>EASTMVGAQASRMQMDKIMSYIDIGREEGAVCLTGGDRHRPG<br>NLEDGYYINPTMLLGKNNMRIFQEEIFGPVASVTTFKDEEEA<br>LEIANASYYGLGAGVWSRDGTRAYRMGRGIEAGRVWTNCYHL<br>YPAHAAFGGYKQSGIGRETHKMALGNYQQTKCLLVSYSANAL<br>GFF* |
| A0A522WLN6 | ALDH-058 | MLYANPGTPGALVQFKARYDNFIGGKWVPPLKGDYFDVVSPG<br>TGKVYTKASRSGAEDIELALDAAHAAADQWGRTPPAERANLL<br>LRIADRIEANLEVLAYAETVDNGKPIRETLNADIPLTIDHFR<br>YFAGCLRAQEGGLSDIDGNTVAYHFHEPLGVVGQIIPWNFPI<br>LMAAWKLAPAIGAGNCVVLKPAESTPISIMILAELIADLLPP<br>GVLNIVNGYGREAGMPLASSRRIAKIAFTGSTATGRVIAQAA<br>ANNLIPATLELGGKSPNIFFADVAEADDGFLDKAIEGLVLFA<br>FNQGEVCTCPSRALIQDSIYDKFMDRVLPRVAAIKQGNPLDT<br>ETMIGAQASKEQLTKILSYLDLGKQEGARVLIGGERAHLGGE<br>LDGGYYVQPTLFQGHNKMRIFQEEIFGPVLAVTTFKDEAQAL<br>EIANDTLYGLGAGVWSRNGNVAYRMGRAIKAGRVWTNCYHAY |

| | | |
|---|---|---|
| | | PAHATFGGYKESGIGRETHKVMLDHYQQTKNLLVSYSESKLG<br>FF* |
| A0A858QBL4 | ALDH-059 | MKTESPFKPCYDNFIGGRWVPPVRGEYFENLTPVTGKPLGKV<br>ARSTADDVELALDAAHAAAAAWGRTSATVRANLLNRIADRME<br>AHLETLALAETWDNGKPVRETRAADIPLAIDHFRYFAGCIRA<br>QEGSIGELDADTVAYHFHEPLGVVGQIIPWNFPILMAAWKLA<br>PALAAGNCVVLKPAEQTPMGIMVLLELIGDLLPPGVLNVVNG<br>FGLEAGKPLASSRRIAKIAFTGETTTGRLIMQYASQNIIPVT<br>LELGGKSPNIFFADVCAREDDFFDKAVEGLAMFALNQGEVCT<br>CPSRALIHESIYDRFMDHALKRIAAMKQGNPLDAETMVGAQA<br>SEEQVEKILSYFEIGKQEGAECLIGGERNLLDGDLRGGYYVK<br>PTVFRGHNRMRIFQEEIFGPVLSVTTFKDDEEALAIANDTLY<br>GLGAGIWTRDINRAYRMGRGIHAGRVWVNCYHLYPAHAAFGG<br>YKQSGIGRENHKMMLAHYQQMKNLLVSYSPKAVGLF* |
| A0A1G9VJH9 | ALDH-060 | MDMFELKRFNLDIAYPYRQQYDNFIGGKWVAPQAGRYFDNVS<br>PITGKPFCKVAQSDAADVNRALDAAHAAQAQWGSTAPAERSR<br>ILLKIADRMEENLELLAIAETIDNGKPLRETMAADLPLGVDH<br>FRYFAGCLRAEEGTISEIDANTVAYHFKEPIGVVGQIIPWNF<br>PLLMACWKLAPALAAGCTVVMKPAEQTPASIMVLMELIGDLL<br>PPGVVNIVNGYGKEAGEALATSKRIAKIAFTGSTPVGQRILH<br>AAADNLIPSTVELGGKSPNIFFADVMDADDEFLDKALEGFSM<br>FALNQGEVCTCPSRILVQESIYDRFIERALARVEAMKQGHPL<br>DKSTMVGAQVSQQQLDRILGYIDIGRGEGAQCLTGGERLRAG<br>GDIDDGFFIKPTLLYGQNHMRVFQEEIFGPVASVTTFKDAED<br>ALRIANDSEYGLGAGVWSRNGSTAYKMGRAIQAGRVWTNCYH<br>LYPAHATFGGYKKSGIGRETHKMAMANYQQTKNLLVSYSSKA<br>LGFF* |
| A0A1B1YTA3 | ALDH-061 | MRYAAPGTPGAKVSFKSRYGNFIGGQWLPPAGGEYFEDISPI<br>TGRAFCEVPRSQKDDIERALDAAHAAKAAWGRTSPAERAGLL<br>LRIADRMEQNLELLAVAETFSNGKPIRESLAADLPLAVDHWR<br>YFAGALRAQEGSLSQIDADTVAYHFHEPLGVVAQIIPWNFPL<br>LMAVWKLAPALAAGNCVVLKPAEQTPTSILVLMELLQDLIPP<br>GVINVVNGFGVEAGKPLASSPRIAKVAFTGETTTGRLIMQYA<br>SANLIPVTLELGGKSPNIFFDDVGARDDAFFDKALEGFAMFA<br>LNQGEVCTCPSRALIQESLYERFMERALKRVAAIRGGDPLDT<br>DTMIGAQSSQEQLEKILSYIDIGKQEGAQLLTGGTRVAHDGD<br>MAGGYYVAPTVFKGHNRMRIFQEEIFGPVVSVATFKDEAEAL<br>ALANDTLYGLGAAVWSREQNTCHRMARAIQSGRVWVNAYHLY<br>PAHAAFGGYKQSGIGRECHKMMLGHYQQTKNVLVSYSDQALG<br>FF* |
| R4G109 | ALDH-062 | MVIYFHLKTDTKGGVYMIYAQPGQPGSLITFKKRYENFIGGE<br>WVPPVDGEYFENISPVTGQVYCEVPRSKSADIELALDAAHAA<br>KEAWGRTSVAERARLLNKIADRMEENLDMLAVAETWENGKPI<br>RETRVADIPLAIDHFRYFAGCIRAEEGTLAELDNDTVAYHFK<br>EPLGVVGQIIPWNFPILMAAWKLAPALAAGNCVVLKPAEQTP<br>TSILVLIELIQDLLPKGVVNVVNGFGLEAGKPLASNPRIAKV<br>AFTGETTTGRLIMQYASQNLIPVTLELGGKSPNIFFADVAAK<br>DDEFFDKALEGFTMFALNQGEVCTCPSRALIEESIYDIFMER<br>ALERVKQIKQGNPLDTTTMIGAQASSEQLEKILSYIDIGKQE<br>GAELLIGGERNMLEGDLQGGYYVKPTVFKGHNRMRIFQEEIF<br>GPVVSVTTFKDKDEALAIANDTLYGLGAGVWTRDVNTAYRFG |

| | | |
|---|---|---|
| | | RGIQAGRVWTNCYHAYPAHAAFGGYKMSGIGRETHKMMLEHY<br>QQTKNLLVSYSPKKLGFF* |
| A0A261U014 | ALDH-063 | MDIATRLAPDTYGTPLDIKSQYGNFIGGKWVEPATGEFFDNL<br>TPVTGAVLSRHARSGERDIEMALDAAHKAAPAWGATPPAERA<br>RVLLRIADLIEANLEQLATAETWDNGKPIREARAADIPLAVD<br>HFRYFASAIRGQEGSLSEIDADTVAYHFREPLGVVGQIIPWN<br>FPILMAAWKLAPALAAGNCVVLKPAEQTPLGILMLAELIAEV<br>VPPGVLNIVTGFGLEAGKPLASNKRIAKIAFTGETTTGRLIM<br>QYASQNIIPVTLELGGKSPNIFFEDVGAHDDDFFDKAIEGFV<br>MFALNQGEVCTCPSRALIHESLYDRFMERALARVAQIKQGNP<br>LDADTMIGAQASSEQLEKILSYLDIGRQEGAEVLAGGARATF<br>DGALGQGYYVQPTVFKGNNRMRVFQEEIFGPVVAVTTFKDAD<br>EALSLANDTLYGLGAGVWSRDVNTCYRMGRAIKAGRVWTNCY<br>HAYPAHAAFGGYKQSGIGRENHKMMLDHYQQTKNMLVSYSPK<br>KLGFF* |
| I3XUN2 | ALDH-064 | MVYAKPIYKSQYENFIGGEWLAPLKGEYFDNLSPVDGELLTK<br>IPRSSTQDVDLAVAAGKKAFETFKHFSVLQRSELLNKIADKI<br>EANLEFLAISETLDNGKAIRETLAADLPLVVDHFRYFASVIR<br>SEAGSVADLDENTISQEVYEPLGVVAQIIPWNFPLLMAAWKI<br>APALAAGNCVVIKPASATPLSILLLMETIQEVLPKGVLNVIN<br>GAGGKIGKHLATHPDIKKVGFTGETTTGQLIMQYATENIIPS<br>TLELGGKSPNVFFPSVMAHDDDFLDKAIEGLVLFAFNSGEVC<br>TCPSRALIHESIYEPFMQRVLERVKAISQENPLDSTTKMGAQ<br>ASLNQKEKILEYIRIGKEEGAQCLIGGEAYENKTFPKGNYIK<br>PTIFKGHNKMRIFQEEIFGPVLCVTTFKDEAETLSIANDTIY<br>GLGSGVWSRDVHEVHRMSRGIEAGRVWVNCYHLYPSHASFGG<br>YKKSGIGRETHMMMLNAYRHTKNILTSYALKPLGFF* |
| N8Z6I5 | ALDH-065 | MRYIDPNQPGSKVHFKAQYENFIGGQWIVPVKGEYFDNLSPV<br>DGKVFTRVPRSSVEDIELALDAAHKAKAQWSKSSPTTRSNIL<br>LKIADRLEANLELLAVAETWDNGKPIRETLAADIPLAIDHFR<br>YFAGCIRAQEGGISEIDEDTIAYHFHEPLGVVGQIIPWNFPI<br>LMATWKLAPALAAGNCIVLKPAEQTPVSILVLVELIQDLLPE<br>GVLNIVNGYGVEVGRPLAVNPRIAKIAFTGSTSVGQQIMQYA<br>TENIIPVTLELGGKSPNIFFEDVMAQQDDYLEKALEGFTMFA<br>LNQGEICTCPSRALVQESIADQFLKLAVERVKRIKTGHPLDT<br>ETMIGAQASQEQQDKILGCIATGRGEGAEVLVGGGARHEVGQ<br>GYYIEPTIFKGTNNMRTFQEEIFGPVLAVTTFKDFDDAIKIA<br>NDTIYGLGAGVWSRSTHTAYRAGRAIEAGRVWTNCYHIYPAH<br>AAFGGYKKSGIGRENHKMMLDHYQQTKNLLVSYSTKPAGFF* |
| A0A850GCI6 | ALDH-066 | MTYAPPNSEGALVNFKSRYANFIGGEWVAPIKGEYFENPSPV<br>TGQSFCEIPRSTAADIDKALDAAHGAREAWGKTAVAERASLL<br>NKIADRIEQNLEKLAVAETWDNGKAVRETMAADLPLTVDHFR<br>YFASCIRAQEGTQGELDHDTVSYQFKEPLGVVGQIIPWNFPL<br>LMAAWKIAPALAAGNCVVLKPAEQTPASILVMLELIGDILPP<br>GVLNVVNGFGLEAGKPLASSSRIAKIAFTGETTTGRLIMQYA<br>SENLIPVTLELGGKSPNVFFSDVWAKDDDFRQKALEGFVMFA<br>LNQGEVCTCPSRALIEDDIFGEFMDAAVARTKKVKLGNPLDP<br>NTMMGAQASADQLEKILSYIDIGKQEGAELLTGGKRAELGGD<br>LAGGYYVEPTIFSGRNDMRVFQEEIFGPVVSVARFSGFEQAM<br>QIANDTLYGLGAGVWTRDGNTAYRAGRTIQAGRVWTNCYHLY |

| | | |
|---|---|---|
| | | PAHAAFGGYKQSGIGRETHEMMLDHYQQTKNLLVSYSEKPMG<br>FF* |
| UPI00234FF82E | ALDH-067 | MSTAQTTSKPKIAAPKFKSQYDNFIGGKWTPPVKGEYFENVS<br>PVDGNSFTKVARSTAEDIDKAIDAAWEAAPQWNSSSATERSN<br>MLLKIASIMEDNLEALARAETWDNGKALRETMAADLPLAVDH<br>FRYFAGVIRAEEGSVSELDANTVSLNVPEPLGVVGQIIPWNF<br>PILMATWKMAPALAAGNCVVLKPAEQTPVGIMILMELIQDVL<br>PAGVLNIVNGFGPEAGKPLASSPRINKVAFTGETTTGQLIMQ<br>YASKNITPVTLELGGKSPNVFFESVMDADDDFFDKAIEGAVM<br>FALNQGEVCTCPSRMLVQESIYEKFMERVVERTNAIKLGHPL<br>DSETMMGAQASNDQYEKILNYIEIGKDEGCEVLAGGEAAYNE<br>GLEGGYYIKPTILKGNNKMRVFQEEIFGPVVCVTTFKDEAEA<br>IEIANDTLYGLGAGVWTRDTHQAYQISRAIKAGRVWVNCYHN<br>YPAHAPFGGYKKSGIGRENHKMMLDHYRQTKNMLISYDKKAL<br>GFF* |
| A0A2U3D9B3 | ALDH-068 | MIYAQPNQPGSKVAFKSRFLNYIGGEWVAPTKGMYFTNFTPV<br>TGQPFCEAARSTADDIERALDAAHKAKISFGQTSAADRAQVL<br>NKIADRMEANLEMLAVAETWDNGKPIRETLAADIPLAIDHFR<br>YFASAIRAQEGRLSELDHDTVAYHFHEPLGVVGQIIPWNFPL<br>LMAAWKLAPALAAGNCVVLKPAEQTPASIMVLMELIGDLLPP<br>GVVNVVNGFGVEAGKPLASNPRIQKVAFTGETTTGRLIMQYA<br>SNNLIPVTLELGGKSPNIFFKDVLDEEDDFKEKAIEGFNMFA<br>LNQGEVCTCPSRALIEKSIYHDFIELAIERTKKIKQGNPLDT<br>ETMMGAQASLDQVEKIMSYVELGKQEGASVLVGGEKVCPSDD<br>LAEGYYIAPTVFVGHNRMRIFQEEIFGPVLSVTTFENFDDAV<br>HIANDTLYGLGAGVWTRDFHLAYRMGRAIQAGRVWVNNYHAY<br>PAHAAFGGYKQSGIGRENHLMMLEHYQQTKNLLVNYSKKPLG<br>FF* |
| A0A1W9L4H0 | ALDH-069 | MIYAQPGSPNAKVTFKTRYDNYIGGAWVKPLDNEYFENLSPV<br>TGKPFCEVARSKAQDIELALDAAHAAKAKWGKTSVTDRANLL<br>NKMADRMEANLEKIAVAETWENGKPVRETLAADIPLAIDHLR<br>YFAGCIRAQEGALSELDEHTVAYHFHEPLGVVGQIIPWNFPI<br>LMAIWKLAPAIAAGNCVILKPAEQTPVSILLLMELLTDLLPP<br>GVLNIVNGFGLEAGKPLAANKRIAKIAFTGETTTGRLIMQYA<br>SQNIIPVTLELGGKSPNIFFADVCDQDDAFFDKALEGFTLFA<br>LNQGEVCTCPSRALIQESIYDRFMERAQVRVQQIKQGHPLDT<br>ETMIGAQVSTEQLEKILSYIDIGKQEGAQILQGGQRQRFPGE<br>LADGYYIEPTLFKGHNKMRIFQEEIFGPVLAITTFKDEAEAL<br>DMANDTLYGLGAGVWTRDGSRAYRLGRGIQAGRVWVNCYHAY<br>PAHAAFGGYKQSGIGRETHKIILEHYQQTKNLLVSYNPNKLG<br>FF* |
| A0A3S0VUQ7 | ALDH-070 | MLYIEPGQPGAKVAFKTRYANFIGGEWRPPVEGRYFENLTPI<br>TGKAFCEVPRSSAADIELALDAAHAAKAAWGRTSPTERANLL<br>LKIADRMEANLAMLAVAETWDNGKPIRESLAADLPLAIDHFR<br>YFAGCIRAQEGTLSQIDDDTVAYHFHEPLGVVGQIIPWNFPI<br>LMAVWKLAPALAAGNCVVLKPAEQTPVSILVLLELIQGVLPP<br>GVLNVVNGFGLEAGKPLATNKRIAKIAFTGETTTGRLIMQYA<br>TENIIPVTLELGGKSPNLFFADVMAQDDDFFDKALEGLAMFA<br>LNQGEVCTCPSRALIQESIYDAFIARAIERVGRIKQGHPLDT<br>DTMLGAQVSSEQMEKIISYLDIGRQEGAKCLIGGGRNALGGG<br>LAGGYYVQPTLLQGHNRMRIFQEEIFGPVLAVTTFKDEAEAL |

| | | |
|---|---|---|
| | | SIANDTLYGLGAGVWTRDANRAYRFGRGIQAGRVWVNCYHAY<br>PAHAAFGGYKQSGIGRETHKMMLDHYQQTKNLLVSYSPKALG<br>FF* |
| A0A177P8X5 | ALDH-071 | MTRPEFLRSTTPKFEARYENFIGGRWVAPRAGRYFENVTPVN<br>GQILCEVPRSDATDIELALDAAHAAKDAWGRASVGERARVLH<br>AVADRMEQNLAALAEAESWDNGKPIRETTAADIPLAIDHFRY<br>FASAIRAQEGGVSEIDHDTVAYHFHEPLGVVGQIIPWNFPLL<br>MAAWKLAPAIAAGNCVVLKPAEQTPASILVWAKIVGDLLPAG<br>VINIVNGFGLEAGKPLASSPRIAKIAFTGETSTGRLIGEYAA<br>RNLIPTTLELGGKSPNIFFADVAAEDDAFLDKALEGFAMFAL<br>NQGEVCTCPSRALVHRSIYDKFIEKAVKRVGAIVQGDPLDAR<br>TMIGAQASKEQMEKILGYIDIGQKEGADILIGGGRAELGGDL<br>SGGYYVKPTVLAGNNKMRVFQEEIFGPVVSVTVFDTDEEAVA<br>IANDTIFGLGAGVWTRDINRAYRVGRAVQAGRVWTNCYHAYP<br>AHAAFGGYKQSGIGRENHRMMLDHYQQTKNLLVSYSPDKLGF<br>F* |
| A0A7Z0SSR0 | ALDH-072 | MDLTQLKYFGIDVEYPYKKHYENYIGGQWVAPVAGNYFENVS<br>PITGQTFCEVPRSDAADVDLALDAAHAARRKWGETSPTERAG<br>ILLRIADVMEKHLKVLAVAESIDNGKPLRETMAADVPLAIDH<br>FRYFAGCIRAQEGGISEIDKDTVAYHFYEPMGVVGQIIPWNF<br>PLLMAAWKLGPALAAGNCVVLKPAEQTPASILVLVELIGGLL<br>PPGVLNIVNGFGKEAGNALATSKRISKIAFTGSTPTGKHILH<br>AAADNLIPSTVELGGKSPNIFFADVMDQDDAFFDKALEGLAM<br>FALNQGEVCTCPSRVLIQESIYDQFIERAVKRVESIKTGHPL<br>DSKTMVGAQVSQVQMDKILSYIDIGRAEGATCLTGGARNAPG<br>EGLEQGFYVKPTLLFGENSMRIFQEEIFGPVASVMKFKDEDH<br>AIEIANDTYFGLGAGVWTRDGSRAYRVGRQVQAGRVWTNCYN<br>LYPAHAAFGGYKQSGIGRETHKVALQAYQQTKCLLVSYSPNA<br>LGFF* |
| UPI000423BFB2 | ALDH-073 | MIYAQPGQADAKVSFKKRYDNFIGGKWVPPVKGEYFENITPV<br>TGKPFCEVARSTSDDIELALDAAHAAKDSWGRTSAIERASLL<br>NKIADRMEANLEALAVAETWDNGKPVRETLAADIPLAIDHFR<br>YFAGCIRVQEGSIGQIDNDTVAYHFHEPLGVVGQIIPWNFPI<br>LMAVWKLAPALAAGNCIVLKPAEQTPASVLVLMELIQDLLPP<br>GVVNIVNGFGLEAGKPLASSNRIAKIAFTGETTTGRLIMQYA<br>SQNIIPVTLELGGKSPNIFFENVFNKDDAFLNKAIEGFVLFA<br>LNQGEVCTCPSRALIQESIYDRFMERALKRVADIKQGNPLDT<br>ETMIGAQASSEQVEKILSYMDIGKQEGAQCLIGGERNTLEGD<br>LQDGYYIKPTVFKGHNKMRIFQEEIFGPVVSVTTFKDFDEAL<br>EIANDTLYGLGSGVWTRDINDAYRMGREIQAGRVWTNCYHSY<br>PAHAAFGGYKGSGIGRETHKMMLEHYQQTKNLLVSYSPDALG<br>FF* |
| A0A543HTC7 | ALDH-074 | MTVYAQPGTPGSVITFQEQYENFIGGQWVPPVEGQYFDVTTP<br>VTGQAFTRAPRSTAPDVELALDAAWEAFPAWSKLSATQRAEI<br>LWKIGDRIDENLEKIAVADTWDNGKPVRETLAADIPLAADHF<br>RYFAAAIRAQESNTNEVNEDLVAYHFQEPLGVVGQIIPWNFP<br>ILMAAWKIAPALAAGNTIVLKPASETPVSLLEFVKSFEDLLP<br>AGVLNIVNGAGSKVGNALANSTRIRKIAFTGSTSVGRSIAQS<br>AANLIIPATLELGGKSPNVFFADVADKDDAFYQKALEGFSLF<br>QLNQGEICTCPSRALVQESIYEKFLTDAVARVESAIQGNPLD<br>TNTQVGAQVSENQMNTILGYIKVGKEEGAEVLTGGERNILEG |

| | | |
|---|---|---|
| | | DLAGGFYVKPTVFKGQNSMRVFQEEIFGPVLSVTSFTDFDDA<br>ISIANDTIYGLGAGVWSRSGRIQYKAGRAIQAGRVWTNTYHN<br>YPAGAAFGGYKESGIGRETHKMVLDAYQQTKNLLVSYSEEPQ<br>GLY* |
| A0A4V3IHV4 | ALDH-075 | MTIYPAPGTPGASVTFKPRYENWIGGEWIKPVKGQYFDNVSP<br>VNGKSFTEAARGTVEDIELALDAAHAAAPAWGKTSAAERAAV<br>LNRIADRIDANLELLAVAETWDNGKPVRETLAADIPLAADHF<br>RYFAAAIRSQDGDHAELDGDNVAYQFHEPLGVVGQIIPWNFP<br>ILMASWKLAPAIAAGNAIVLKPAEQTPVSILVLMELLADILP<br>AGVINVVNGFGQEAGAALAASTRIRKIGFTGSTPVGRLILKA<br>ASANIIPTTVELGGKSPNIFFDDIDAKRDAFYDKAQEGFALF<br>AFNQGEVCTAPTRALVQKSMYATFVGDAVQRTAKAIQGNPLD<br>TATQVGAQVSAEQLEQILGYIEIGKQEGATLLLGGKQVDLGG<br>DLSGGFYVEPTIFEGTNDMRIFQEEIFGPVLAVTSFDDYDDA<br>IAIANDTEFGLGAGVWSRNGNITYRAGRDIQAGRVWVNNYHT<br>YPAGAAFGGYKNSGIGRENNKLALGHYQQTKTLLVSYSEDAL<br>GLF* |
| A0A2X4UZ81 | ALDH-076 | MKYVHPGLPGSLVSFKQRYGNYIGGKFVEPVSGNYFTDTTPV<br>TGKTVAEFPRSDAQDIERALDAAHAAAEDWGNTSAQNRANVM<br>LAIADRMESQLEMLALTESWDNGKPIRETLNADIPLAIDHFR<br>YFAGCLRAQEGTATEIDQNTLAYHIYEPLGVVGQIIPWNFPL<br>LMAAWKLAPALTAGNCVVLKPAEQTPLSICVLLELIGDLLPP<br>GVLNVVHGFGKEAGEALATSKRIDKIAFTGSTPVGRHILACA<br>AENIIPSTVELGGKSPNIFFEDIMQAEPEFIDKAVEGLILGF<br>FNQGEVCTCPSRALIQESIYPQFMEKVLARITTIRQGDPFDT<br>DTMIGAQASQQQFDKILSYIEIAKNEGGKIIAGGERTSQEQT<br>LQQGFYLQPTLITGNNSMRFFREEIFGPVIGITTFKDEAEAL<br>ALANDTEFGLGAGLWTRDSNRAYRMGRAIKAGRVWTNCYHLY<br>PAHAAFGGYKNSGVGRETHKVALSHYQQVKNLLVSYDAKPQG<br>LF* |
| A0A2S0MQS8 | ALDH-077 | MNVQNQLNLSAALDTSHLGVGMPFKARYGNFINGEFVEPKSG<br>RYFENTTPITGEVVCEVARSDAKDVEAALDAAHAAFPAWGRT<br>SLAERSNMLMKIADVIEANTKLLAAAETIDNGKPIRESIAAD<br>ITLAVDHFRYFGGVIRAEEGSVAEIDANTYAYHINEPLGVVG<br>QIIPWNFPLLMATWKIAPALAAGNCIVLKPAEQTPASIMVLM<br>ELIGDLIPPGVLNVVNGFGLEAGKPLASSNRIAKIAFTGETT<br>TGRLIMQYASQNIIPVTLELGGKSPNIFFEDIMAADDAYFDK<br>TLEGFTMFALNQGEVCTCPSRALIQESIYDAFIERAIERVGK<br>VRQGNPLSMDTMIGAQASTEQMEKISSYLQLGRDEGADVLIG<br>GDIAALNGGLENGNFIQPTVFRGHNKMRIFQEEIFGPVLSVT<br>TFKDEAEALEIANDTLYGLGAGVWTRDGSRAFRMGRAIQAGR<br>VWTNCYHAYPAHAAFGGYKQSGIGRENHKMMLDHYRQTKNLL<br>VSYDANALGFF* |
| A0A849HPN9 | ALDH-078 | MLDAAIAPLAGSAVAGFSFKARYDNFIGGKFTAPLAGQYFDN<br>VSPITGQVFTQAARSTEADINLALDAAHAAADAWGKTSPAAR<br>ALILNRIADKMEQNLDLLAYAETVDNGKPLRETTFADIPLAI<br>DHMRYFAACVRAQEGALSQIDEDTVAYHFHEPLGVVGQIIPW<br>NFPILMAVWKLAPALAAGNCVVLKPAEQTPVSIMVLAELIGD<br>MLPPGVLNIVNGFGLEAGKPLASSNRIAKIAFTGETTTGRLI<br>MQYASHNLIPVTLELGGKSPNIFFEDVMAQDDDFLDKAVEGL<br>VMFAFNQGEVCTCPSRALIQESIYDRFMEKVLKRVHAIKQGN |

| | | |
|---|---|---|
| | | PLDMSTMVGAQASSEQVQKILSYIDIGKQEGAKLLTGGVRPA LEGGLSDGFYIQPTMFLGNNKMRLFQEEIFGPVLAVTTFKDE AEALELANDTLYGLGAGVWSRDGSRAYRMGRGIKAGRVWTNC YHAYPAHAAFGGYKQSGIGRENHKMMLDHYQQTKNLLVSYSP KALGFF* |
| A0A4R6PRX6 | ALDH-079 | MIYGKPGSDTGIVSYAARYDNFIGGEWVAPVEGRYIDNTSPV DGEAFCAVARSGAADIDLALTAARRAAPGWAATSPADRAGIL LAIADRIENSIEPLAVAETWDNGKPVRETLHTDLPLAVDHFR YFAGVLRGQEGTISTVDSETVAYHLSGPLGVVGHILPWNYPI LTAAWELAPALAAGNCVVLKPAEQTPASLLLVVELIADLLPD GVLNVVNGFGAEAGVPLARSPQLDKLTFTGDTTTARLILRHA AENIVPVSLRLGGKCANVFTADVLAADDTFLDSAVEGFVMFV LNQGEVGTCPSRALIHSSIYDEFLARCIARTEAVASGHPLDT ETMIGAQAGNDQYERILSYFDIGRAEGARVLTGGEARKVDGL PGGYYIEPTVFEGDNSMRIFQEEIFGPVLGVTRYDTLDEAVR VTDDTSHRLGAGVWTRDLGTAHRFAGAVGTRRVWANCYHPYP AQAAGENTAMMLGRYQRSKNLLVRHSTTTPGMF* |
| A0A251X6G7 | ALDH-080 | MLYAQPNQDGAKVHFKPRYGNYINGQWQAPVKGEYFENPTPV TGKNFCEIARSTAEDIELALDAAHAAKTAWGRTPVAQRANVL LKIADRMEANLEALAVAETWDNGKPIRETLAADVPLAIDHFR YFAGCIRAQEGTLSEISPDLVAYHFHEPLGVVGQIIPWNFPL LMAAWKIAPALAAGNCIVLKPAEHTPVSMLIWAEIVGDLLPP GVLNIVNGFGLEAGKPLVTSSRIAKIAFTGETTTGRLIMQYA SQNIIPVTLELGGKSPNIFFEDVMSQDDAFLNKALEGFTLFA LNQGEICTCPSRALVQASIYDAFIEKAIARVQKIRRGNPLDT DTMIGAQASTEQMEKIMSYIDVGQQEGATLLTGGQRCQLDGE LSSGYYIEPTVFAGHNRMRIFQEEIFGPVLSVAKFHDEEEAL HLANDTLYGLGAGVWSRDGSRAFRMGREIQAGRVWTNCYHLY PAHAAFGGYKQSGIGRETHKMMLSHYQQTKNLLVSYDPNALG FF* |
| UPI0005427CB8 | ALDH-081 | MATWKLAPALAAGNCVVLKPAEQTPASIMVLMELIGDLIPPG VVNVVNGFGLEAGKPLASNSRIAKVAFTGETTTGRLIMQYAS QNLIPVTLELGGKSPNIFFEDVMAKDDAFFDKALEGFTLFAL NQGEVCTCPSRALIQESIYDAFMERALKRVASIKQGHPLDPA TMIGAQVSTEQMEKILSYIDIGRQEGAECLIGGGRASIPGEL SDGYYIQPTVLKGQNNMRIFQEEIFGPVVAVTTFRDREEALA IANDTLYGLGAGVWTRDVNTAYRVGREIQAGRVWTNCYHAYP AHAAFGGYKSSGIGRENHRMMLSHYQQTKNLLVSYSPKALGF F* |
| A0A395JK05 | ALDH-082 | MQYQNPNTEGAIVSFKQRYENYIGGEWVKPVNGKYFDNISPV NGKAFCEIPRSDEQDINLALDAAHEAKDAWGSTSVSERSNIL LKIADRMEQHLEALAVAETWDNGKAIRETLNADVPLCIDHLR YYAGCLRAQEGSISELDNNTVAYHFHEPLGVVGQIIPWNFPL LMAIWNLTPAIAAGNCVVLKPAEQTPATILMLMELVGDIMPK GVLNVVNGFGEEAGNALATSDRIAKIAFTGSTPVGQHILRCA AASLIPSSVELGGKSPNVFFADILNQEDEFVSKCVEGAVLAF FNQGEVCTCPSRLLIEESIYDEFIGMVIDRASQIKRGSPLDT DVMVGAQVSQEQFDRILDYIKIGLEEGAEVITGGNAANIGDG CEAGYYVEPTLLKGSNNMRVFQEEIFGPVVSVTTFKTPEEAL EIANDSDFGLGAGVWTRDMNLSYRMGRGIQAGRVWTNCYHLY |

| | | |
|---|---|---|
| | | PAHAAFGGYKKSGIGRYTHKVALEHYQQTKNLLVSYDTNPLG<br>FF* |
| A0A2E2EWH7 | ALDH-083 | MSYSKPQFKDKYSNFIDGKFVPPVGGDYFENTSPIDGSLIAK<br>YPRSQKEDVENAVAAANAAKEAWGNTSVTERAALLNKVADII<br>EENLEEFALVETCDNGKPIRETLNADIPLCVDHWRYFAACIR<br>AEEGSATELDANTLSMNIKEPLGVVGQIIPWNFPLLMLSWKL<br>PPALATGNCVVLKPAEQTPSSATLLMEKIADVFPPGVINIIH<br>GFGPEAGKPLASSSKIDKVAFTGETTTGQLIMQYASKNLNPV<br>TMELGGKSPNIFFNSVMDADDEYLDKAIEGAVLFAFNQGEVC<br>TCPSRILVQEDIYDKFMERVIARTEAIVQTSPYSTDCMVGAQ<br>ASNDQYEKIQSYIKIGKDEGAKVLCGGEANREGDLANGYYIK<br>PTILEGHNKMRVFQEEIFGPVVCVTKFKDEAEALEIANDTLY<br>GLGAGVWTRDAHQLYQIPRAIKAGRVWVNCYHAYPAHAPFGG<br>YKKSGFGRENHQMMMSHYRQTKNMLISYDKNKLGFF* |
| Q9ZA11 | ALDH-084 | MLYAAPGTADAIFAFKPRYDNFIDGTWQPPVRGEYFDNVTPI<br>TGKVFCKAARSTEEDITLALDAAHAAADRWGRTTAAERALIL<br>NRIADRLQDNLETLAYVESIDNGKPIRETLAADIPLAIDHFR<br>YFAACIRAQEGSLSQIDETTIAYHFNEPLGVVGQIIPWNFPI<br>LMATWKLAPALAAGNCIVLKPAEQTPISILVLTELIADLLPP<br>GVLNVVNGFGLEAGKPLASSKRIAKIAFTGETATGRLIMQYA<br>SQNLIPVTLELGGKSPNVFFDDIASADDSFFDKAVEGFVMFA<br>LNQGEICTCPSRALIHESLYDRFIERALARVAVIKQGSPLDT<br>ETMIGAQASTEQMDKILSYMDIGREEGATVLAGGARAELGGE<br>LDGGYYVQPTVFKGNNSMRIFQEEIFGPVVAVTTFKDEDEAL<br>HLANDTHYGLGSGVWTRDGNRAFRFGRGIKAGRVWTNCYHLY<br>PAHAAFGGYKQSGIGRENHHMMLDHYQQTKNLLVSYDPKAMG<br>FF* |
| A0A3B0T5Y3 | ALDH-085 | MEAQKIEIPKLFKEQYGHFINNEWVAPASGEYFDNNCPIDNS<br>LIFKTARGNQEDVEKAITAAANAFPAWSKTPAPARSAVLLKI<br>AQVMEDNIELLAKAEVVDKGKPMREALLVDLPVAIDNFRYFA<br>GVIRAEEGGTSMQDEHTLQINQPEALGPVGAIIAWNFPLLLF<br>AWKVAPAIAAGCTIVVKTAEQTPAGANLLMDLLKEADAVPPG<br>VINVVTGFGLEAGKPLAMSPNIAKISFTGETATGRLILQYAA<br>ENIIPATMELGGKSPNIFMPSIADKDDDFFDKCIEGAVMFAV<br>NQGEACTCPSRLFVHEDIYDKFMGRVVDRTKAIVQGDPFEMT<br>TMMGAQASKDQYDKILKYIEIGKQEGAKVLCGGDANSDGACD<br>NGYFIQPTILEGNNKMKVFQEEIFGPVVCVTKFKTTEEVLEM<br>ANDSIYGLAAAVWTRDAHEAFAMPRAIEAGRVWVNCYHAYPT<br>HAAFGGFKKSGFGRENHKMALASFQRTKNIFQSNAQQKLGFF<br>* |
| A0A3B1AP81 | ALDH-086 | MIYSKPGSEGTLLSFKDSYDNFIGGNWVAPVEGAYFDNISPV<br>DGKVYCRIARSTAADIEQALDAAHAAKDDWGKTSVTERANIL<br>LQIADRIEQNIEQLALTETWDNGKPVRETLAADVPLAVDHFR<br>YFAGCIRAQEGGIGEIDEATVSYQYHEPLGVVGQIIPWNFPL<br>LMATWKLAPALAAGNCVVLKPAEQTPVSICLLLEIIGDLLPP<br>GVLNIVQGYGEEAGKALATSNRIAKIAFTGSTAVGQQIMAYA<br>SQNIIPVTLELGGKSPNIFFADVMQQSPDYIDKAIEGLALGF<br>FNQGEVCTCPSRALIQEDIYDDFIAQVIGRVRQIKQGNPLDT<br>ETMIGAQASQEQLDKILGYMRIAEQEGAQCLIGGQRNTSGEL<br>TNGYYVEPTLYKGDNSMRIFQEEIFGPVLAITTFKDEAEAIA<br>IANDTEYGLGAGVWTMDMNRAYRLGRAIQAGRIWTNCYHAYP |

| | | |
|---|---|---|
| | | AHAAFGGYKKSGIGRETHKVMLEHYQQTKNVLVSYSTQALGF<br>F* |
| A0A443K8I0 | ALDH-087 | MPEDQILAETAFKSSPFKARYENYIGGKWMAPKSGRYMQNIS<br>PVTGHVVCEVPRSDAADVEAALDAAHAARRAWGRASNAERAG<br>ILNRVADRLEQNLSAIALAETWDNGKPLRETMNADIPLAIDH<br>FRYFAGVIRAEEGSIGEIDHDTVAYHYKEPLGVVGQIIPWNF<br>PILMAAWKIAPALAAGNCIVMKPAEQTPASILVLLEFIEDLL<br>PPGVLNIVNGTGIEVGAPLASSNRIAKIAFTGSTPVGKSIMK<br>AASEHVTNITLELGGKSPNIFFADVMAEEDDFLDKALEGFSF<br>FALNQGEICTCPSRALVQESIYDRFMEKALKRVEAIQAGDPR<br>ATGTMIGAQASKQQFDKILSYMEIGLNEGAKILTGGAAQSFS<br>GDIEGGYYIKPTIFEGNNRMRVFQEEIFGPVVSVTTFKTLED<br>ALEIANDTVFGLGAGVWSRDMNTAYRAGRGIEAGRVWTNCYH<br>LYPAGAAFGGYKQSGIGRENHKMMLDHYQETKNVLVSYSPKK<br>LGFF* |
| A0A4P6V3E3 | ALDH-088 | MNVMTNIAAEYTSPFKQRYDNFIGGRFVPPVDGRYFDNITPI<br>TGQKVCEVARSGAADVEMALDAAHAAKGAWGRTSAAERSSVL<br>LKIADRLEDNLDTLAQAETWDNGKPIRETTAADIPLAIDHFR<br>YFAGVLRAQEGNMSEIDADTVAYHFHEPLGVVGQIIPWNFSI<br>LMAAWKLAPALAAGNCVVLKPAEQTPAAIMVLAELVADLLPD<br>GVLNIVNGFGPEVGGPLAESDRIAKIAFTGSTETGRIIMKAA<br>TKNLIPVTLELGGKSPNIFFSDVMAEDDAFLDKAVEGFVLFA<br>FNQGEVCTCPSRALIQEDIYEAFIERCIARVEAIVQGDPRDM<br>NTMVGAQASREQQDKIKSYLTIGVEEGAEVLTGGAEARFDGE<br>IANGFYIQPTILKGHNKMRVFQEEIFGPVVSVTTFKDEEEAL<br>AIANDTMYGLGAGVWSRDANRCYRFGRAIEAGRVWVNNYHAY<br>PAHAAFGGYKQSGIGRETHKMMLDHYQQTKNILMSYNPNKLG<br>FF* |
| A0A7Y9PEX6 | ALDH-089 | MATIPQIHPGKYGYLVEFKTRYGNFINGKWAAPEGGNYFENI<br>SPVTGKPFCEVPRSTAADIERALDAAHAAKGAWGKTSTAARA<br>RILEQIATRIDENLELLATVETWDNGKPIRETTAADIPLAAD<br>HFRYFAAAVRAQEGSISEIDGDTVAYHYHEPLGVIGQIIPWN<br>FPILMAAWKLAPALAAGNCVVLKPAEQTPVSILVLMELIHDL<br>LPAGVLNIVNGFGLEAGKPLASNARVNKVAFTGETTTGRLIM<br>QYASQNIIPVTLELGGKSPNIFFADIMSADDALFDKALEGFV<br>LFAFNQGEVCTAPSRALIQASIYDAFLERAIARVKKIKRGNP<br>LDPATMIGAQASSEQLEKILSYIDIGKQEGAKLLTGGKRAVQ<br>EGDLADGYYVEPTVFEGHNKMRIFQEEIFGPVLSVTKFENDD<br>EALAIANDTLYGLGAGVWTRDIHRAYRFGRGIQAGRVWTNCY<br>HLYPAGAAFGGYKASGIGRENHKMMLDHYQQTKNQLISYNPN<br>ALGFF* |
| A0A023CJ10 | ALDH-090 | MLYAQPGQPGALVTFKQRYENFIGGKWVPPVDGEYFENISPI<br>TGKPYCEVPRSKAADIELALDAAHAAKDAWGRTSVTERARIL<br>NKIADRMEENLEMLAVAETWENGKPIRETLAADIPLAIDHFR<br>YFAGCIRAQEGTLAEIDENTVAYHFKEPLGVVGQIIPWNFPI<br>LMAAWKLAPALAAGNCVVLKPAEQTPTSILVLMELIQDLLPP<br>GVVNIVNGFGLEAGKPLASNPRVAKVAFTGETTTGRLIMQYA<br>SQNIVPVTLELGGKSPNIFFADVMEQDDEFLDKALEGFTMFA<br>LNQGEVCTCPSRALIEESIYDAFMERALERVKQIKQGNPLDT<br>ETMIGAQASSEQLEKILSYIDIGKQEGAELLIGGERNYLEGD<br>LRDGYYVKPTIFKGHNKMRIFQEEIFGPVLAVTTFKDKDEAL |

| | | |
|---|---|---|
| | | AIANETLYGLGAGVWTRDINTAYRFGRGIQAGRVWTNCYHVY<br>PAHAAFGGYKMSGIGRETHKMMLDHYQQTKNLLVSYSPKKLG<br>LF* |
| A0A0J0V889 | ALDH-091 | MIYAQPGQPGALVTLKKRYENFIGGKWVPPVDGEYFENITPI<br>TGQPYCEVPRSKAADIELALDAAHAAKDAWGRTSPAERARLL<br>NKIADRMEEHLEMLAVAETWENGKPIRETLAADIPLAIDHFR<br>YFASCIRAQEGTISEIDHDTVAYHFKEPLGVVGQIIPWNFPI<br>LMAAWKLAPALAAGNCVVLKPAEQTPTSILVLIELIEDLLPP<br>GVVNVVNGFGLEAGKPLASNPRVAKVAFTGETTTGRLIMQYA<br>SQNIVPVTLELGGKSPNIFFADVMDNNDEFLDKALEGFTMFA<br>LNQGEVCTCPSRALIHESIYDAFMERALERVKQIKQGNPLDT<br>ETMIGAQASSEQLEKILSYIDIGKQEGAELLIGGERNMLEGE<br>LAGGYYVKPTIFKGHNKMRIFQEEIFGPVLAVTTFKDHDEAL<br>AIANETLYGLGAGVWTRDINTAYRFGRGIQAGRVWTNCYHVY<br>PAHAAFGGYKMSGIGRETHKMMLDHYQQTKNLLVSYSPKKLG<br>LF* |
| A0A0K6GMR6 | ALDH-092 | MIYAQPGQPGSLITFKKRYENFIGGEWVPPIDGEYFENISPV<br>TGQVYCEVPRSKAVDIELALDAAHAAKEAWGRTSVAERARLL<br>NKIADRMEENLDMLAVAETWENGKPIRETRAADIPLAIDHFR<br>YFAGCIRAEEGTLAELDNDTVAYHFKEPLGVVGQIIPWNFPI<br>LMAAWKLAPALAAGNCVVLKPAEQTPTSILVLIELIQDLLPK<br>GVVNVVNGFGLEAGKPLASNPRIAKVAFTGETTTGRLIMQYA<br>SQNLIPVTLELGGKSPNIFFADVAAKDDEFFDKALEGFTMFA<br>LNQGEVCTCPSRALIEESIYDIFMERALERVKQIKQGNPLDT<br>STMIGAQASSEQLEKILSYIDIGKQEGAELLVGGERNMLEGI<br>LQGGYYVKPTVFKGHNRMRIFQEEIFGPVVSVTTFKDQEEAL<br>SIANDTLYGLGAGVWTRDMNTAYRFGRGIQAGRVWTNCYHAY<br>PAHAAFGGYKMSGIGRETHKMMLDHYQQTKNLLVSYSPKKLG<br>FF* |
| A0A142D425 | ALDH-093 | MIYAQPGQPGALVTFKKRYENFIGGKWVPPVDGEYFENITPI<br>TGQPYCEVPRSKAADIELALDAAHAAKDAWGRTSPAERARLL<br>NKIADRMEEHLEMLAVAETWENGKPIRETLAADIPLAIDHFR<br>YFASCIRAQEGTISEIDHDTVAYHFKEPLGVVGQIIPWNFPI<br>LMAAWKLAPALAAGNCVVLKPAEQTPTSILVLIELIEDLLPP<br>GVVNIVNGFGLEAGKPLASNPRVAKVAFTGETTTGRLIMQYA<br>SQNIVPVTLELGGKSPNIFFADVMDNDDEFLDKALEGFTMFA<br>LNQGEVCTCPSRALIHESIYDVFMERALERVKQIKQGNPLDT<br>ETMIGAQASSEQLEKILSYIDIGKQEGAELLIGGERNILEGE<br>LAGGYYVKPTIFKGHNKMRIFQEEIFGPVLAVTTFKDHDEAL<br>AIANETLYGLGAGVWTRDINTAYRFGRGIQAGRVWTNCYHVY<br>PAHAAFGGYKMSGIGRETHKMMLDHYQQTKNLLVSYSPKKLG<br>LF* |
| UPI0007A942CE | ALDH-094 | MIYAQPGQPGSLVTFKKRYENFIGGKWVPPVDGEYFENITPI<br>TGQPYCEVPRSKAADIELALDAAHAAKEEWGRTSPAKRARLL<br>NKIADRMEENLERLAVAETWENGKPIRETLAADIPLAIDHFR<br>YFASCIRAQEGTISEIDHDTVAYHFKEPLGVVGQIIPWNFPI<br>LMAAWKLAPALAAGNCVVLKPAEQTPTSILVLIELIEDLLPP<br>GVVNVVNGFGLEAGKPLASNPRVAKVAFTGETTTGRLIMQYA<br>SQNIVPVTLELGGKSPNIFFPDVMDKDDEFLDKALEGLTMFA<br>LNQGEVCTCPSRAIIHESIYDAFMERALERVKQIKQGNPLDT<br>ETMIGAQASSEQLEKILSYIDIGKQEGAELLIGGERNMLEGE |

| | | |
|---|---|---|
| | | FAGGYYVKPTIFKGHNKMRIFQEEIFGPVLAVTTFKDNDEAL AIANETLYGLGAGVWTRDINTAYRFGRGIQAGRVWTNCYHVY PAHAAFGGYKMSGIGRETHKMMLDHYQQTKNLLVSYSPKKLG LF* |
| A0A178T558 | ALDH-095 | MIYAQPGQPGSLITFKKRYENFIGGEWVPPIDGEYFENISPV TGQVYCEVPRSKAADIELALDAAHAAKEAWGRTSVAERARLL NKIADRMEENLDMLAVAETWENGKPIRETRAADIPLAIDHFR YFAGCIRAEEGTLAEIDADTVAYHFKEPLGVVGQIIPWNFPI LMAAWKLAPALAAGNCVVLKPAEQTPTSILVLIELIQDLLPK GIVNVVNGFGLEAGKPLASNPRIAKVAFTGETTTGRLIMQYA SQNLIPVTLELGGKSPNIFFADVAAKDDEFFDKALEGFTMFA LNQGEVCTCPSRALIEESIYDVFMERALERVKQIKQGNPLDT STMIGAQASSEQLEKILSYIDIGKQEGAELLIGGERNMLEGD LQGGYYMKPTVFKGHNRMRIFQEEIFGPVVSVTTFKDQDEAL SIANDTLYGLGAGIWTRDINTAYRFGRGIQAGRVWTNCYHTY PAHAAFGGYKMSGIGRETHKMMLEHYQQTKNLLVSYSPKKLG FF* |
| A0A1I0TE05 | ALDH-096 | MIYAQPGQPGSLITFKKRYENFIGGKWVPPVDGEYFENISPV TGKPYCEVPRSKAADIELALDAAHAAKDAWGRTSPAERARIL NKIADRMEENLEMLAVAETWENGKPIRETLNADIPLAIDHFR YFASCIRAEEGTLAEIDHDTVAYHFKEPLGVVGQIIPWNFPL LMAAWKLAPALAAGNCVVLKPAEQTPTTILVLMELIEDLLPP GVVNIVNGFGLEAGKPLASSSRIAKIAFTGETTTGRLIMQYA SQNIIPVTLELGGKSPNIFFEDVAAKDDEFFDKALEGFTLFA LNQGEVCTCPSRALIQESIYEKFMERALERVKQIKQGNPLDT ETMIGAQASSEQLEKILSYIDIGKQEGAELLIGGERNILEGD LKDGYYVKPTVFKGHNKMRIFQEEIFGPVVSVTTFKDIDEAL EIANETLYGLGAGVWTRDINTAYRVGRGIQAGRVWTNCYHMY PAHAAFGGYKLSGFGRETHKMMLEHYQQTKNLLVSYSPKKLG LF* |
| UPI0009ADBF94 | ALDH-097 | MIYAQPGQPGALVTFKKRYENFIGGKWVPPVDGEYFENITPI TGQPYCEVPRSKAADIELALDAAHAAKDAWGRTSPAERARLL NKIADRMEENLEMLAVAETWENGKPIRETLAADIPLAIDHFR YFASCIRAQEGTISEIDHDTVAYHFKEPLGVVGQIIPWNFPI LMAAWKLAPALAAGNCVVLKPAEQTPTSILVLIELIEDLLPP GVVNIVNGFGLEAGKPLASNPRVAKVAFTGETTTGRLIMQYA SQNIVPVTLELGGKSPNIFFADVMDKDDEFLDKALEGFTMFA LNQGEVCTCPSRALIHESIYDAFMERALERVKQIKQGNPLDT ETMIGAQASSEQLEKILSYIDIGKQEGAELLIGGERNMLEGE LAGGYYVKPTVFKGHNKMRIFQEEIFGPVLAVTTFKDHDEAL SIANETLYGLGAGVWTRDINTAYRFGRGIQAGRVWTNCYHIY PAHAAFGGYKMSGIGRETHKMMLDHYQQTKNLLVSYSPKKLG LF* |
| A0A1V9B5J1 | ALDH-098 | MIYAQPGQSGALVTFKKRYENFIGGKWVPPMDGEYFENITPI TGQPYCEVPRSKAADIELALDAAHAAKEEWGRTSPAKRARLL NKIADRMEENLERLAVAETWENGKPIRETLAADIPLAIDHFR YFASCIRAQEGTISEIDHDTVAYHFKEPLGVVGQIIPWNFPI LMAAWKLAPALAAGNCVVLKPAEQTPTSILVLIELIEDLLPP GVVNVVNGFGLEAGKPLASNPRVAKVAFTGETTTGRLIMQYA SQNIVPVTLELGGKSPNIFFPDVMDKDDEFLDKALEGLTMFA LNQGEVCTCPSRALIHESIYDAFMERALERVKQIKQGNPLDT |

| | | |
|---|---|---|
| | | ETMIGAQASSEQLEKILSYIDIGKQEGAALLIGGERNMLEGE<br>LAGGYYVKPTIFKGHNKMRIFQEEIFGPVLAVTTFKDNDEAL<br>AIANETLYGLGAGVWTRDINTAYRFGRGIQAGRVWTNCYHVY<br>PAHAAFGGYKMSGIGRETHKMMLDHYQQTKNLLVSYSPKKLG<br>LF* |
| UPI00017E6B4A | ALDH-099 | MIYAQPGQPGALVTFKKRYENFIGGKWVPPVDGEYFENITPI<br>TGQPYCEVPRSKAADIELALDAAHAAKEEWGRTSPAKRARLL<br>NKIADRMEENLEMLAVAETWENGKPIRETLAADIPLAIDHFR<br>YFASCIRAQEGTISEIDHNTVAYHFKEPLGVVGQIIPWNFPI<br>LMAAWKLAPALAAGNCVVLKPAEQTPTSILVLIELIEDLLPP<br>GVVNVVNGFGLEAGKPLASNPRVAKVAFTGETTTGRLIMQYA<br>SQNIVPVTLELGGKSPNIFFADVMEKDDDFLDKALEGFTMFA<br>LNQGEVCTCPSRALIHESIYDAFMERALERVKQIKQGNPLDT<br>ETMIGAQASSEQLEKILSYIDIGKQEGAELLVGGERNMLEGE<br>LAGGYYVKPTIFKGHNKMRIFQEEIFGPVLAVTTFKDNDEAL<br>AIANETLYGLGAGVWTRDINTAYRFGRGIQAGRVWTNCYHVY<br>PAHAAFGGYKMSGIGRETHKMMLDHYQQTKNMLVSYSPKKLG<br>LF* |
| UPI000A26C654 | ALDH-100 | MIYAQPGQPGSLITFKKRYENFIGGEWVPPIDGEYFENISPV<br>TGQVYCEVPRSKAADIELALDAAHAAKETWGRTSVAERARLL<br>NKIADRMEENLDMLAVAETWENGKPIRETRVADIPLAIDHFR<br>YFAGCIRAEEGTLAELDDDTVAYHFKEPLGVVGQIIPWNFPI<br>LMAAWKLAPALAAGNCVVLKPAEQTPTSILVLIELIQDLLPK<br>GVVNVVNGFGLEAGKPLASNPRITKVAFTGETTTGRLIMQYA<br>SQNLIPVTLELGGKSPNIFFADVVAKDDEFFDKALEGFTMFA<br>LNQGEVCTCPSRALIEESIYDVFMERALERVKQIKQGNPLDT<br>STMIGAQASSEQLEKILSYIDIGKQEGAELLIGGERNMLEGD<br>LQGGYYVKPTVFKGHNRMRIFQEEIFGPVVSVTTFKDKDEAL<br>AIANDTLYGLGAGVWTRDINTAYRFGRGIQAGRVWTNCYHAY<br>PAHAAFGGYKMSGIGRETHKMMLEHYQQTKNLLVSYSPKKLG<br>FF* |
| A0A2M9T1Z9 | ALDH-101 | MIYAQPGQLGALVTFKKRYENFIGGKWVPPVDGEYFENITPI<br>TGQPYCEVPRSKAADIELALDAAHAAKDAWGRTSPAERARLL<br>NKIADRMEENLEMLAVAETWENGKPIRETLAADIPLAIDHFR<br>YFASCIRAQEGTISEIDHDTVAYHFKEPLGVVGQIIPWNFPI<br>LMAAWKLAPALAAGNCVVLKPAEQTPTSILVLIELIEDLLPP<br>GVVNIVNGFGLEAGKPLASNPRVAKVAFTGETTTGRLIMQYA<br>SQNIVPVTLELGGKSPNIFFADVMDKDDDFLDKALEGFTMFA<br>LNQGEVCTCPSRALIHESIYDAFMERALERVKQIKQGNPLDT<br>ETMIGAQASSEQLEKILSYIDIGKQEGAELLIGGERNMLEGE<br>LAGGYYVKPTVFKGHNKMRIFQEEIFGPVLAVTTFKDHDEAL<br>SIANETLYGLGAGVWTRDINTAYRFGRGIQAGRVWTNCYHVY<br>PAHAAFGGYKMSGIGRETHKMMLDHYQQTKNLLVSYSPKKLG<br>LF* |
| A0A327YHK1 | ALDH-102 | MIYAQPGQPGSLITFKKRYENFIGGKWVPPVDGEYFENISPV<br>TGLAYCEVPRSKAADIELALDAAHAAKEAWGRTSAAERARIL<br>NKIADRMEENLEMLAVAETWENGKPIRETLAADIPLAIDHFR<br>YFAGCIRAEEGSLAEIDHDTVAYHFKEPLGVVGQIIPWNFPI<br>LMAAWKLAPALAAGNCVVLKPAEQTPTSILVLMELIQDLLPP<br>GVVNIVNGFGLEAGKPLASNSRIAKIAFTGETTTGRLIMQYA<br>SQNIIPVTLELGGKSPNIFFEDVMEKDDEFFDKALEGFTMFA |

| | | |
|---|---|---|
| | | LNQGEICTCPSRALIQESIYDAFIERALERVKQIKQGNPLDT ETMIGAQASSEQLEKILSYIDIGKQEGAELLIGGERNVLEGD LSGGYYVKPTVFKGHNKMRIFQEEIFGPVVAVTTFKDQEEAL AIANETLYGLGAGVWTRNINTAYRVGRGIQAGRVWTNCYHAY PAHAAFGGYKLSGIGRETHKMMLEHYQQTKNLLVSYSPKKLG FF* |
| A0A4Q1RV81 | ALDH-103 | MIYAQPGQPGALVTFKKRYENFIGGKWVPPVDGEYFENITPI TGQPYCEVPRSKAADIELALDAAHAAKDAWGRTSPAERARLL NKIADRMEEHLEMLAVAETWENGKPIRETLAADIPLAIDHFR YFASCIRAQEGTISEIDHDTVAYHFKEPLGVVGQIIPWNFPI LMAAWKLAPALAAGNCVVLKPAEQTPTSILVLIELIEDLLPP GVVNIVNGFGLEAGKPLASNPRVAKVAFTGETTTGRLIMQYA SQNIVPVTLELGGKSPNIFFADVMDKDDDFLDKALEGFTMFA LNQGEVCTCPSRALIHESIYDAFMERALERVKQIKQGNPLDT ETMIGAQASSEQLEKILSYIDIGKQEGAELLIGGERNILEGE LAGGYYVKPTIFKGHNKMRIFQEEIFGPVLAVTTFKDHDEAL AIANETLYGLGAGVWTRDINTAYRFGRGIQAGRVWTNCYHVY PAHAAFGGYKMSGIGRETHKMMLDHYQQTKNLLVSYSPKKLG LF* |
| A0A4R1QFD0 | ALDH-104 | MIYAQPGHSNSLITFKKRYENFIGGKWVPPVDGEYFENISPV TGEVYCEVPRSKAADIELALDAAHAAKEAWGRTSVAERARIL NKIADRMEENLEMLAVAETWENGKPIRETLAADIPLAIDHFR YFASCIRAHEGSLAEIDHDTVAYHFKEPLGVVGQIIPWNFPI LMAVWKLAPALAAGNCAVLKPAEQTPTSVLVLMELIEDLLPP GVVNIVNGFGLEAGKPLASSNRIAKVAFTGETTTGRLIMQYA SQNIIPVTLELGGKSPNIFFEDVLEKDDEFFDKALEGFTMFA LNQGEVCTCPSRALIQESIYDEFMERALKRVQQIKQGNPLDT ETMIGAQASSEQLEKILAYIDIGKQEGAELLIGGERNFLEGD LRNGYYVKPTVFKGHNRMRIFQEEIFGPVVAVTTFKDAEEAL EIANDTLYGLGAGVWTRDINTAYRFGRGIQAGRVWTNCYHAY PAHAAFGGYKMSGIGRETHKMMLEHYQQTKNLLVSYSPKKLG FF* |
| A0A6G9J022 | ALDH-105 | MIYAQPGQPGSLITFKKRYENFIGGKWVPPVDGEYFENISPV TGQAYCEVPRSQAADIELALDAAHAAKDAWGRTSPAERARIL NKIADRMEENLEMLAVAETWENGKPIRETLNADIPLAIDHFR YFAGCIRAEEGTLSEIDHDTVAYHFKEPLGVVGQIIPWNFPL LMAAWKLAPALAAGNCVVLKPAEQTPTSILVLMELIQDILPP GVVNVVNGFGVEAGKPLASSSRIAKIAFTGETTTGRLIMQYA SQNIIPVTLELGGKSPNIFFEDVAAKDDEFFDKAIEGFTLFA LNQGEICTCPSRALIHESIYDKFMERALERVKQIKQGNPLDT ETMIGAQASSEQLEKILSYIDIGKQEGAELLAGGERNILEGD LKDGYYVKPTVFKGHNKMRIFQEEIFGPVVAVTTFKDMDEAL EIANDTLYGLGAGVWTRDINTAYRVGRGIQAGRVWTNCYHMY PAHAAFGGYKQSGFGRETHKMMLEHYQQTKNLLVSYSPKKLG LF* |
| UPI00050099C5 | ALDH-106 | MIYAQPGQPGALVTFKKRYENFIGGKWVPPVDGEYFENITPI TGQPYCEVPRSKAADIELALDAAHAAKEAWGRTSPAERARLL NKIADRMEENLEMLAVAETWENGKPIRETLAADIPLAIDHFR YFASCIRTQEGTISEIDHDTVAYHFKEPLGVVGQIIPWNFPI LMAAWKLAPALAAGNCVVLKPAEQTPTSILVLIELIEDLLPP GVVNVVNGFGLEAGKPLASNPRVAKVAFTGETTTGRLIMQYA |

| | | |
|---|---|---|
| | | SQNIVPVTLELGGKSPNIFFPDVMDKDDEFLDKALEGLTMFA LNQGEVCTCPSRALIHESIYDAFMERALERVKQIKQGNPLDT ETMIGAQASSEQLEKILSYIDIGKQEGAELLIGGERNMLEGE LAGGYYVKPTIFKGHNKMRIFQEEIFGPVLAVTTFKDNDEAL AIANETLYGLGAGVWTRDINTAYRFGRGIQAGRVWTNCYHIY PAHAAFGGYKMSGIGRETHKMMLDHYQQTKNLLVSYSPRKLG LF* |
| A0A7U3YC36 | ALDH-107 | MIYAQPGQPGSLITFKKRYENFIGGKWVPPVDGEYFENISPV TGKPYCEVPRSKAADIELALDAAHAAKDAWGRTSPAERARIL NKIADRIEENLEMLAVAETWENGKPIRETLNADIPLAIDHFR YFAGCIRAEEGTLAEIDNDTVAYHFKEPLGVVGQIIPWNFPL LMATWKLAPALAAGNCVVLKPAEQTPTSILVLMELIEDLLPP GVVNIVNGFGLEAGKPLASSNRIAKIAFTGETTTGRLIMQYA SQNIIPVTLELGGKSPNIFFEDVAAKDDEFFDKAIEGFTLFA LNQGEICTCPSRALIQESIYDQFIERALERVKQIKQGNPLDT ETMIGAQASSEQLEKILSYIDIGKQEGAELLIGGERNFLEGD LRDGYYVKPTVFKGHNKMRIFQEEIFGPVVSVTTFKDVNEAL EIANETLYGLGAGVWTRDINTAYRVGRGIQAGRVWTNCYHIY PAHAAFGGYKLSGFGRETHKMMLEHYQQTKNLLVSYSPKKLG LF* |
| A0A7V9YZI5 | ALDH-108 | MIYAQPGKPGSLITFKKRYENFIGGKWVPPVDGEYFENISPV TGEVYCEVPRSKAADIELALDAAHAAKEAWGRTSPAERARIL NKIADRMEENLEMLAVAETWENGKPIRETLNADIPLAIDHFR YFAGCIRAEEGSLAEIDHDTVAYHFKEPIGVVGQIIPWNFPL LMGVWKLAPALAAGNCVVLKPAEQTPTSILVLMELIQDLLPP GVVNIVNGFGLEAGKPLASSSRIGKIAFTGETTTGRLIMQYA SQNIIPVTLELGGKSPNIFFEDVMDKDDEFFDKALEGFTMFA LNQGEICTCPSRALIQESIYDAFIERALERVKQIKQGNPLDT ETMIGAQASAEQLEKILSYIDIGKQEGAELLIGGERNFLEGE LSNGYYVKPTVFKGHNKMRIFQEEIFGPVVSVTTFKDQEEAL AIANESLYGLGAGVWTRDINTAYRVGRGIQAGRVWTNCYHAY PAHAAFGGYKLSGIGRETHKMMLEHYQQTKNLLVSYSPKKLG FF* |
| A0A7W0BY09 | ALDH-109 | MIYAQPGQPGSLVTFKKRYENFIGGKWVPPVDGEYFENISPV TGQVYCEVPRSKAADIELALDAAHAAKDAWGRTSAAERARIL NKIADRMEENLEMLAVAETWENGKPVRETLAADIPLAIDHFR YFAGCIRAEEGTLAEIDHDTVAYHFKEPLGVVGQIIPWNFPI LMAAWKLAPALAAGNCVVLKPAEQTPTSILVLIELIQDLVPP GVINIVNGFGLEAGKPLASNSRISKIAFTGETTTGRLIMQYA SQNIIPVTLELGGKSPNIFFEDVAEKDDEFFDKALEGFTMFA LNQGEVCTCPSRALIQESIYEKFIERALERVKQIKQGNPLDT ETMIGAQASSEQLEKILSYIDIGKQEGAELLIGGERNFLEGD LNGGYYVKPTVFKGHNKMRIFQEEIFGPVVSVTTFKDKDEAL AIANETLYGLGAGVWTRDINTAYRFGRGIQAGRVWTNCYHAY PAHAAFGGYKLSGIGRETHKMMLEHYQQTKNLLVSYSPKKLG FF* |
| A0A7W8JJ24 | ALDH-110 | MIYAQPGQPGSLITFKKRYENFIGGEWVPPIDGEYFENVSPV TGQVYCEVPRSKAADIELALDAAHAAKETWGRTSVAERARLL NKIADRMEENLDMLAVAETWENGKPIRETRAADIPLAIDHFR YFAGCIRAEEGTLAELDDDTVAYHFKEPLGVVGQIIPWNFPI LMAAWKLAPALAAGNCVVLKPAEQTPTSILVLIELIQDLLPK |

| | | |
|---|---|---|
| | | GVVNVVNGFGLEAGKPLASNPRIAKVAFTGETTTGRLIMQYA<br>SQNLIPVTLELGGKSPNIFFADVAAKDDEFFDKALEGFTMFA<br>LNQGEVCTCPSRALIEESIYDVFMERALERVKQIKQGNPLDT<br>TTMIGAQASSEQLEKILSYIDIGKQEGAELLIGGERNMLEGD<br>LQGGYYIKPTVFKGHNRMRIFQEEIFGPVVSVTTFKDKEEAL<br>AIANDTLYGLGAGVWTRDMNTAYRFGRGIQAGRVWTNCYHAY<br>PAHAAFGGYKMSGIGRETHKMMLDHYQQTKNLLVSYSPKKLG<br>FF* |
| A0A7W9YP99 | ALDH-111 | MIYAQPGQPGSLITFKKRYENFIGGEWVPPIDGEYFENISPV<br>TGQVYCEVPRSKAADIELALDAAHAAKEAWGRTSVAERARLL<br>NKIADRMEENLDMLAVAETWENGKPIRETRAADIPLAIDHFR<br>YFAGCIRAEEGTLAELDADTVAYHFKEPLGVVGQIIPWNFPI<br>LMAAWKLAPALAAGNCVVLKPAEQTPTSILVLIELIQDLLPK<br>GVVNVVNGFGLEAGKPLASNPRIAKIAFTGETTTGRLIMQYA<br>SQNLIPVTLELGGKSPNIFFADVAAKDDEFFDKALEGFTMFA<br>LNQGEVCTCPSRALIEESIYDVFMERALERVKQIKQGNPLDT<br>STMIGAQASSEQLEKILSYIDIGKQEGAELLIGGERNMLEGD<br>LQGGYYVKPTVFKGHNRMRIFQEEIFGPVVSVTTFKDQEEAL<br>SIANDTLYGLGAGVWTRDMNTAYRFGRGIQAGRVWTNCYHAY<br>PAHAAFGGYKMSGIGRETHKMMLDHYQQTKNLLVSYSPKKLG<br>FF* |
| A0A7W9YPV4 | ALDH-112 | MIYAQPGQPGSLITFKKRYENFIGGEWVPPIDGEYFENISPV<br>TGQVYCEVPRSKAVDIELALDAAHAAKEAWGRTSVAERARLL<br>NKIADRMEENLDMLAVAETWENGKPIRETRAADIPLAIDHFR<br>YFAGCIRAEEGTLAELDNDTVAYHFKEPLGVVGQIIPWNFPI<br>LMAAWKLAPALAAGNCVVLKPAEQTPTSILVLIELIQDLLPK<br>GVVNVVNGFGLEAGKPLASNPRIAKVAFTGETTTGRLIMQYA<br>SQNLIPVTLELGGKSPNIFFADVAAKDDEFFDKALEGFTMFA<br>LNQGEVCTCPSRALIEESIYDVFMERALERVKQIKQGNPLDT<br>STMIGAQASSEQLEKILSYIDIGKQEGAELLIGGERNMLEGD<br>LQGGYYVKPTVFKGHNRMRIFQEEIFGPVVSVTTFKDQEEAL<br>SIANDTLYGLGAGVWTRDMNTAYRFGRGIQAGRVWTNCYHAY<br>PAHAAFGGYKMSGIGRETHKMMLEHYQQTKNLLVSYSPKKLG<br>FF* |
| A0A840DPH9 | ALDH-113 | MIYAQPGQPGSLITFKKRYENFIGGNWVPPVDGEYFENISPV<br>TGLPYCEVPRSKAADIELALDAAHEAKEAWGRTSVAERARIL<br>NKIADRMEENLELLAVVETWENGKPIRETLAADIPLAIDHFR<br>YFAGCIRAQEGSLAELDYDTVAYHFKEPLGVVGQIIPWNFPI<br>LMAAWKLAPALAAGNCVVLKPAEQTPTSILVLIELIEDLLPK<br>GVVNVVNGFGLEAGKPLASNPRIAKVAFTGETTTGRLIMQYA<br>SQNIIPVTLELGGKSPNIFFPDVAEKDDEFFDKALEGFTMFA<br>LNQGEVCTCPSRALIHESIYDVFMERALERVKQIKQGNPLDT<br>ETMIGAQASSEQLEKILSYIDIGKQEGAELLIGGERNFLEGE<br>LSGGYYVQPTVFKGHNKMRIFQEEIFGPVVSVTTFKDEDEAL<br>AIANETLYGLGAGVWSRNINTAYRFGRGIQAGRVWTNCYHAY<br>PAHAAFGGYKMSGIGRETHKMMLEHYQQTKNLLVSYSPKKLG<br>FF* |
| UPI001363667A | ALDH-114 | MIYAQPDQPGALVTFKKRYENFIGGKWVPPVDGEYFENISPV<br>TGQAYCEVPRSRAADIELALDAAHAAKDAWGRTSPAERARIL<br>NKIADRMEENLEMLAVAETWENGKPIRETLNADIPLAIDHFR<br>YFAGCIRAEEGTLSEIDHDTVAYHFKEPLGVVGQIIPWNFPL |

| | | |
|---|---|---|
| | | LMAAWKLAPALAAGNCVVLKPAEQTPTSILVLMELIQDILPP GVVNVVNGFGVEAGKPLASSSRIAKIAFTGETTTGRLIMQYA SQNIIPVTLELGGKSPNIFFEDVAAKDDEFFDKAIEGFTLFA LNQGEICTCPSRALIHESIYDKFMERALERVKQIKQGNPLDT ETMIGAQASSEQLEKILSYIDIGKQEGAELLAGGERNILEGD LKDGYYVKPTVFKGHNKMRIFQEEIFGPVVAVTTFKDMDEAL EIANDTLYGLGAGVWTRDINTAYRVGRGIQAGRVWTNCYHMY PAHAAFGGYKQSGFGRETHKMMLEHYQQTKNLLVSYSPKKLG LF* |
| C5D8G6 | ALDH-115 | MIYAQPGQPGSLITFKKRYENFIGGKWVPPVDGEYFENISPV TGKPYCEVPRSKAADIELALDAAHAAKDAWGRTSPAERARIL NKIADRMEENLEMLAVAETWENGKPIRETLNADIPLAIDHFR YFAGCIRAEEGTLSEIDHDTVAYHFKEPLGVVGQIIPWNFPL LMAAWKLAPALAAGNCVVLKPAEQTPTSILVLMELIQDILPP GVVNVVNGFGVEAGKPLASSSRIAKIAFTGETTTGRLIMQYA SQNIIPVTLELGGKSPNIFFEDVAAKDDEFFDKAIEGFTLFA LNQGEICTCPSRALIHESIYDKFMERALERVKQIKQGNPLDT ETMIGAQASSEQLEKILSYIDIGKQEGAELLAGGERNILEGD LKDGYYVKPTVFKGHNKMRIFQEEIFGPVVAVTTFKDMDEAL EIANDTLYGLGAGVWTRDINTAYRVGRGIQAGRVWTNCYHMY PAHAAFGGYKQSGFGRETHKMMLEHYQQTKNLLVSYSPKKLG LF* |
| UPI0002BF8D7A | ALDH-116 | MIYAQPGQPGSLITFKKRYENFIGGEWVPPIDGEYFENISPV TGQVYCEVPRSKAADIELALDAAHAAKETWGRTSVAERARLL NKIADRMEENLDMLAVAETWENGKPIRETRAADIPLAIDHFR YFAGCIRAEEGTLAELDDDTVAYHFKEPLGVVGQIIPWNFPI LMAAWKLAPALAAGNCVVLKPAEQTPTSILVLIELIQDLLPK GVINVVNGFGLEAGKPLASNPRIAKVAFTGETTTGRLIMQYA SQNLIPVTLELGGKSPNIFFADVVAKDDEFFDKALEGFTMFA LNQGEVCTCPSRALIEESIYDVFMERALERVKQIKQGNPLDT STMIGAQASSEQLEKILSYIDIGKQEGAELLIGGERNMLEGD LQGGYYVKPTVFKGHNRMRIFQEEIFGPVVSVTTFKDKDEAL AIANDTLYGLGAGVWTRDINTAYRFGRGIQAGRVWTNCYHAY PAHAAFGGYKMSGIGRETHKMMLEHYQQTKNLLVSYSPKKLG FF* |
| M8CWA2 | ALDH-117 | MIYAQPGQPGSLITFKKRYENFIGGEWVPPIDGEYFENISPV TGQVYCEVPRSKAADIELALDAAHAAKEAWGRTSVAERARLL NKIADRMEENLDMLAVAETWENGKPIRETRAADIPLAIDHFR YFAGCIRAEEGTLAELDADTVAYHFKEPLGVVGQIIPWNFPI LMAAWKLAPALAAGNCVVLKPAEQTPTSILVLIELIQDLLPK GVVNVVNGFGLEAGKPLASNPRIAKVAFTGETTTGRLIMQYA SQNLIPVTLELGGKSPNIFFADVATKDDEFFDKALEGFTMFA LNQGEVCTCPSRALIEESIYDVFMERALERVKQIKQGNPLDT STMIGAQASSEQLEKILSYIDIGKQEGAELLIGGERNMLEGD LQGGYYVKPTVFKGHNRMRIFQEEIFGPVVSVTTFKNQDEAL SIANDTLYGLGAGIWTRDMNTAYRFGRGIQAGRVWTNCYHAY PAHAAFGGYKMSGIGRETHKMMLEHYQQTKNLLVSYSPKKLG FF* |
| S5ZGT3 | ALDH-118 | MIYAQPGQPGALVTFKTRYENFIGGKWVPPVDGEYFENITPI TGKPYCEVPRSKAADIELALDAAHAAKEAWGRTSPAERARLL NKIADRMEENLEMLAVAETWENGKPIRETLAADIPLAIDHFR |

| | | |
|---|---|---|
| | | YFASCIRAQEGAISEIDHNTVAYHFKEPLGVVGQIIPWNFPI LMAAWKLAPALAAGNCVVLKPAEQTPTSILVLIELIEDLLPP GVVNIVNGFGLEAGKPLASNPRVAKVAFTGETTTGRLIMQYA SQNIVPVTLELGGKSPNIFFADVMDKDDEFLDKALEGFTMFA LNQGEVCTCPSRALIHESIYDAFMERALERVKQIKQGNPLDT ETMIGAQASSEQLEKILSYIDIGKQEGAELLIGGERNMLEGE LSGGYYVKPTIFKGHNKMRIFQEEIFGPVLAVTTFKDQDEAL SIANETLYGLGAGVWTRDMNTAYRFGRGIQAGRVWTNCYHVY PAHAAFGGYKMSGIGRETHKMMLDHYQQTKNLLVSYSPKKLG LF* |
| S7SR93 | ALDH-119 | MEEESGMIYAQPGQPGALVTFKKRYENFIGGKWVPPVDGEYF ENITPITGQPYCEVPRSKAADIELALDAAHAAKDAWGRTSPA ERARLLNKIADRMEEHLEMLAVAETWENGKPIRETLAADIPL AIDHFRYFASCIRAQEGTISEIDHDTVAYHFKEPLGVVGQII PWNFPILMAAWKLAPALAAGNCVVLKPAEQTPTSILVLIELI EDLLPPGVVNIVNGFGLEAGKPLASNPRVAKVAFTGETTTGR LIMQYASQNIVPVTLELGGKSPNIFFADVMDKDDDFLDKALE GFTMFALNQGEVCTCPSRALIHESIYDAFMERALERVKQIKQ GNPLDTETMIGAQASSEQLEKILSYIDIGKQEGAELLIGGER NMLEGELSGGYYVKPTVFKGHNKMRIFQEEIFGPVLAVTTFK DHDEALSIANETLYGLGAGVWTRDINTAYRFGRGIQAGRVWT NCYHVYPAHAAFGGYKMSGIGRETHKMMLDHYQQTKNLLVSY SPKKLGLF* |
| U2WRT1 | ALDH-120 | MEEESGMIYAQPGQPGALVTFKKRYENFIGGKWVPPVDGEYF ENITPITGQPYCEVPRSKAADIELALDAAHAAKDAWGRTSPA ERARLLNKIADRMEENLEMLAVAETWENGKPIRETLAADIPL AIDHFRYFASCIRAQEGTISEIDHDTVAYHFKEPLGVVGQII PWNFPILMAAWKLAPALAAGNCVVLKPAEQTPTSILVLIELI EDLLPPGVVNIVNGFGLEAGKPLASNPRVAKVAFTGETTTGR LIMQYASQNIVPVTLELGGKSPNIFFADVMDKDDEFLDKALE GFTMFALNQGEVCTCPSRALIHESIYDAFMERALERVKQIKQ GNPLDTETMIGAQASSEQLEKILSYIDIGKQEGAELLIGGER NMLEGELAGGYYVKPTIFKGHNKMRIFQEEIFGPVLAVTTFK DHDEALSIANETLYGLGAGVWTRDINTAYRFGRGIQAGRVWT NCYHVYPAHAAFGGYKMSGIGRETHKMMLDHYQQTKNLLVSY SPKKLGLF* |
| UPI0004DFAE51 | ALDH-121 | MIYAQPGQPGALVTFKKRYENFIGGKWVPPVDGEYFENITPI TGQPYCEVPRSKAADIELALDAAHAAKDAWGRTSPAKRARLL NQIADRMEENLEMLAVAETWENGKPIRETLAADIPLAIDHFR YFASCIRAQEGTISEIDHDTVAYHFKEPLGVVGQIIPWNFPI LMAAWKLAPALAAGNCVVLKPAEQTPTSILVLIELIQDLLPP GVVNVVNGFGLEAGKPLASNPRVAKVAFTGETTTGRLIMQYA SQNIVPVTLELGGKSPNIFFADVMDKDDEFLDKALEGFTMFA LNQGEVCTCPSRALIHESIYDAFMERALERVKQIKQGNPLDT ETMIGAQASSEQLEKILSYIDIGKQEGAELLIGGERNMLEGE LAGGYYVKPTIFKGHNKMRIFQEEIFGPVLAVTTFKDNDEAL AIANETLYGLGAGVWTRDINTAYRFGRGIQAGRVWTNCYHVY PAHAAFGGYKMSGIGRETHKMMLDHYQQTKNLLVSYSPKKLG LF* |
| UPI001EEBB712 | ALDH-122 | MIYAQPGQLGSLITFKKRYENFIGGEWVSPIDGEYFENISPV TGQVYCEVPRSKAADIELALDAAHAAKEAWGRTSVAERARLL |

| | | |
|---|---|---|
| | | NKIADRMEENLDMLAVAETWENGKPIRETRAADIPLAIDHFR YFAGCIRAEEGTLAELDNDTVSYHFKEPLGVVGQIIPWNFPI LMAAWKLAPALAAGNCVVLKPAEQTPTSILVLIELIQDLLPK GVVNVVNGFGLEAGKPLASNPRIAKVAFTGETTTGRLIMQYA SQNLIPVTLELGGKSPNIFFADVAAKDDEFFDKALEGFTMFA LNQGEVCTCPSRALIEESIYDIFMERALERVKQIKQGNPLDT TTMIGAQASSEQLEKILSYIDIGKREGAELLIGGERNMLEGD LQGGYYVKPTVFKGHNRMRIFQEEIFGPVVSVTTFKDKDEAL AIANDTLYGLGAGVWTRDVNTAYRFGRGIQAGRVWTNCYHAY PAHAAFGGYKMSGIGRETHKMMLEHYQQTKNLLVSYSPKKLG FF* |
| UPI0015843A2B | ALDH-123 | MIYAQPGQPGSLVTFKKRYENFIGGKWVPPVDGEYFENISPV TGQAYCEVPRSKAADIELALDAAHAAKDAWGRTSPAERARIL NKIADRMEENLEMLAVAETWENGKPIRETLNADIPLAIDHFR YFAGCIRAEEGTLAEIDHNTVAYHFKEPLGVVGQIIPWNFPL LMAAWKLAPALAAGNCVVLKPAEQTPTSILVLMELIQDLLPP GVVNIVNGFGLEAGKPLASNPRVAKVAFTGETTTGRLIMQYA SQNIIPVTLELGGKSPNIFFEDVAEKDDEFFDKAIEGFTLFA LNQGEVCTCPSRALIQESIYDKFMERALERVKQIKQGNPLDT ETMIGAQASSEQLEKILSYIDIGKQEGAELLIGGERNYLEGD LRNGYYVKPTVFKGHNKMRIFQEEIFGPVVAVTTFKDKDEAL AIANDTLYGLGAGVWTRDINTAYRFGRGIQAGRVWTNCYHIY PAHAAFGGYKMSGFGRETHKMMLEHYQQTKNLLVSYSPKKLG LF* |
| UPI0014923DAD | ALDH-124 | MIYAQPGQPGSLITFKKRYENFIGGEWVPPIDGEYFENISPV TGQVYCEVPRSKAADIELALDAAHAAKEAWGRTSVAERARLL NKIADRMEENLDMLAVAETWENGKPIRETRAADIPLAIDHFR YFAGCIRAEEGTLAEIDADTVAYHFKEPLGVVGQIIPWNFPI LMAAWKLAPALAAGNCVVLKPAEQTPTSILVLIELIQDLLPK GVVNVVNGFGLEAGKPLASNPRIAKVAFTGETTTGRLIMQYA SQNLIPVTLELGGKSPNIFFADVAAKDDEFFDKALEGFTMFA LNQGEVCTCPSRALIEESIYDIFMERALERVKQIKQGNPLDT TTMIGAQASSEQLEKILSYIDIGKQEGAELLIGGERNMLEGD LQGGYYVKPTVFKGHNRMRIFQEEIFGPVVSVTTFKDKDEAL AIANDTLYGLGAGVWTRDVNTAYRFGRGIQAGRVWTNCYHAY PAHAAFGGYKMSGIGRETHKMMLEHYQQTKNLLVSYSPKKLG FF* |
| UPI00030EE752 | ALDH-125 | MIFAQPGQPGSLITFKKRYENFIGGEWVPPIDGEYFENISPV TGQVYCEVPRSKAADIELALDAAHAAKESWGRTSVAERARLL NKIADRMEEHLDMLAVAETWENGKPIRETRAADIPLAIDHFR YFAGCIRAEEGTLAELDNDTVSYHFKEPLGVVGQIIPWNFPI LMAAWKLAPALAAGNCVVLKPAEQTPTSILVLIELIQDLLPK GVVNVVNGFGLEAGKPLASNPRIAKVAFTGETTTGRLIMQYA SQNLIPVTLELGGKSPNIFFADVATKDDEFFDKALEGFTMFA LNQGEVCTCPSRALIEESIYDLFMERALERVKQIKQGNPLDT TTMVGAQASSEQLEKILSYIDIGKQEGAELLIGGERNMLEGD LQGGYYVKPTVFKGHNRMRIFQEEIFGPVVSVTTFKDKDEAL SIANDTLYGLGAGVWTRDVNTAYRFGRGIQAGRVWTNCYHAY PAHAAFGGYKMSGIGRETHKMMLEHYQQTKNLLVSYSPKKLG FF* |

| | | |
|---|---|---|
| UPI0005CD4570 | ALDH-126 | MIYAQPGQPGSLVTFKKRYENFIGGKWVPPVDGEYFENISPV TGKVYCEVPRSKAADIELALDAAHAAKEAWGRTSAAERAKIL NKIADRMEENREMLAVVETWENGKPIRETLAADIPLAIDHFR YFAGCIRAEEGSLAELDHDTVAYHFKEPLGVVGQIIPWNFPI LMAAWKLAPALAAGNCVVLKPAEQTPTSILVLMELIEDLLPP GVVNIVNGFGLEAGKPLASSSRIAKVAFTGETTTGRLIMQYA SQNIIPVTLELGGKSPNIFFEDVMEKDDEFFDKALEGFTMFA LNQGEVCTCPSRALIQESIYDAFMERALERVKQIKQGNPLDT ETMIGAQASSEQLEKILSYIDIGKQEGAELLIGGERNVLEGD LSGGYYVKPTVFKGHNKMRIFQEEIFGPVVSVTTFKDQEEAL AIANETLYGLGAGVWTRDINTAYRVGRGIQAGRVWTNCYHAY PAHAAFGGYKLSGVGRENHKMMLEHYQQTKNLLVSYSPKKLG FF* |
| UPI00228641DC | ALDH-127 | MIYAQPGQPGSLITFKKRYENFIGGQWVPPVDGEYFENISPV TGQVYCEVPRSKAADIELALDAAHAAKDAWGRTSVAERARIL NKIADRMEENLEMLAVAETWENGKPIRETLAADIPLAIDHFR YFAGCIRAEEGTLAELDHDTVAYHFKEPLGVVGQIIPWNFPI LMAAWKLAPALAAGNCVVLKPAEQTPTSILVLMELIADLLPK GVVNVVNGFGLEAGKPLASSPRIAKVAFTGETTTGRLIMQYA SQNIIPVTLELGGKSPNIFFADVAAKDDEFFDKALEGFTMFA LNQGEVCTCPSRALIEESIYDVFMERALERVKQIKQGNPLDT ETMIGAQASSEQLEKILSYIDIGKQEGAELLIGGERNFLEGE LSGGYYVKPTVFKGHNKMRIFQEEIFGPVVSVTTFKDKDEAL AIANETLYGLGAGVWTRDINTAYRFGRGIQAGRVWTNCYHAY PAHAAFGGYKLSGIGRETHKMMLEHYQQTKNLLVSYSPKKLG FF* |
| UPI00135864F6 | ALDH-128 | MIYAQPGQPGALVTFKKRYENFIGGKWVPPVDGEYFENITPI TGQPYCEVPRSKAADIELALDAAHAAKDAWARTSPAERARLL NKIADRMEENLEMLAVAETWENGKPIRETLAADIPLAIDHFR YFASCIRAQEGTISEIDHDTVAYHFKEPLGVVGQIIPWNFPI LMAAWKLAPALAAGNCVVLKPAEQTPTSILVLIELIEDLLPP GVVNIVNGFGLEAGKPLASNPRVAKVAFTGETTTGRLIMQYA SQNIVPVTLELGGKSPNIFFADVMDKDDEFLDKALEGFTMFA LNQGEVCTCPSRALIHESIYDAFMERALERVKQIKQGNPLDT ETMIGAQASSEQLEKILSYIDIGKQEGAELLIGGERNMLEGE LSGGYYVKPTVFKGHNKMRIFQEEIFGPVLAVTTFQDHDEAL SIANETLYGLGAGVWTRDINTAYRFGRGIQAGRVWTNCYHIY PAHAAFGGYKMSGIGRETHKMMLDHYQQTKNLLVSYSPKKLG LF* |
| UPI00208DAAE2 | ALDH-129 | MIYAQPGQPGSLITFKKRYENFIGGEWVPPIDGEYFENISPV TGQVYCEVPRSKAADIELALDAAHAAKEAWGRTSVAERARLL NKIADRMEENLDMLAVAETWENGKPIRETRAADIPLAIDHFR YFAGCIRAEEGTLAELDNDTVSYHFKEPLGVVGQIIPWNFPI LMAAWKLAPALAAGNCVVLKPAEQTPTSILVLIELIQDLLPK GVVNVVNGFGLEAGKPLASNPRIAKVAFTGETTTGRLIMQYA SQNLIPVTLELGGKSPNIFFADVAAKDDEFFDKALEGFTMFA LNQGEVCTCPSRALIEESIYDIFMERALERVKQIKQGNPLDT TTMIGAQASSEQLEKILSYIDIGKQEGAELLIGGERNMLGGD LQGGYYVKPTVFKGHNRMRIFQEEIFGPVVSVTTFKDKDEAL AIANDTLYGLGAGVWTRDVNTAYRFGRGIQAGRVWTNCYHAY |

| | | |
|---|---|---|
| | | PAHAAFGGYKMSGIGRETHKMMLEHYQQTKNLLVSYSPKKLG FF* |
| UPI001315BED2 | ALDH-130 | MIYTQPGHPGSLITFKKRYENFIGGKWVPPVDGEYFENISPV TGQPYCEVPRSKAADIELALDAAHEAKEAWSRTSVTERARIL NKIADRMEENLEMLAVAETWENGKPVRETLAADIPLAIDHFR YFAGCIRAQEGSLAEIDNDTVAYHFKEPLGVVGQIIPWNFPI LMAAWKLAPALAAGNCVVLKPAEQTPTSILVLIELIEDLLPK GVVNVVNGFGLEAGKPLASNPRIAKVAFTGETTTGRLIMQYA SQNIIPVTLELGGKSPNIFFADVAEQDDEFFDKALEGFTMFA LNQGEVCTCPSRALIQESIYDTFMERALERVKQIKQGNPLDT ETMIGAQASSEQLEKILSYIDIGKQEGAELLIGGERNFLEGE LRNGYYVKPTVFKGHNKMRIFQEEIFGPVVSVTTFKDKEEAL AIANETLYGLGAGVWTRDVNTAYRFGRGIQAGRVWTNCYHAY PAHAAFGGYKMSGIGRETHKMMLEHYQQTKNLLVSYSPQKLG FF* |
| UPI000489A7CA | ALDH-131 | MIYAQPGQPGSLITFKKRYENFIGGKWVPPVDGEYFENISPV TGQAYCEVPRSKAADIELALDAAHEAKEAWGRTSPAERARIL NKIADRMEENLEMLAVAETWENGKPIRETLAADIPLAIDHFR YFAGCIRAQEGSLAELDDDTVAYHFKEPLGVVGQIIPWNFPI LMAAWKLAPALAAGNCVVLKPAEQTPTSILVLIELIEDLLPK GVINVVNGFGLEAGKPLASNPRVAKVAFTGETTTGRLIMQYA SQNIIPVTLELGGKSPNIFFEDVMEKDDEFFDKALEGFTMFA LNQGEVCTCPSRALIQESIYDAFIERALERVKQIKQGNPLDT TTMIGAQASSEQLEKILSYIDIGKQEGAELLIGGERNFLEGE LRSGYYVKPTVFKGHNKMRIFQEEIFGPVVSVTTFKDKDEAL AIANETLYGLGAGVWTRDINTAYRFGRGIQAGRVWTNCYHAY PAHAAFGGYKMSGIGRETHKMMLEHYQQTKNLLISYSPQKLG FF* |
| UPI001FCBF132 | ALDH-132 | MIYAQPGQPGSLITFKKRYENFIGGKWVPPVDGEYFENISPV TGKPYCEVPRSKAADIELALDAAHAAKDAWGRTSPAERARIL NKIADRMEENLEMLAVAETWENGKPIRETLNADIPLAIDHFR YFAGCIRAEEGTLAEIDHDTVAYHFKEPLGVVGQIIPWNFPL LMAAWKLAPALAAGNCVVLKPAEQTPTSILVLMELIEDLLPP GVVNIVNGFGLEAGKPLASSNRIAKIAFTGETTTGRLIMQYA SQNIIPVTLELGGKSPNIFFEDVAAKDDEFFDKALEGFTLFA LNQGEVCTCPSRALIQESIYDKFMERALERVKQIKQGNPLDT ETMIGAQASSEQLEKILSYIDIGKQEGAELLIGGERNYLEGD LRDGYYVKPTVFKGHNKMRIFQEEIFGPVVSVTTFKDFDEAL EIANETLYGLGAGVWTRDINTAYRVGRGIQAGRVWTNCYHVY PAHAAFGGYKLSGFGRETHKMMLEHYQQTKNLLVSYSPKKLG LF* |
| UPI0001D589C3 | ALDH-133 | MIYAQPGQPGALVTFKKRYENFIGGKWVPPVDGEYFENITPI TGQPYCEVPRSKAADIELALDAAHAAKDAWGRTSPAERARLL NKIADRMEENLEMLAVAETWENGKPIRETLAADIPLAIDHFR YFASCIRAQEGTISEIDHDTVAYHFKEPLGVVGQIIPWNFPI LMAAWKLAPALAAGNCVVLKPAEQTPTSILVLIKLIEDLLPP GVVNIVNGFGLEAGKPLASNPRVAKVAFTGETTTGRLIMQYA SQNIVPVTLELGGKSPNIFFADVMDKDDEFLDKALEGFTMFA LNQGEVCTCPSRALIHESIYDAFMERALERVKQIKQGNPLDT ETMIGAQASSEQLEKILSYIDIGKQEGAELLIGGERNMLEGE LAGGYYVKPTIFKGHNKMRIFQEEIFGPVLAVTTFKDHDEAL |

| | | |
|---|---|---|
| | | SIANETLYGLGAGVWTRDINTAYRFGRGIQAGRVWTNCYHVY PAHAAFGGYKMSGIGRETHKMMLEHYQQTKNLLVSYSPKKLG LF* |
| M2XRT2 | ALDH-134 | MNGSPKSYPRPGESGCPVTFKSQYENFIGGKWVPPVKGQYFD NVSPVNGKVFCKSARSTAEDIELALDAAHAAKDKFGATTFEQ RAKLLNQMADAIEQNLEKLALAEVWDNGKPIREALAADIPLA ADHLRYFASAVRTQEGFIAEHKENTVAYHFHEPLGVVGAIIP WNFPILMAIWKLSPALAGGNCIIIKPAEQTPASIMVLMEVWA NIVPPGVINVVTGYGPETGKPLACSPRIAKIAFTGETTTGQL IMQYASQNIIPVTLELGGKSPNIFFADIANENDEFFEKAVEG CVMFVLNQGEVCTCPSRALVHESIYDKFIEKVVQRLGKIKQG DPLNMETMVGAQVSTEQMDKILHYVELGKKEGAQCIVGGSGK KAIGGELEGGYYIEPTIFKGDNKMRIFQEEIFGPVLSVTTFR TEEEAIQIANDTSYGLGAGLWTRDIMKAYRVSRAIKAGRVWV NCYHEYPSHAAFGGYKKSGFGRECHKLTLEHYQQLKNIIISY SPQPTGLF* |

‡ALDH-044 was isolated from *E. coli* BL21 genomic DNA, and this specific sequence does not have a readily searchable accession number. The amino acid sequence given in the right column is the sequence used in this work. †To clone a complete ALDH-054 from fragment sequences, A0A7L1RJ35 (residues 1-112) was concatenated to A0A7L1RY59 (residues 1-339).

**Supplementary Table 7:** Amino Acid Sequence of TcADH

| Enzyme Name | Plasmid | Amino Acid Sequence |
|---|---|---|
| TcADH | pSAP179 | MAGDGAREQQTSYEHFLTQRKNSDSELDVIKCKAAVLWEVKKPFSIEE VEVAPPKAHEVRIKMVATGICRSDDHVVSGNLAVPFPVILGHEAAGIV ESIGEGVTSVKPGDKVIPLFTPQCGKCRICKHPESNYCLKNDLDKPRG TLQDGTTRFTCRGKSIHHFLSTSTFSQYTVVDEISVVKIDDASPLEKV CLIGCGFSTGYGSAVKVAKVTRGSTCAVFGLGGVGLSVIMGCKAAGAA RIIGVDINKDKFAKAKEVGATECINPQDYKKPIQEVLREMSDGGVDFS FEVIGRLDTMMAALLCCENSCGVSVIVGVPPGSQTLSVDPMLLLTGRT WKGAIFGGFKSKDSVPKLVADFMAKKFSLDPLITHVLPFEKINEGFDL LRSGKSIRTVLTF* |

### B. Supplementary References

1. Li, X. *et al.* Characterization of a broad-range aldehyde dehydrogenase involved in alkane degradation in Geobacillus thermodenitrificans NG80-2. *Microbiol. Res.* **165**, 706–712 (2010).

2. Chenault, H. K. & Whitesides, G. M. Regeneration of nicotinamide cofactors for use in organic synthesis. *Appl. Biochem. Biotechnol.* **14**, 147–197 (1987).

3. Li, H. & Liao, J. C. Engineering a cyanobacterium as the catalyst for the photosynthetic conversion of CO2 to 1,2-propanediol. *Microb. Cell Factories* **12**, 4 (2013).

4. Zhang, L. *et al.* Directed evolution of phosphite dehydrogenase to cycle noncanonical redox cofactors via universal growth selection platform. *Nat. Commun.* **13**, 5021 (2022).
